# Microglial cannabinoid receptor type 1 mediates social memory deficits produced by adolescent THC exposure and 16p11.2 duplication

**DOI:** 10.1101/2023.07.24.550212

**Authors:** Yuto Hasegawa, Juhyun Kim, Gianluca Ursini, Yan Jouroukhin, Xiaolei Zhu, Yu Miyahara, Feiyi Xiong, Samskruthi Madireddy, Mizuho Obayashi, Beat Lutz, Akira Sawa, Solange P. Brown, Mikhail V. Pletnikov, Atsushi Kamiya

**Author notes:** Correspondence (A.K.) and (M.V.P.).

## Abstract

Adolescent cannabis use increases the risk for cognitive impairments and psychiatric disorders. Cannabinoid receptor type 1 (Cnr1) is expressed not only in neurons and astrocytes, but also in microglia, which shape synaptic connections during adolescence. Nonetheless, until now, the role of microglia in mediating the adverse cognitive effects of delta-9-tetrahydrocannabinol (THC), the principal psychoactive constituent of cannabis, has been unexplored. Here, we report that adolescent THC exposure produces microglial apoptosis in the medial prefrontal cortex (mPFC), which was exacerbated in the mouse model of 16p11.2 duplication, a representative copy number variation (CNV) risk factor for psychiatric disorders. These effects are mediated by microglial Cnr1, leading to reduction in the excitability of mPFC pyramidal-tract neurons and deficits in social memory in adulthood. Our findings highlight the importance of microglial Cnr1 to produce the adverse effect of cannabis exposure in genetically vulnerable individuals.

## Introduction

Adolescent cannabis use is associated with an increased risk for psychiatric disorders and cognitive abnormalities^1–3^. The adverse effects of cannabis are mainly mediated by delta-9-tetrahydrocannabinol (THC), the principal psychoactive constituent of cannabis^4^. The effect of THC on brain function is known to result from its binding to the cannabinoid receptor type 1 (Cnr1) on presynaptic terminals, thereby modulating cognitive function^1, 5–7^. However, recent studies have highlighted the importance of Cnr1 expressed on astrocytes in mediating THC-induced cognitive deficits^8–11^. Interestingly, Cnr1 is also expressed in microglia, which play critical roles in synaptic pruning during brain maturation^12, 13^ and in the control of social and cognitive function^14, 15^. The pathological implication of microglia in various psychiatric disorders of neurodevelopmental origin has gained significant attention in recent years^16^. Nevertheless, the role of microglia in mediating the adverse cognitive effects of THC exposure remains unexplored.

The contribution of cannabis use to risk for psychiatric disorders appears to be modulated by genetic vulnerability to psychosis in the context of gene-environment interaction (GxE)^1^. Thus, an unmet need exists in determining the convergent mechanisms whereby adolescent cannabis exposure interacts with genetic susceptibility to psychiatric disorders, ultimately producing adult psychopathology and cognitive impairments. The ∼600-kb duplication (breakpoint 4[BP4]-BP5) on 16p11.2 (16p11dup) is a copy number variation (CNV) that reproducibly increases the risk for a range of cognitive defects present in psychiatric disorders^17^. Preclinical studies have shown that the 16p11dup mouse model exhibits behavioral abnormalities in cognitive domains, as well as abnormalities in the dendritic structure of pyramidal neurons and in the GABAergic synapses of the prefrontal cortex (PFC), an area critically involved in social and cognitive functions^18–20^. Nonetheless, potential GxE in the 16p11dup model has yet to be investigated.

Here we sought to identify the role of microglia for the adverse effects of THC exposure on adolescent brain maturation and cognitive functions. We explored whether and how adolescent THC exposure affects microglial function via cannabinoid receptors and whether 16p11dup exacerbates these effects, leading to alterations in PFC function and impairments in social and cognitive function in adulthood.

## Results

### Cnr1 is expressed in the microglia of mouse brains

We first explored cell type-specific expression of Cnr1 in the mouse brain. CD45^+^CD11b^+^TMEM119^+^ microglia, ACSA-2^+^ astrocytes, and NeuN^+^ neurons from the cerebral cortex of wild type mice were isolated by fluorescence activated cell sorting (FACS) at postnatal day 90 (P90) (**Fig. 1a**). We confirmed that Cnr1 mRNA is expressed in microglia, though to a lesser extent than in neurons and astrocytes (**Fig. 1b**). Using the same approach, we also found that microglial Cnr1 mRNA expression was suppressed in *Cnr1^flox/flox^* mice crossed with the *Cx3cr1^CreER^*line (*Cx3cr1^CreER/+^*;*Cnr1^flox/flox^* mice which were given tamoxifen orally once a day for 5 consecutive days), as compared to littermate controls. There were no such changes in Cnr1 expression in neurons or astrocytes between these groups (**Fig. 1c**). In order to confirm Cnr1 expression at the protein level, we used magnetic activated cell sorting (MACS) to isolate microglia-enriched CD11b^+^ cells, ACSA-2^+^ astrocytes, and remaining cells including neurons from the cerebral cortex of *Cx3cr1^CreER/+^*;*Cnr1^flox/flox^*mice and littermate controls (*Cx3cr1^CreER/+^*;*Cnr1^+/+^*). We used this approach as it produces a higher cell yield than FACS with over 95% of collected CD11b^+^ cells being microglia^21^. Western blot analysis showed that the ∼50 kDa major immunoreactive band, which was previously identified as the Cnr1 protein^22–24^, was detected at lower levels in CD11b^+^ cells than in ACSA-2^+^ astrocytes and remaining cells (**Fig. 1d**). ∼50 kDa protein expression was specifically suppressed in the microglia-enriched CD11b^+^ cells of *Cx3cr1^CreER/+^*;*Cnr1^flox/flox^*mice (**Fig. 1d**). Altogether, these results provide compelling evidence for Cnr1 expression in microglia.

**Fig. 1.**
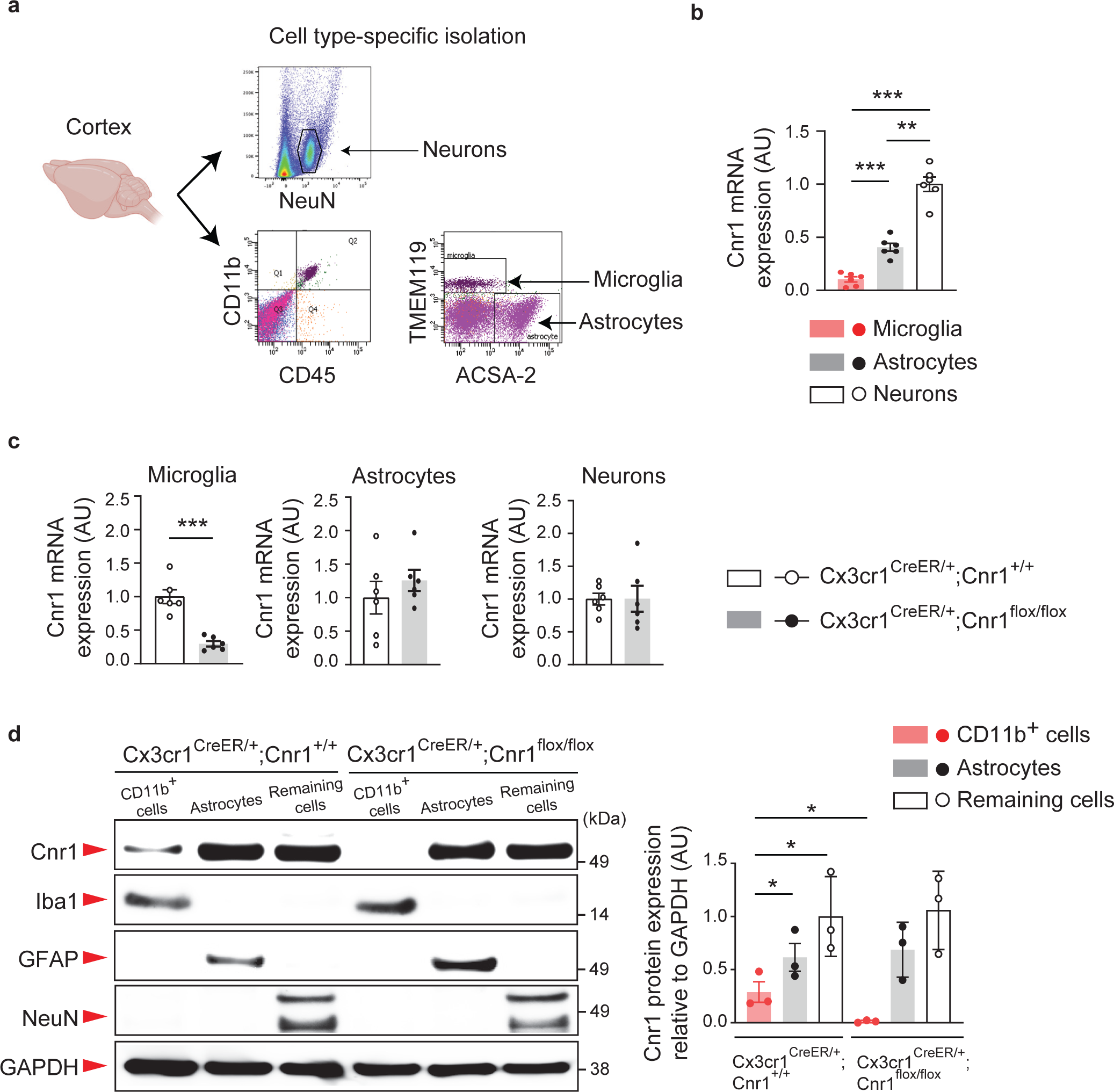
Cnr1 expression in the microglia of the mouse brain. **a,** Experimental flow of Fluorescence-Activated Cell Sorting (FACS)-based microglia (CD45^+^CD11b^+^TMEM119^+^), astrocyte (ACSA-2^+^), and neuron (NeuN^+^) isolation using specific cell markers. **b,** Relative mRNA expression levels of Cnr1 in microglia, astrocytes, and neurons isolated from wild type mice were measured by quantitative real time PCR (qPCR) using the TaqMan assay protocol. *n* = 6 mice per group. \*\*\**p* < 0.001, \*\**p* < 0.01, determined by one-way ANOVA with post hoc Tukey test. **c,** Relative mRNA expression levels of Cnr1 in microglia, astrocytes, and neurons isolated from *Cx3cr1^CreER/+^*;*Cnr1^+/+^*and *Cx3cr1^CreER/+^*;*Cnr1^flox/flox^* mice were measured by qPCR. *n* = 6 mice per group. \*\*\**p* < 0.001, determined by Student *t*-test. **d,** Microglia-enriched CD11b^+^ cells, ACSA-2^+^ astrocytes, and remaining cells including neurons were collected from the cerebral cortex of *Cx3cr1^CreER/+^*;*Cnr1^flox/flox^*mice and littermate controls (*Cx3cr1^CreER/+^*;*Cnr1^+/+^*) and by magnetic activated cell sorting (MACS). For each cell type, expression of Cnr1, marker proteins (Iba1, GFAP, NeuN) and a loading control (GAPDH) in the total protein were analyzed with SDS-PAGE followed by Western blotting with 10 μg of protein sample loaded in each well. *n* = 3 mice per group. \**p* < 0.05, determined by two-way ANOVA with post hoc Tukey test. **(b, c, d)** Data are presented as the mean ± s.e.m.

### Adolescent THC exposure and 16p11dup produce microglial changes in the mPFC

We next explored the effects of adolescent THC treatment and 16p11dup on microglia. Using previously published protocols^25–28^, male and female 16p11dup or wild type littermate control mice were chronically treated with a single daily subcutaneous injection of THC during adolescence P30-P51, corresponding to human adolescence from 12 to 19 years of age^29^ (**Fig. 2a**). At P51, upon completion of adolescent THC treatment in the wild type male mice, we found a reduction of Iba1 mRNA expression in the mPFC, but not other brain regions involved in social and cognitive functions, including ventral and dorsal hippocampus, nucleus accumbens, and amygdala (**Supplementary Fig. 1**). Iba1 expression was also reduced in the mPFC of 16p11dup male mice, but not in other tested brain regions (**Supplementary Fig. 1**). Interestingly, mPFC-specific Iba1 reduction was more severe in 16p11dup mice treated with THC during adolescence than in either wild type mice treated with THC or 16p11dup predisposition alone, suggesting an mPFC-specific GxE effect on microglia (THC × 16p11dup interaction for Iba1 expression [*F*_1,31_ = 4.207, *p* = 0.0488]) (**Supplementary Fig. 1**). We also performed co-staining experiments for Iba1 and P2ry12, which label putative non-activated/homeostatic microglia^30, 31^. We found that adolescent THC treatment and 16p11dup separately reduced the number of both Iba1^+^P2ry12^+^ and Iba1^+^P2ry12^-^ microglia in the mPFC while increasing the percentage of Iba1^+^P2ry12^-^ microglia in the total Iba1^+^ microglia (**Fig. 2b**). These findings were consistent with the observed effects of THC and 16p11dup on mPFC microglial morphology, including a reduction in their cellular processes and an increase in the size of their cell bodies (**Fig. 2c**). It is worth noting that these phenotypes were exacerbated in 16p11dup mice treated with THC during adolescence (THC × 16p11dup interaction for the number of Iba1^+^P2ry12^+^ microglia [*F*_1,20_ = 21.80, *p* = 0.0001] and Iba1^+^P2ry12^-^ microglia [*F*_1,20_ = 5.385, *p* = 0.0310], cellular process area [*F*_1,8_ = 26.62, *p* = 0.0009], and cell body size [*F*_1,8_ = 121.6, *p* < 0.0001]) (**Fig. 2b, c**). Interestingly, we did not observe these microglial changes in young adulthood (P72), following a 3-week abstinence period after the THC treatment (**Supplementary Fig. 2a, b, c**). Although our previous studies reported the impact of THC on astrocyte function in another GxE context^26^, neither adolescent THC treatment nor 16p11dup had an effect on the number of Aldh1l1^+^ astrocytes in these brain regions (**Supplementary Fig. 2d**). Neither adolescent THC treatment nor 16p11dup predisposition affected Iba1 mRNA expression or the total number of Iba1^+^ microglia in the female mice (**Supplementary Fig. 2e, f**).

**Fig. 2.**
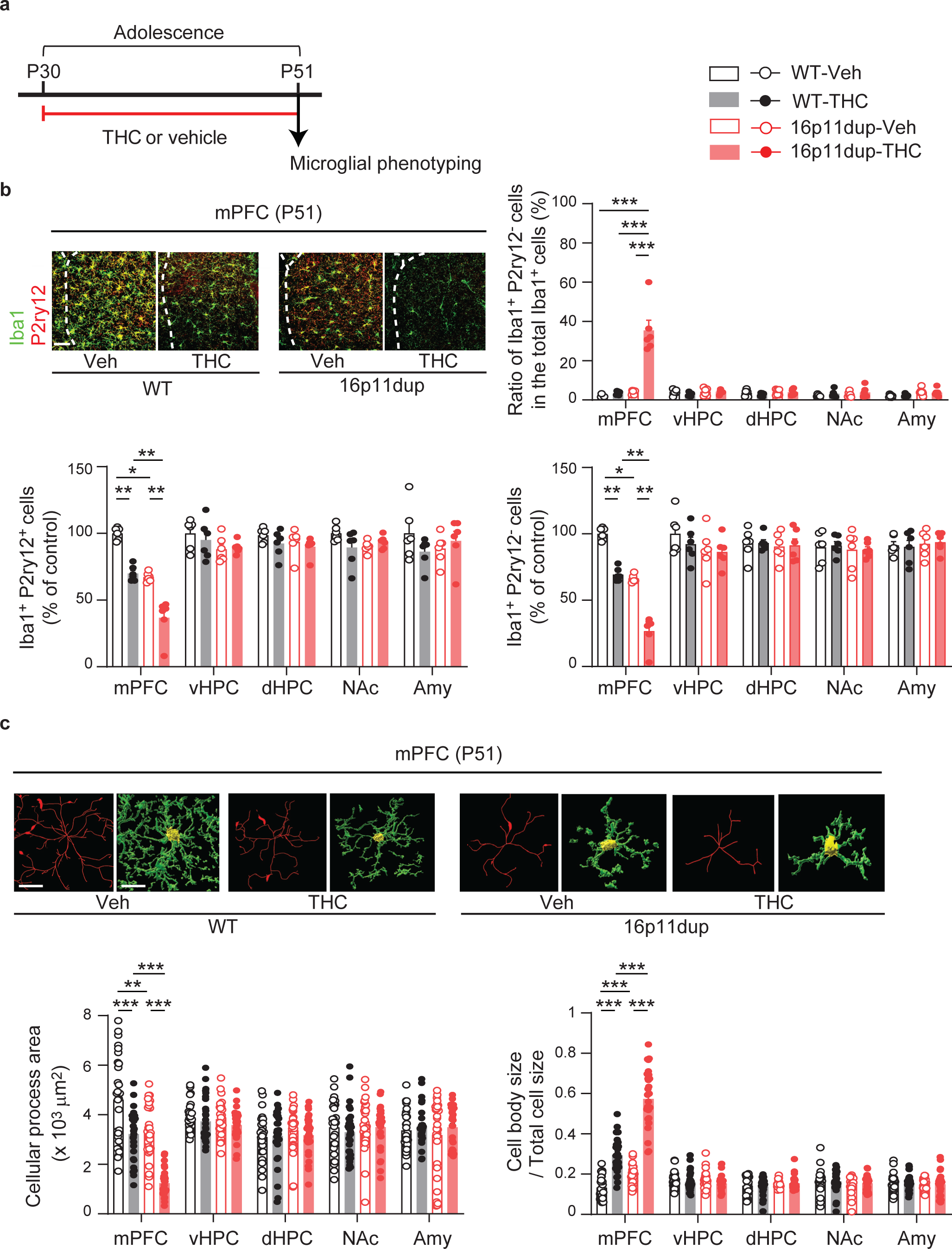
Microglial changes in the mPFC of 16p11dup male mice produced by adolescent THC treatment. **a,** Schematic diagram of the adolescent THC treatment protocol. 16p11dup mice and wild type littermate controls (WT) were treated with THC (s.c., 8mg/kg) or vehicle (Veh) during adolescence (P30-P51), followed by microglial phenotyping at P51 upon completion of THC treatment. **b,** (Top left) Immunohistochemistry of Iba1 (green) and P2ry12 (red) in the medial prefrontal cortex (mPFC) at P51. Scale bar, 50 μm. (Top right) The percentage of Iba1^+^P2ry12^-^ cells among all Iba1^+^ cells in the mPFC, ventral hippocampus (vHPC), dorsal hippocampus (dHPC), nucleus accumbens (NAc), and amygdala (Amy). (Bottom left) The number of Iba1^+^P2ry12^+^ cells and (bottom right) Iba1^+^P2ry12^-^ cells in these brain regions, presented as % of control (*n* = 6 slices in 3 mice per condition). **c,** Microglial morphology analysis of individual Iba1^+^ cells in these brain regions. (Top) Representative tracing images (red) along with images of cellular processes (green) and cell bodies (yellow) of Iba1^+^ cells. Scale bar, 10 μm. (Bottom left) Quantification of cellular process area of Iba1^+^ cells (*n* = 30 cells in 3 mice per condition). (Bottom right) Quantification of the ratio of cell body size to total cell size of Iba1^+^ cells (*n* = 30 cells in 3 mice per condition). (**b, c**) \*\*\**p* < 0.001, \*\**p* < 0.01, \**p* < 0.05, determined by two-way ANOVA with post hoc Tukey test. Data are presented as the mean ± s.e.m.

### Adolescent THC exposure and 16p11dup produce microglial apoptosis in the mPFC

THC induces apoptosis in peripheral immune cells, such as dendritic cells^32, 33^. Therefore, using primary microglia cultures produced from male mice, we next sought to examine whether THC treatment induces microglial apoptosis via cannabinoid receptor signaling and whether these microglial phenotypes are enhanced by 16p11dup predisposition. We observed that THC treatment increased microglial apoptosis and reduced their cellular area and processes (**Fig. 3a, c-f**). In line with the possible role of 16p11dup genes in microglial biology^34–37^, THC-induced microglial apoptosis was synergistically exacerbated by 16p11dup (THC × 16p11dup interaction for microglial apoptosis [*F*_1,44_ = 188.2, *p* < 0.0001]) (**Fig. 3a, e**). THC-induced morphological changes were worsened by 16p11dup (**Fig. 3c, d, f**). In contrast, 22q11 deletion, another major CNV conferring higher risk of psychiatric disorders^38^, did not enhance these microglial phenotypes produced by THC treatment (**Supplementary Fig. 3a, c-f**). Notably, genetic deletion of *Cnr1*, but not *Cnr2* that is primarily expressed in immune cells including microglia^39^, blocked THC-induced microglial apoptosis and morphological changes (**Fig. 3g, i-l**). There was no effect of THC treatment on necrotic cell death of microglia from these mouse models (**Fig. 3b, e, h, k, Supplementary Fig. 3b, e**).

**Fig. 3.**
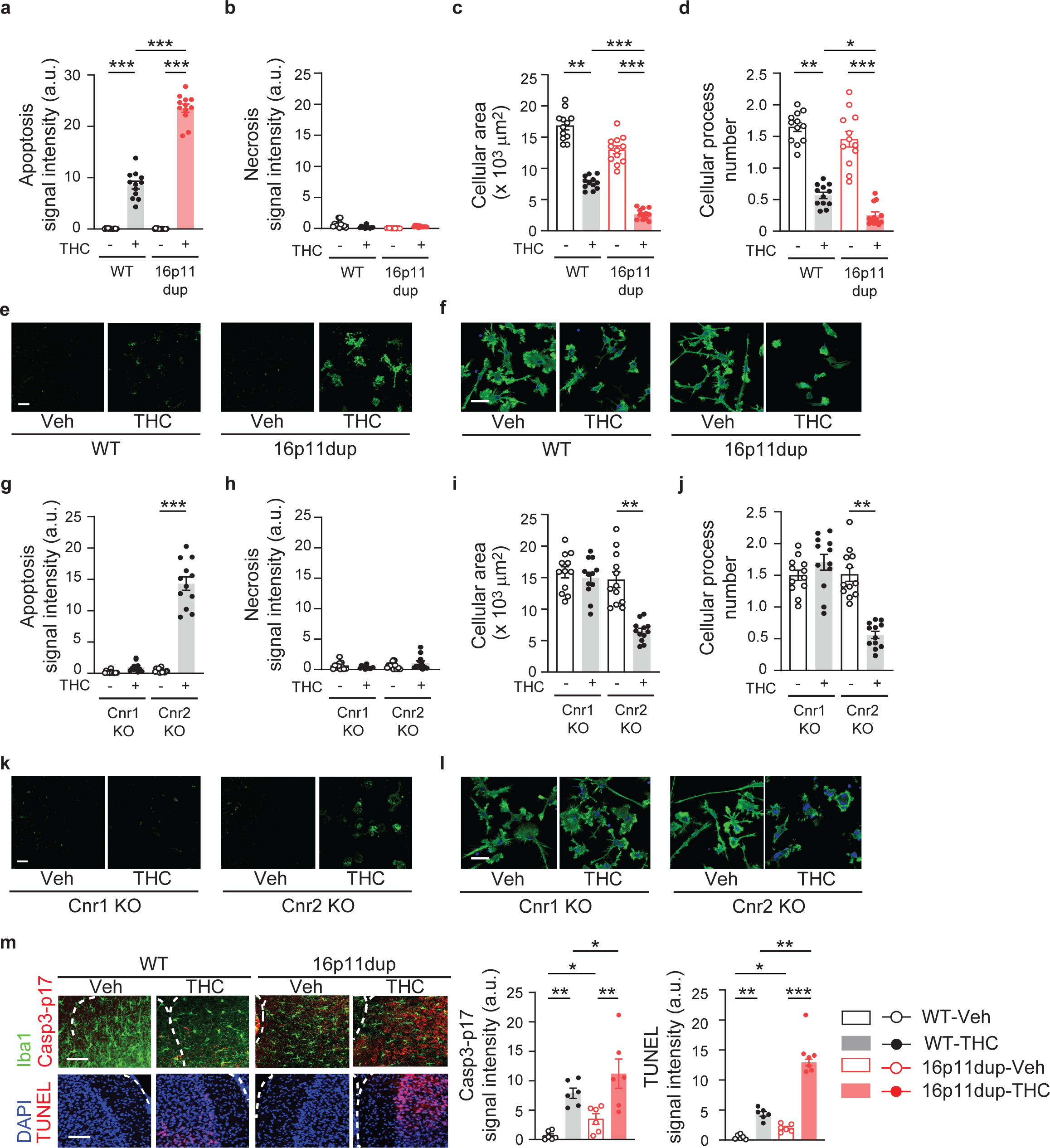
THC-induced Cnr1-mediated microglial apoptosis is exacerbated by 16p11dup. **a**, Apoptosis assay of primary microglia cultures produced from WT and 16p11dup male mice. Quantification of signal intensity of apoptosis marker apopxin (*n* = 12 fields in 3 mice per condition). **b,** Necrosis assay of primary microglia cultures produced from WT and 16p11dup male mice. Quantification of signal intensity of necrosis marker 7-AAD (*n* = 12 fields in 3 mice per condition). **c,** Quantification of cellular process area of phalloidin-stained microglia cultures produced from WT and 16p11dup male mice (*n* = 12 fields in 3 mice per condition). **d**, Quantification of cellular process number of phalloidin-stained microglia cultures produced from WT and 16p11dup male mice (*n* = 12 fields in 3 mice per condition). **e,** Representative images of microglia cell cultures produced from WT and 16p11dup male mice in apoptosis and necrosis assays. Apopxin (green) and 7-AAD (red) are shown. Scale bar, 50 μm. **f,** Immunohistochemistry with antibody against phalloidin (green) of primary microglia cultures produced from WT and 16p11dup male mice. Scale bar, 25 μm. **g**, Apoptosis assay of primary microglia cultures produced from genetic deletion of *Cnr1* (Cnr1 KO) and genetic deletion of *Cnr2* (Cnr2 KO) male mice. Quantification of signal intensity of apoptosis marker apopxin (*n* = 12 fields in 3 mice per condition). **h,** Necrosis assay of primary microglia cultures produced from Cnr1 KO and Cnr2 KO male mice. Quantification of signal intensity of necrosis marker 7-AAD (*n* = 12 fields in 3 mice per condition). **i,** Quantification of cellular process area of phalloidin-stained microglia cultures produced from Cnr1 KO and Cnr2 KO male mice. (*n* = 12 fields in 3 mice per condition). **j**, Quantification of cellular process number of phalloidin-stained microglia cultures of Cnr1 KO and Cnr2 KO male mice (*n* = 12 fields in 3 mice per condition). **k,** Representative images of microglia cell cultures produced from Cnr1 KO and Cnr2 KO male mice in apoptosis and necrosis assays. Apopxin (green) and 7-AAD (red) are shown. Scale bar, 100 μm. **l,** Immunohistochemistry with antibody against phalloidin (green) of primary microglia cultures of Cnr1 KO and Cnr2 KO male mice. Scale bar, 25 μm. **m,** Representative images of immunohistochemistry of Iba1 (green) and Casp3-p17 (red) (top left) as well as TUNEL signals (red) and DAPI (blue) (bottom left) in the mPFC at P51. Scale bar, 50 μm. Quantification of signal intensity of Casp3-p17 and TUNEL (right) (*n* = 6 mice per group). (**a, c, d, m**) \*\*\**p* < 0.001, \*\**p* < 0.01, \**p* < 0.05, determined by two-way ANOVA with post hoc Tukey test. (**g, i, j**) \*\*\**p* < 0.001, \*\**p* < 0.01, determined by Student *t*-test. Data are presented as the mean ± s.e.m.

To obtain *in vivo* evidence of microglial apoptosis in the mPFC, we measured the expression of the active subunit of Caspase3-p17, a marker of the ‘before the point of no return’ in apoptosis with co-labeling for Iba1, and DNA fragmentation (TUNEL), a marker of the ‘after the point of no return’, at P51 upon completion of adolescent THC treatment. Adolescent THC treatment increased the immunoreactivity of Caspase3-p17 in the Iba1^+^ cells and TUNEL signals and these were enhanced in the 16p11dup mice (THC × 16p11dup interaction for Caspase3-p17 immunoreactivity [*F*_1,20_ = 5.023, *p* = 0.0365] and TUNEL signals [*F*_1,20_ = 94.36, *p* < 0.0001]) (**Fig. 3m**). These results suggest that adolescent THC treatment induces *Cnr1*-mediated microglial apoptosis in the mPFC, which was specifically exacerbated by 16p11dup predisposition.

### p53 signaling mediates the convergent effect of adolescent THC exposure and 16p11dup on microglial apoptosis

To gain molecular insights into the THC-induced microglial apoptosis in the GxE context, we performed RNA sequencing (RNA-seq)-based analyses to determine transcriptome changes in the microglia produced by adolescent THC treatment and 16p11dup. Microglia were collected by FACS from the mPFC at P51, followed by RNA-seq analysis using the Smart-seq2 protocol that is applicable for a limited number of FACS-collected cells with full-length coverage. We applied a multicolor flow cytometry analysis using multiple fluorescent markers to collect microglia populations (CD11b^+^/CD45^+^/P2ry12^+^) (**Fig. 4a**). We prioritized the genes that were more likely to be differentially expressed, based on THC treatment alone (n = 569), 16p11dup alone (n = 508), or the genes that were differentially regulated as a GxE (16p11dup with THC; n = 772) (**Fig. 4b, c**). Pathway analysis using the Reactome database revealed gene sets that were significantly altered in the GxE group, including the p53 signaling pathway as well as cell cycle-, mRNA regulation-, cellular response-, and innate immune-related pathways (**Fig. 4d, Supplementary Table 1**). Functional gene interaction networks identified by Ingenuity Pathway Analysis (IPA) demonstrated that many prioritized genes (68 genes in THC vs. WT and 88 genes in WT vs. 16p11dup with THC) have an upstream regulator, *Trp53,* which is the mouse ortholog of *tumor suppressor protein 53* (*TP53*) (**Fig. 4e**). Indeed, the top-master regulator was NOP53, whose predicted inhibition activates TP53^40^ (**Fig. 4e**). We also confirmed a synergistic elevation of Trp53 mRNA expression in the microglia isolated from mPFC by quantitative real time PCR (qPCR) (THC × 16p11dup interaction for Trp53 mRNA expression [*F*_1,20_ = 13.62, *p* = 0.0015]) (**Fig. 4f**).

**Fig. 4.**
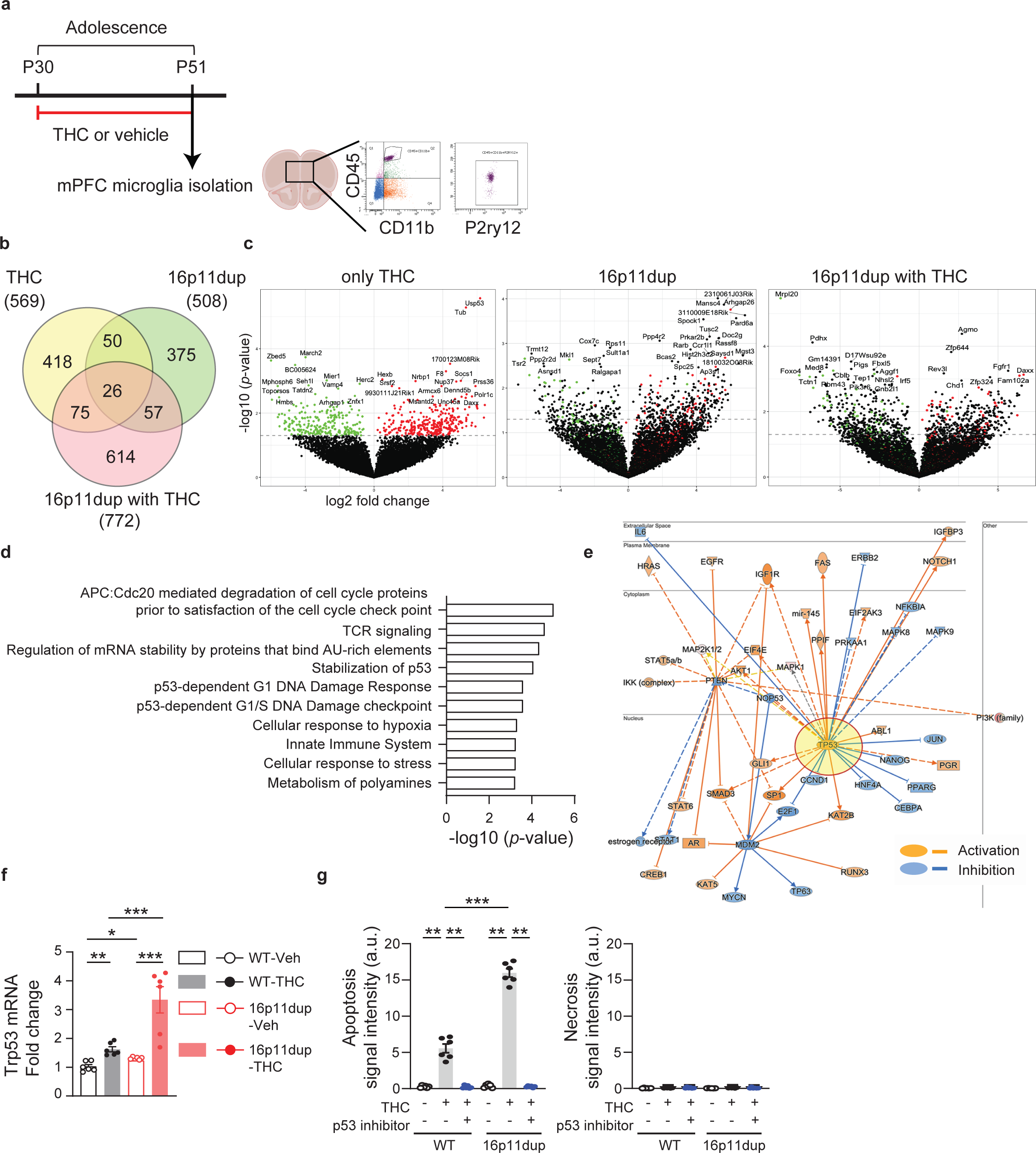
Up-regulation of p53 signaling pathway identified by mPFC microglia-specific transcriptome profiling. **a,** Experimental flow of Fluorescence-Activated Cell Sorting (FACS)-based mPFC microglia isolation by using 3 markers (CD45/CD11b/P2ry12) at P51, followed by RNA sequencing (RNA-seq). **b**, Venn diagram showing the overlap of differentially expressed genes (DEGs, with uncorrected p<0.05) in THC treatment or 16p11dup alone, and 16p11dup exposed to THC (16p11dup with THC), compared to controls. *p* < 0.05. *n* = 5 mice per condition. **c**, Volcano plot representations of the differential expression and pathway enrichment analyses of THC alone, 16p11dup alone, and 16p11dup with THC showing down-regulated genes (green), up-regulated genes (red), and not significantly differentially expressed genes (black). **d**, Significantly enriched pathways of 16p11dup with THC in the Reactome Pathway analysis. **e,** Functional gene interaction network from the upstream regulators / Ingenuity Pathway Analysis (IPA) for DEGs in the 16p11dup with THC treatment condition showing predicted inhibition (blue), activation (orange), and unknown directionality (grey). **f,** Relative mRNA expression level of Trp53 in microglia isolated from mPFC at P51. *n* = 6 mice per condition. **g,** Apoptosis (left) and necrosis (right) assays using primary microglia cultures treated by THC or vehicle with and without pifithrin-α. (Left) Quantification of signal intensity of apopxin. *n* = 6 fields in 3 mice per condition. (Right) Quantification of signal intensity of 7-AAD. *n* = 6 fields in 3 mice per condition. (**f, g**) \*\*\**p* < 0.001, \*\**p* < 0.01, \**p* < 0.05, determined by two-way ANOVA with post hoc Tukey test. Data are presented as the mean ± s.e.m.

To obtain direct evidence that activation of the p53 signaling pathway may play a role in mediating the GxE effect on microglial apoptosis, primary microglia cultures were produced from 16p11dup mice and littermate controls and treated with pifithrin-α (10μM), an inhibitor of p53 transcriptional activity, for 7 hours, starting from 1 hour before the THC or vehicle treatment. These cells were subjected to apoptosis and necrosis assays as described above. We observed that inhibition of p53 signaling prevented THC-induced microglial apoptosis and apoptotic phenotypes produced by THC treatment and 16p11dup (THC × 16p11dup interaction for microglial apoptosis [*F*_1,20_ = 148.8, *p* < 0.0001]) (**Fig. 4g, Supplementary Fig. 3g**). These results suggest that up-regulation of the p53 signaling pathway in microglia is the GxE mechanism whereby adolescent THC treatment interacts with 16p11dup to produce microglial apoptosis.

### Impairment of social memory produced by adolescent THC treatment and 16p11dup

We next examined whether adolescent THC treatment produces adult behavioral outcomes in 16p11dup mice after a 3-week abstinence period (**Fig. 5a**). Previous studies have reported that 16p11dup mice exhibit social behavioral deficits^20^. We found no effect of adolescent THC treatment and 16p11dup on sociability in the three-chamber social interaction test in male mice (**Fig. 5b, Supplementary Fig. 4a**). However, 16p11dup male mice exhibited less preference for social novelty compared to controls. This effect was enhanced by adolescent THC treatment (THC × 16p11dup interaction for social novelty [*F*_1,50_ = 6.392, *p* = 0.0147]) (**Fig. 5c, Supplementary Fig. 4b**). Interestingly, although some studies have reported that female mice are more vulnerable to behavioral effects of THC treatment^41, 42^, we did not observe these behavioral phenotypes in the female mice (**Supplementary Fig. 4a, b**). We further examined the impact of adolescent THC treatment and 16p11dup on social behaviors in the male mice by the 5-trial social memory test^43^. The 16p11dup × THC group exhibited no significant changes in habituation (decreased exploration) during the first four trials, but showed impaired recognition of novel mice on the fifth trial compared to other groups of mice (THC × 16p11dup interaction for social novelty [*F*_1,50_ = 5.665, *p* = 0.0239]) (**Fig. 5d**). Neither THC nor 16p11dup produced abnormalities in locomotion and in anxiety-like phenotypes, as assessed by the open field test and elevated plus maze test in both male and female mice (**Supplementary Fig. 4c**). Importantly, there were no abnormalities in the habituation/dis-habituation test, novel object recognition test, nor novel place recognition tests among the four groups of mice (**Fig. 5e, Supplementary Fig. 4d, e**), suggesting the specific effects of GxE on social memory.

**Fig. 5.**
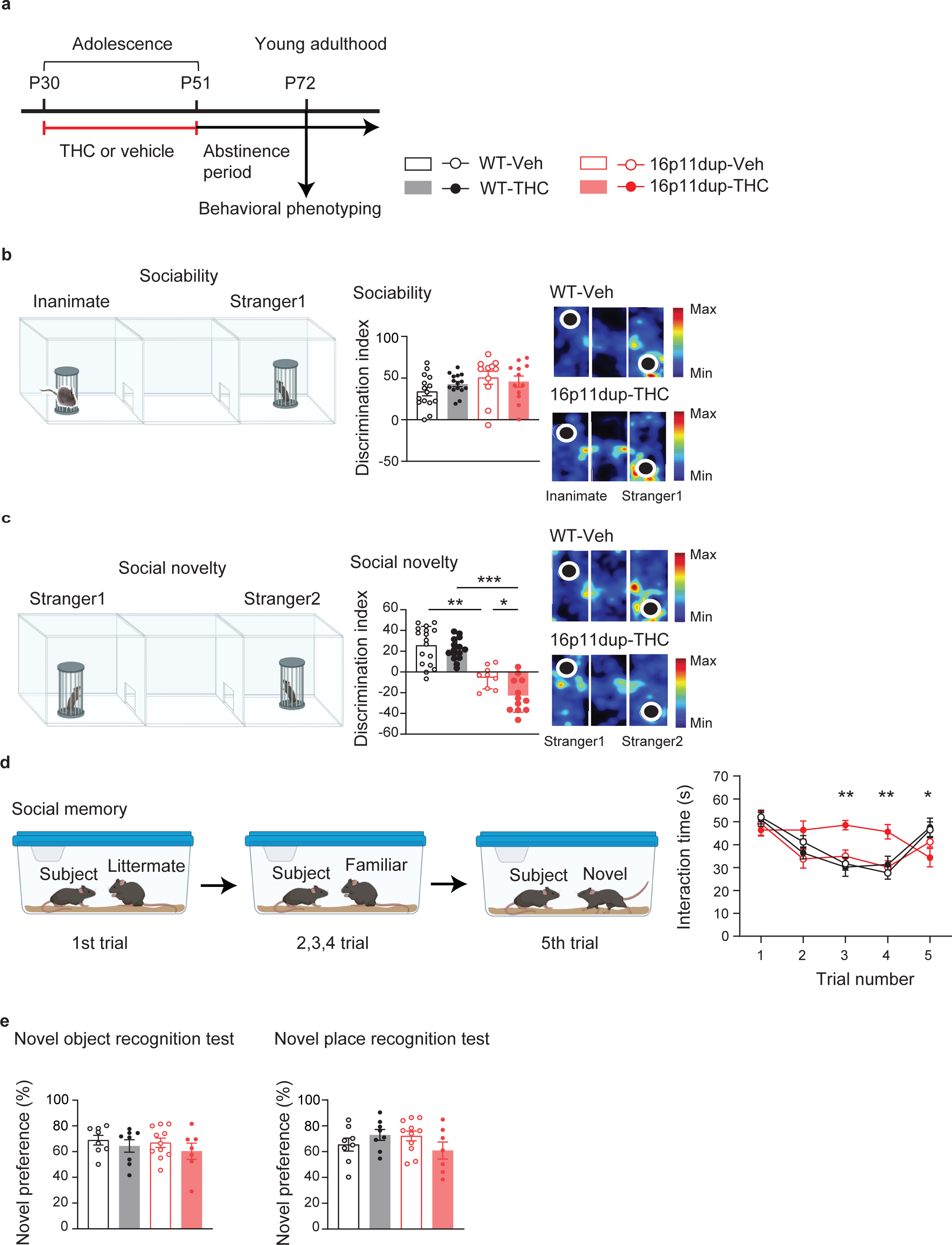
Social novelty recognition and memory deficits synergistically produced by adolescent THC treatment and 16p11dup. **a,** Schematic diagram of the adolescent THC treatment protocol. 16p11dup mice and wild type littermate controls (WT) were treated with THC (s.c., 8mg/kg) or vehicle (Veh) during adolescence (P30-P51), followed by behavioral assays at P72 after a 3-week abstinence period at adulthood. **b,** (Left) Schematic diagram of the three-chamber social interaction test for sociability. (Middle) Sociability phenotypes in WT and 16p11dup mice with adolescent THC or Veh treatment as indicated by discrimination index ([Stranger 1 – Inanimate mice sniffing time] / [Stranger 1 + Inanimate mice sniffing time] × 100 (%)). (Right) Representative heat maps depict movements of the WT-Veh mice vs. 16p11dup-THC mice. **c,** (Left) Schematic diagram of the three-chamber social interaction test for social novelty preference. (Middle) Preference of social novelty in WT and 16p11dup mice with adolescent THC or Veh treatment as indicated by discrimination index ([Stranger 2 – Stranger 1 sniffing time] / [Stranger 2 + Stranger 1 sniffing time] × 100 (%)). (Right) Representative heat maps depict movements of the WT-Veh vs. 16p11dup-THC mice. **d,** (Left) Schematic diagram of the 5-trial social memory test. (Right) Interaction time of WT and 16p11dup mice receiving adolescent THC or Veh treatment with an ovariectomized female mouse. (**b**, **c**, **d**) WT-Veh (*n* = 17 mice), WT-THC (*n* = 15 mice), 16p11dup-Veh (*n* = 11 mice), and 16p11dup-THC (*n* = 11 mice). **e,** (Left) Preference of novel object in novel object recognition test (NORT) in WT and 16p11dup mice with adolescent THC or Veh treatment. (Right) Preference of novel place in novel place recognition test (NPRT) in WT and 16p11dup mice with adolescent THC or Veh treatment. (**e**) WT-Veh (*n* = 8 mice), WT-THC (*n* = 8 mice), 16p11dup-Veh (*n* = 11 mice), 16p11dup-THC (*n* = 7 mice). (**b, c, d, e**) \*\*\**p* < 0.001, \*\**p* < 0.01, \**p* < 0.05, determined by two-way ANOVA with post hoc Tukey test. Data are presented as the mean ± s.e.m.

In order to examine whether the above synergistic impairment of social memory was dependent on adolescent THC exposure, 16p11dup male mice and littermate male controls were chronically treated with THC in adulthood (P70-P91) using the same protocol (**Supplementary Fig. 5a**). Adult THC treatment reduced the number of microglia and produced morphological changes in the microglia of the mPFC at P91, which was similar to the outcomes in the mice with adolescent THC treatment (**Supplementary Fig. 5b-d**). However, there was no effect of adult THC treatment on social memory after a 3-week abstinence period in neither 16p11dup mice nor wild type littermates (**Supplementary Fig. 5e-g**). These results suggest that adolescence is a critical period for the adverse effect of microglial apoptosis on social memory.

### Reduction in PT neuron excitability produced by adolescent THC treatment and 16p11dup

Recent studies reported that pyramidal-tract (PT) neurons in layer 5 of mPFC contribute to modulating social behaviors^44^. Although microglial phenotypes are specifically observed in the mPFC at P51, but not at P72 after a 3-week abstinence period from THC treatment (**Fig. 2, Supplementary Fig. 2**), synergistic deficits in social memory emerged at P72 (**Fig. 5**). Therefore, we next examined the impact of adolescent THC treatment and 16p11dup on neuronal function in the mPFC by whole-cell patch clamp recording in acute PFC slices obtained from male 16p11dup and littermate control mice at P72 (**Fig. 6a**). We compared the responses of two major subtypes of layer 5 prefrontal pyramidal neurons, PT neurons and intra-telencephalic (IT) neurons^45, 46^, whose cell-type identities are distinguishable based on their sag amplitude and kinetics (**Fig. 6b**). We measured basal membrane properties and evoked action potentials in the presence of glutamatergic and GABAergic synaptic blockers. We observed that adolescent THC treatment interacts with 16p11dup to reduce input resistance, increase rheobase, and reduce action potential firing frequency in PT neurons as signs of decreased intrinsic excitability (THC × 16p11dup interaction for input resistance [*F*_1,39_ = 5.805, *p* = 0.0208] and rheobase [*F*_1,39_ = 4.353, *p* = 0.0435]) (**Fig. 6c**). However, there were no significant effects of THC treatment, 16p11dup, or both combined on the functional properties of IT neurons (**Fig. 6d**). To ascertain whether these adverse THC effects are adolescent treatment-specific, we used the same approach to examine adult THC treatment-induced effects on PT and IT neuron function via electrophysiological assays. We observed no significant effects of adult THC treatment, 16p11dup, or both combined on functional properties of PT and IT neurons including input resistance, rheobase, and action potential firing frequency (**Supplementary Fig. 5h, i**). These results suggest that adolescence is a critical period for THC’s adverse effects on PT neuron function in the mPFC of 16p11dup mice in adulthood.

**Fig. 6.**
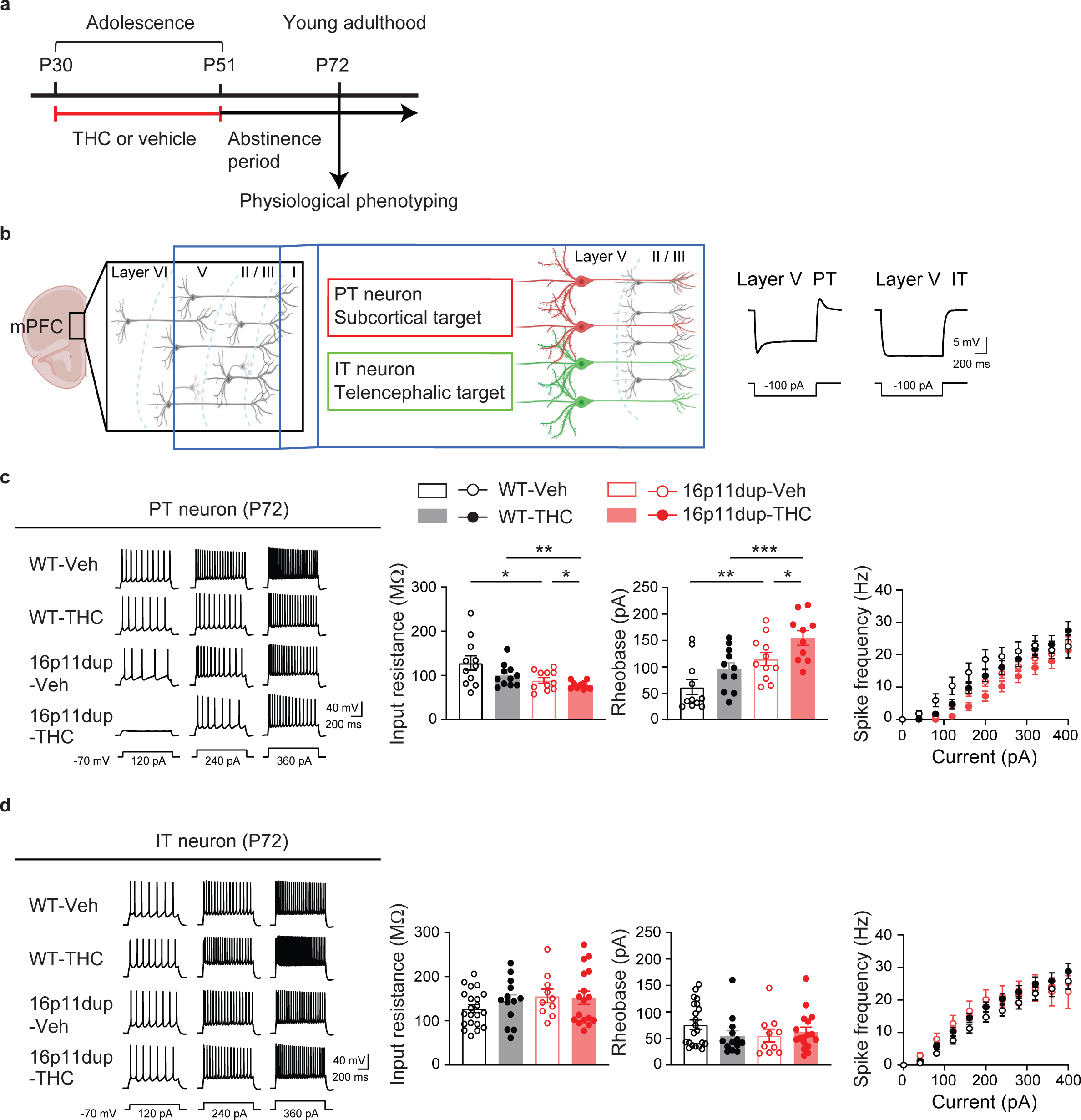
Synergistic reduction in intrinsic excitability in PT neurons produced by adolescent THC and 16p11dup. **a,** Schematic diagram of the adolescent THC treatment protocol. 16p11dup mice and wild type littermate controls (WT) were treated with THC (s.c., 8mg/kg) or vehicle (Veh) during adolescence (P30-P51), followed by electrophysiological assays at P72 after a 3-week abstinence period at adulthood. **b,** (Left) Schematic structure of multiple layers of the mPFC. Pyramidal tract (PT, red) and intra-telencephalic (IT, green) neurons in layer 5 of the mPFC are depicted. (Right) Representative voltage traces elicited by applying -100 pA current steps. These traces show typical hyperpolarizing responses of PT and IT neurons. **c,** (Left) Representative voltage traces recorded from PT neurons in response to current step injections in the presence of blockers of AMPA, NMDA, and GABAa receptors. (Right) The intrinsic excitability assessed by measurement of input resistance (left), rheobase (middle), and spike frequency (right). WT-Veh (*n* = 11 cells in 6 mice), WT-THC (*n* = 11 cells in 7 mice), 16p11dup-Veh (*n* = 11 cells in 6 mice), and 16p11dup-THC (*n* = 10 cells in 5 mice). **d,** (Left) Representative voltage traces recorded from IT neurons in response to current step injections in the presence of blockers of AMPA, NMDA, and GABAa receptors. (Right) The intrinsic excitability assessed by measurement of input resistance (left), rheobase (middle), and spike frequency (right). WT-Veh (*n* = 21 cells in 8 mice), WT-THC (*n* = 13 cells in 7 mice), 16p11dup-Veh (*n* = 10 cells in 6 mice), and 16p11dup-THC (*n* = 17 cells in 7 mice). (**c**) \*\*\**p* < 0.001, \*\**p* < 0.01, \**p* < 0.05, determined by two-way ANOVA with post hoc Tukey test. Data are presented as the mean ± s.e.m.

### Microglial *Cnr1* mediates the microglial phenotypes, PT neuron impairments, and social behavioral deficits produced by adolescent THC treatment and 16p11dup

To determine whether *Cnr1* expression in microglia causally mediates the above-mentioned phenotypes produced by adolescent THC treatment and 16p11dup, we used 16p11dup mice crossed with *Cnr1^flox/flox^*mice and the *Cx3cr1^CreER^* line to generate the following 4 groups: (i) wild type littermate (16p11^wt^); *Cx3cr1^CreER/+^*;*Cnr1^+/+^*, (ii) 16p11^wt^;*Cx3cr1^CreER/+^*;*Cnr1^flox/flox^*, (iii) 16p11^dup^;*Cx3cr1^CreER/+^*; *Cnr1^+/+^*, and (iv) 16p11^dup^;*Cx3cr1^CreER/+^*;*Cnr1^flox/flox^*. For induction of the microglial deletion of *Cnr1* as previously shown, tamoxifen (0.1 mg/g body weight) was given orally once a day for 5 consecutive days at P21-P25. The animals were then subjected to adolescent THC treatment (P30-P51), followed by assessing microglial phenotypes at P51 (**Fig. 7a**). We observed that genetic deletion of *Cnr1* in microglia ameliorated microglia reduction as well as microglial morphological changes and apoptosis in the mPFC produced by adolescent THC treatment in 16p11dup mice or wild type littermates (**Fig. 7b, c, d, e, f**). Using an independent cohort of these groups of mice, we also examined whether the THC effect on microglial Cnr1 was causally linked to impairments in the intrinsic excitability of mPFC PT neurons and deficits in social recognition and memory after a 3-week THC abstinence period (**Fig. 8a**). We found that genetic deletion of *Cnr1* in microglia reversed the functional deficits of PT neurons induced by adolescent THC treatment in 16p11dup mice, with no effects on IT neurons (**Fig. 8b, c**). At the behavioral level, while no changes were seen in spontaneous locomotor activity or olfaction (**Supplementary Fig. 6**), genetic deletion of *Cnr1* in microglia normalized deficits in social memory synergistically produced by adolescent THC treatment and 16p11dup (**Fig. 8d, e**). Although *Cnr1* is also expressed in other cell types (**Fig. 1d**), these results suggest that microglial Cnr1 expression contributes not only to adolescent THC treatment-induced microglial abnormalities, but also to impairments in the intrinsic excitability of PT neurons and deficits in social memory in 16p11dup mice.

**Fig. 7.**
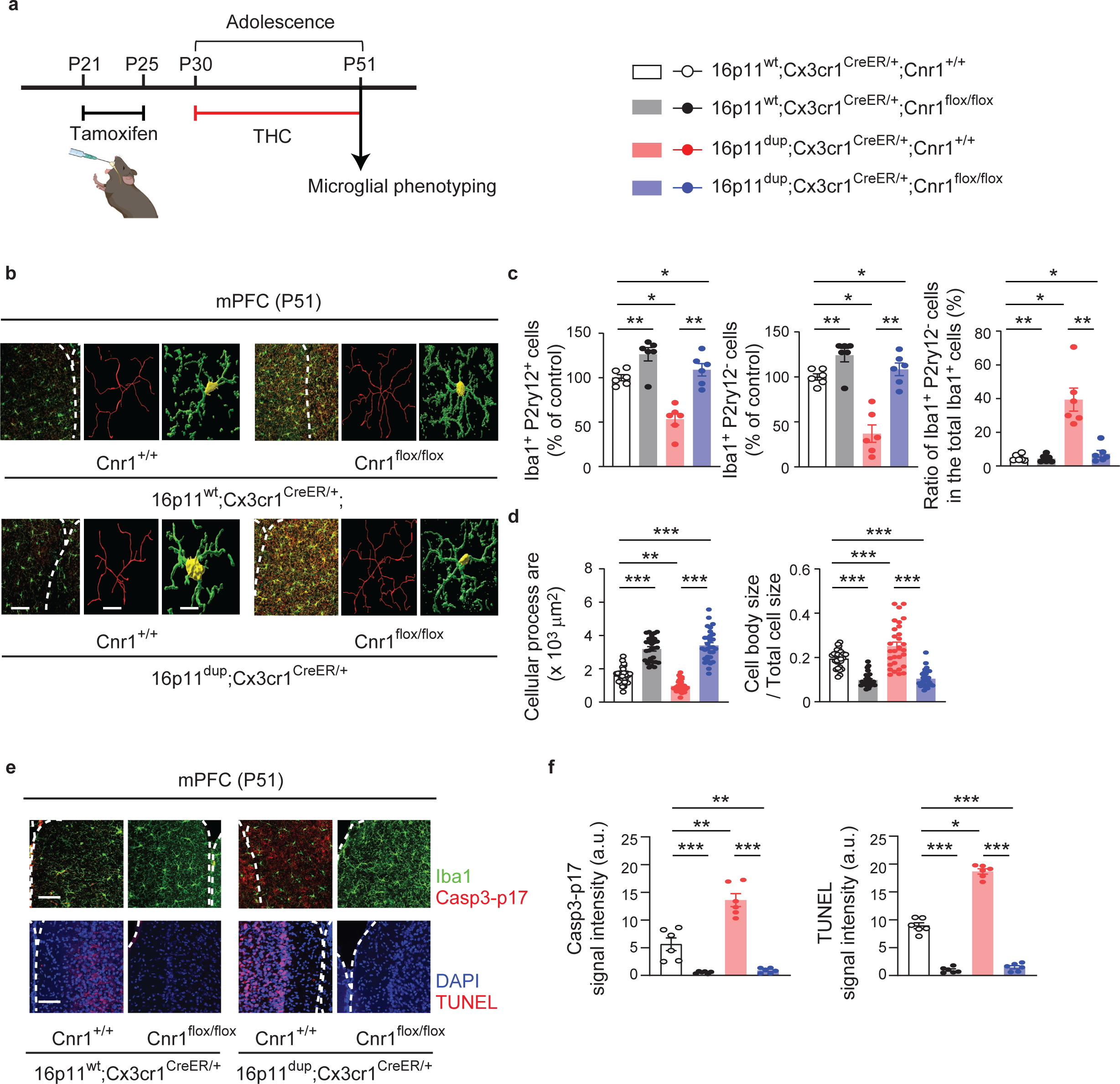
*Cnr1* deletion in the microglia normalizes microglial abnormalities produced by adolescent THC treatment and 16p11dup. **a,** Schematic diagram of the experimental design. 16p11^dup^;*Cx3cr1^CreER/+^*;*Cnr1^+/+^*, 16p11^dup^;*Cx3cr1^CreER/+^*;*Cnr1^flox/flox^*, wild type littermate (16p11^wt^);*Cx3cr1^CreER/+^*;*Cnr1^+/+^*, and 16p11^wt^;*Cx3cr1^CreER/+^*;*Cnr1^flox/flox^* mice were given tamoxifen (0.1 mg/g body weight) orally once a day for 5 consecutive days at P21-P25, 5 days before the start of THC (s.c., 8mg/kg) treatment during adolescence (P30-P51), followed by microglial phenotyping at P51 (upon completion of THC treatment). **b,** Immunohistochemical analysis of Iba1 (green) in the mPFC at P51. (Left) Representative images of the mPFC, representative tracing images (red), and images of cellular processes (green) and cell bodies (yellow) of Iba1^+^ cells. Scale bar, 50 μm (left) and 10 μm (middle and right). **c,** The number of Iba1^+^P2ry12^+^ cells (left) and Iba1^+^P2ry12^-^ cells (middle) in the mPFC, presented as % of control. (Right) The percentage of Iba1^+^P2ry12^-^ cells among all Iba1^+^ cells in the mPFC. (*n* = 6 slices in 3 mice per condition). **d,** Quantification of the ratio of cellular process area (left) and cell body size to total cell size (right) of Iba1^+^ cells. (*n* = 30 cells in 3 mice per condition). **e,** Representative images of immunohistochemistry of Iba1 (green) and Casp3-p17 (red) (top left) as well as TUNEL signals (red) and DAPI (blue) (bottom left) in the mPFC at P51. Scale bar, 50 μm. **f**, Quantification of signal intensity of Casp3-p17 in Iba1^+^ cells (left) and TUNEL (right). *n* = 6 slices in 3 mice per condition. (**c, d, f**) \*\*\**p* < 0.001, \*\**p* < 0.01, \**p* < 0.05, determined by two-way ANOVA with post hoc Tukey test. Data are presented as the mean ± s.e.m.

**Fig. 8.**
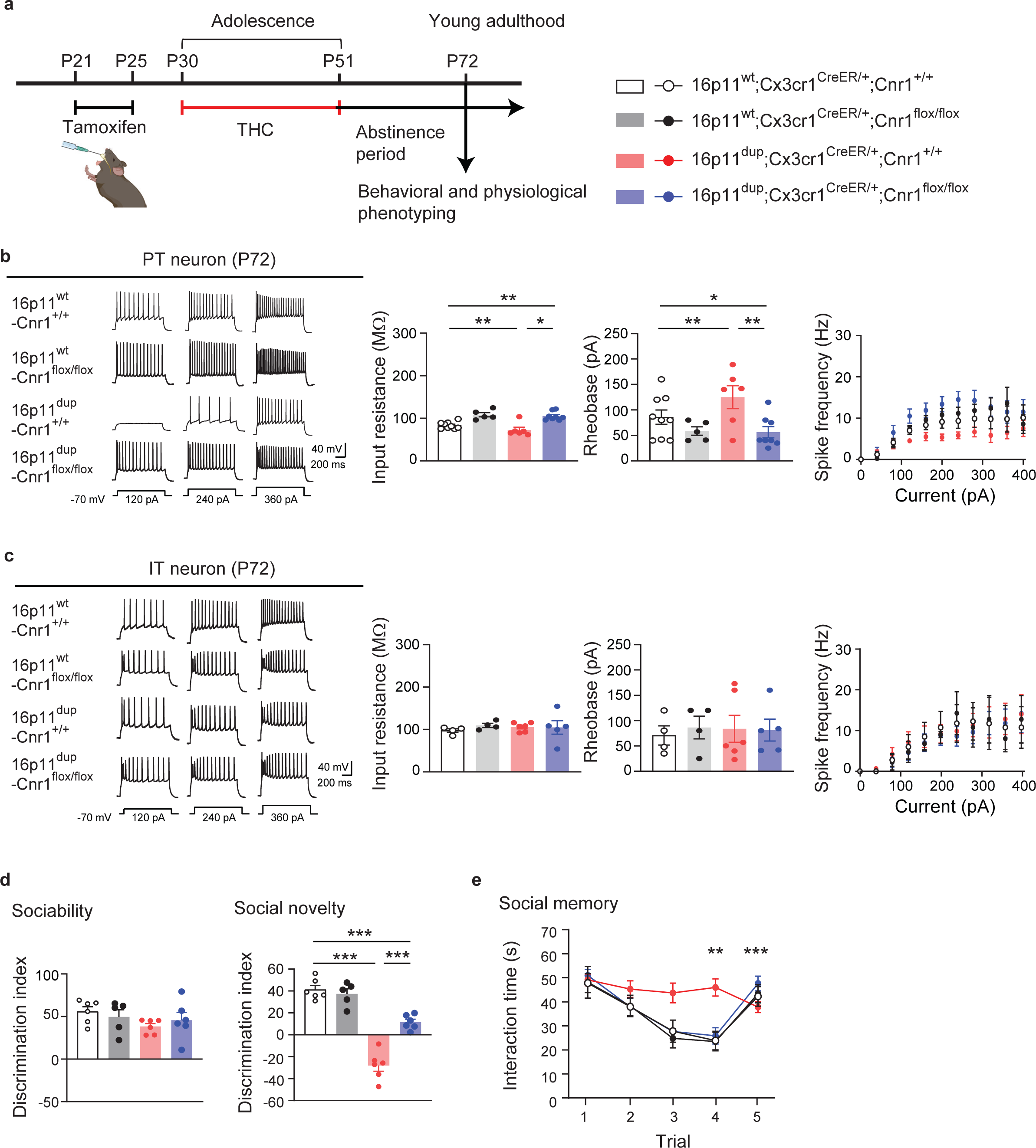
*Cnr1* deletion in the microglia normalizes deficits in PT neurons and social memory that are synergistically produced by adolescent THC treatment and 16p11dup. **a**, Schematic diagram of the experimental design. 16p11^dup^;*Cx3cr1^CreER/+^*;*Cnr1^+/+^*, 16p11^dup^;*Cx3cr1^CreER/+^*;*Cnr1^flox/^*^flox^, wild type littermate (16p11^wt^);*Cx3cr1^CreER/+^*;*Cnr1^+/+^*, and 16p11^wt^;*Cx3cr1^CreER/+^*;*Cnr1^flox/flox^*mice were given tamoxifen (0.1 mg/g body weight) orally once a day for 5 consecutive days at P21-P25, 5 days before the start of THC (s.c., 8mg/kg) treatment during adolescence (P30-P51), followed by electrophysiological experiments or behavioral assays at P72 (after a 3-week abstinence period at adulthood). **b,** (Left) Representative voltage traces recorded from PT neurons in response to current step injections in the presence of blockers for AMPA, NMDA, and GABAa receptors. (Right) The intrinsic excitability of PT neurons, as quantified by input resistance (left), rheobase (middle), and spike frequency (right). 16p11^wt^;*Cx3cr1^CreER/+^*;*Cnr1^+/+^*(*n* = 9 cells in 2 mice), 16p11^wt^;*Cx3cr1^CreER/+^*;*Cnr1^flox/flox^*(*n* = 5 cells in 3 mice), 16p11^dup^; *Cx3cr1^CreER/+^*;*Cnr1^+/+^*(*n* = 6 cells in 2 mice), and 16p11^dup^;*Cx3cr1^CreER/+^*;*Cnr1^flox/flox^*(*n* = 8 cells in 3 mice). \*\**p* < 0.01, \**p* < 0.05, determined by two-way ANOVA with post hoc Tukey test. **c,** (Left) Representative voltage traces recorded from IT neurons in response to current step injections in the presence of blockers for AMPA, NMDA, and GABAa receptors. (Right) The intrinsic excitability of IT neurons, as quantified by input resistance (left), rheobase (middle), and spike frequency (right). 16p11^wt^;*Cx3cr1^CreER/+^*;*Cnr1^+/+^*(*n* = 4 cells in 2 mice), 16p11^wt^;*Cx3cr1^CreER/+^*;*Cnr1^flox/flox^*(*n* = 4 cells in 2 mice), 16p11^dup^;*Cx3cr1^CreER/+^*; *Cnr1^+/+^* (*n* = 6 cells in 2 mice), and 16p11^dup^;*Cx3cr1^CreER/+^*;*Cnr1^flox/flox^*(*n* = 5 cells in 2 mice). **d,** Sociability phenotypes (left) and preference of social novelty (right) as indicated by the discrimination index in the three-chamber social interaction test. **e,** Time spent with an ovariectomized female mouse in the 5-trial social memory test. (**d**, **e**) 16p11^wt^;*Cx3cr1^CreER/+^*;*Cnr1^+/+^* (*n* = 6 mice), 16p11^wt^;*Cx3cr1^CreER/+^*; *Cnr1^flox/flox^*(*n* = 5 mice), 16p11^dup^; *Cx3cr1^CreER/+^*;*Cnr1^+/+^*(*n* = 6 mice), 16p11^dup^;*Cx3cr1^CreER/+^*; *Cnr1^flox/flox^*(*n* = 6 mice). (**d, e**) \*\*\**p* < 0.001, \*\**p* < 0.01, determined by two-way ANOVA with post hoc Tukey test. Data are presented as the mean ± s.e.m.

## Discussion

Here, we showed that adolescent THC exposure and 16p11dup interact to produce Cnr1-mediated microglial apoptosis in the mPFC, leading to aberrant PT neuron function and social cognitive deficits in adulthood. Our findings demonstrated for the first time the pathological role of microglial Cnr1 in mediating adverse cognitive effects of adolescent cannabis use in the context of GxE. Previous studies have reported that treatment with relatively high doses of THC at 5 mg/kg or 10 mg/kg has induced short-lived reduction in locomotor activity in the wake of THC treatment^47, 48^. These results suggest that reduced locomotion acutely induced by THC treatment is transient and may not affect cognitive deficits observed after a 3-week abstinence period. Adolescence is a critical period for PFC development, whereby microglia mediate neuronal maturation via synaptic pruning and remodeling, contributing to higher cognitive function^12–16, 49, 50^. Furthermore, adult cognitive function is greatly influenced by various environmental factors during developmental periods, such as adolescent cannabis use^1, 3^. Thus, elucidating microglia-mediated pathways during adolescence is important for understanding the complex nature of pathophysiological mechanisms underlying the impact of adolescent cannabis use on cognitive impairments and psychiatric disorders.

It has been shown that subchronic THC treatment during adolescence increases Iba1 expression and induces neuroinflammation in the PFC of female rats^41, 51^. Therefore, we were surprised to find that adolescent THC exposure induced microglial apoptosis in the mPFC of male mice, but not of female mice. These results suggest that microglial phenotypes induced by adolescent THC treatment depend on multiple factors, such as the duration of THC exposure, dose, abstinence periods, and/or sex.

Interestingly, although we identified a decrease in the total number of Iba1^+^ microglia following THC treatment, the percentage of Iba1^+^P2ry12^-^ cells in the total microglia increased. These results are consistent with the morphological transition of microglia from a ramified to an amoeboid-like phenotype upon completion of THC treatment. The NF-κB signaling pathway has previously been identified as an upstream mediator for THC-induced apoptosis of dendritic cells^32^. p53-deficient microglia have also shown an increased expression of genes involved in anti-inflammation and tissue repair^52^. Additionally, pharmacological microglial activation up-regulates p53 expression^53^. Thus, examining other markers of microglial activation^54, 55^ and studying the mechanistic links between apoptotic and inflammatory signaling pathways in microglia is needed to understand the microglial mechanisms underpinning the adverse effects of THC. Given that enzymatic dissociation of brain tissue during the process of cell isolation affects molecular expression in microglia^56–59^, interpreting such data requires caution. Our results also showed that THC-induced microglial apoptosis is mediated by Cnr1 expressed in the microglia. Notably, although THC also activates Cnr2 that is primarily (but not exclusively) expressed in various immune cells, including microglia^39^, our results showed that *Cnr2* deletion does not affect THC-induced microglial apoptosis.

Microglia have increasingly been reported to exhibit molecular heterogeneity across brain regions^60^. Compared to other regions, Cx3cr1 and CD80, which are involved in microglial activation, were found to be highly expressed in the PFC^61, 62^. Interestingly, lipopolysaccharide (LPS) administration induced a higher expression of pro-inflammatory cytokines in the PFC, compared to other brain regions^63^. These studies suggest that the PFC may be uniquely sensitive to microglia-mediated inflammatory responses, potentially contributing to the THC-induced phenotypical changes seen in mPFC microglia.

The adverse effects of THC varied depending on the genetic risks conferring susceptibility to psychiatric disorders^25, 26, 28^. THC-induced microglial apoptosis was enhanced by 16p11dup, but not 22q11 deletion, suggesting that specific genetic risks underlie microglial vulnerability to cannabis exposure. Interestingly, previous studies, including ours, reported that many genes in the 16p11dup risk loci, including *Coro1a, Mapk3*, *Mvp*, *Tmem219*, *Kctd13*, *Maz*, *Taok2*, *Ypel3*, and *Aldoa*, are involved in apoptotic cell death or p53 signaling pathways^64–72^. This risk locus also contains multiple genes including*, Mapk3, Gdpd3, Ino80e, Hirip3, Mvp, Maz, and Spn*, which are highly expressed in microglia, as we found in a large-scale single-cell RNA-seq dataset from juvenile mouse cortices^73^. These results suggest that 16p11 CNV risk may contribute to increasing microglia’s susceptibility to apoptosis. While understanding the biological impact of CNVs may provide us with unprecedented insight into the complex mechanisms of psychiatric disorders, only some people with CNVs develop cognitive impairments, suggesting roles for environmental risk factors and GxE. This notion is supported by current work demonstrating that microglial apoptosis induced by adolescent THC exposure and 16p11dup yields long-lasting dysfunction in mPFC PT neurons and deficits in social memory. It will be important to further investigate the critical genes in 16p11dup loci whose dosage sensitivity plays the driving role in inducing microglial apoptosis in conjunction with THC-induced Cnr1 signaling alterations, leading to neuronal and cognitive deficits.

Aberrant prefrontal structure and function are implicated in psychiatric disorders of neurodevelopmental origins, such as autism spectrum disorder and schizophrenia^74^. Social cognitive impairments are often exhibited in patients with these disease conditions^75^. Recent preclinical evidence indicates that the mPFC is highly interconnected with subcortical regions, contributing to information processing involved in social recognition and memory. For instance, proper synaptic input from the ventral hippocampus onto layer 5 pyramidal neurons of mPFC is crucial in regulating social memory^76^. The nucleus accumbens-projecting mPFC neurons modulate social novelty recognition^77^. Although our current study supports the importance of PT neurons in the regulation of social memory, it is unclear how the adverse effects of THC on microglia selectively impact PT neurons, but not IT neurons, in the 16p11dup mice model. Recent studies have shown that microglia engulf the presynaptic elements and dendritic spines of pyramidal neurons in layer 5 of the mPFC during adolescence^13^. Thus, it is of interest to further investigate the differential microglia-mediated maturation processes of these neuronal subtypes and to elucidate how PT neurons specifically regulate social memory.

Although *Cx3cr1^CreER/+^*mice are widely used as a Cre driver line for studying microglia, Cnr1 expression in non-microglial Cx3cr1^+^ cells may also be suppressed in *Cx3cr1^CreER/+^*;*Cnr1^flox/flox^* mice, warranting the use of other Cre lines targeting microglia to further investigate the precise roles of microglial Cnr1^78^.

In summary, our findings highlight the unexplored impact of adolescent THC exposure on microglial function. Furthermore, we identified the role of Cnr1 expressed in microglia for mediating its GxE effect on adolescent mPFC maturation and adult social memory. With the growing interest in the pathological implication of microglia in a wide range of psychiatric disorders, there is a clear need to further investigate the precise role of microglia in mediating GxE in the context of adolescent cannabis exposure. Considering that cannabis is an increasingly popular drug for medical and recreational use^79^, these lines of study will be particularly critical for building a mechanistic understanding of the adverse effects of cannabis, as well as providing translational insight for clinical research to address how cannabis exposure contributes to psychopathology in individuals genetically predisposed to psychiatric disorders.

## METHODS

### Mice

C57BL/6J and *Cx3cr1^CreER/CreER^* (*Cx3cr1^tm^*^2^.^1^(cre/ERT2)*^/Litt^*) mice were purchased from Jackson Laboratories (JAX; Bar Harbor, ME, USA). The mouse model of 16p11dup was originally developed by Drs. Wigler and Mills’s group^18^ and 22q11del mouse model was originally provided by Dr. Antonio Baldini^80^. *Cnr1^flox/flox^* mice were a gift from Dr. George Kunos (originally developed by Lutz’s lab), and *Cnr2* KO mice were a gift from Dr. Ken Mackie^81^. These genetically modified lines were backcrossed onto the C57BL/6J background. For genetic deletion of *Cnr1* in the microglia, wild type (16p11^wt^);*Cx3cr1^+/+^*;*Cnr1^flox/+^*female mice were crossed with male 16p11^dup^;*Cx3cr1^CreER/CreER^*; *Cnr1^flox/+^* mice. All genetically engineered mice used for experiments were genotyped to verify the presence of genetic components specific for each mouse line. When purchased from JAX, animals were maintained for at least 1 week to allow for habituation before experimentation. When both male and female mice were used, animals were sex- and age-matched in each experiment. All animal procedures were approved by the Institutional Animal Care and Use Committee of Johns Hopkins University School of Medicine, and adhered to ethical consideration in animal research.

### Drug administration (THC and tamoxifen)

Delta-9-tetrahydrocannabinol (THC) was obtained from RTI International through the NIDA Drug Supply Program and Cayman Chemical. Using previously established protocols^25–28^, the mice were treated daily with a single subcutaneous injection (s.c.) of THC (8mg/kg) or vehicle during adolescence (Postnatal day 30 [P30]-P51) or adulthood (P70-P91). The THC solution was prepared with saline and Cremophor (18:1, saline:Cremophor). A control cohort was treated by injection of a matched saline and Cremophor (Vehicle) mixture. After chronic THC treatment, a drug washout period of 3 weeks was used for some experiments, including electrophysiology and behavioral tests, to minimize any direct effects from THC treatment. For induction of genetic deletion of *Cnr1*, tamoxifen (0.1 mg/g body weight) was given by oral gavage once a day for 5 consecutive days at P21-P25. The animals were then subjected to adolescent THC treatment, followed by behavioral tests after a 3-week abstinence period from THC treatment

### Fluorescence-Activated Cell Sorting (FACS)

For microglia-bulk RNA-seq and qPCR experiments, we prepared cell samples using FACS as previously published^82, 83^. Briefly, mPFC tissue was minced in Hibernate A low fluorescence reagent (BrainBits), and dissociated using mechanical dissociation methods. The resulting homogenates were pushed through a 70 μm cell strainer and centrifuged at 300×g for 10 min before removing supernatants. Cell pellets were resuspended and subjected to Miltenyi debris removal reagent (cat # 130-109-398, Miltenyi Biotec Inc) according to manufacturer instructions. Cell pellets were again resuspended in 50 μl FACS buffer (0.5% bovine serum albumin in PBS) for cell surface marker staining. The blocking of nonspecific binding was induced in the cell suspensions via incubation for 10 min at 4°C with anti–CD16/CD32 antibody (cat # 101320, BioLegend, clone 93, 5 ng/μl). This was followed by staining for 30 min at 4°C with the appropriate antibodies: PE-Cy7 rat anti-mouse CD45 (cat # 561868, BD Biosciences, clone 30-F11, 2 ng/μl), BV421 rat anti-mouse CD11b (cat # 562605, BD Biosciences, clone M1/70, 2 ng/μl), APC rat anti-mouse P2ry12 (cat # 848006, BioLegend, clone S16007D, 2 ng/μl), BV421 rat anti-mouse CD45 (cat # 103133, BioLegend, clone 30-F11, 2 ng/μl), PE rat anti-mouse CD11b (cat # 101208, BioLegend, clone M1/70, 2 ng/μl), APC rat anti-mouse ACSA-2 (cat # 130-117-535, Miltenyi Biotec, clone IH3-18A3, 1:25 dilution), AF 488 rat anti-mouse TMEM119 (cat # 53-6119-82, Invitrogen, clone V3RT1GOsz, 5 ng/μl), and Alexa 488 rat anti-mouse NeuN (cat # MAB377X, Millipore sigma, clone A60, 5 ng/μl). After incubation, cells were washed, then resuspended in 300 μl of FACS buffer. Gates were validated by 7-AAD Viability Staining Solution (cat # 00-6993-50, Invitrogen) to identify live and dead cells. CD45^+^/CD11b^+^/P2ry12^+^ or CD45^+^/CD11b^+^/TMEM119^+^ microglia, ACSA-2^+^ astrocytes, and NeuN^+^ neurons were acquired by a FACS Aria Flow Cytometer for further experimental use.

### Primary microglia culture

Microglia were isolated from whole-brain homogenates of different adult mouse models according to a previously described method with minor modification^84^. In brief, mice were anesthetized with isoflurane and transcardially perfused with 20 ml cold PBS. The brains were isolated, minced in HBSS (cat # 55021C, Sigma-Aldrich, St. Louis, MO, USA) and dissociated with the neural tissue dissociation kit (cat # 130-092-628, MACS Miltenyi Biotec, Auburn, CA) according to manufacturer instructions. After passing through a 70 μm cell strainer, resulting homogenates were centrifuged at 300 g for 10 min. Supernatants were removed, and cell pellets were resuspended. Myelin was removed by Myelin Removal Beads II (cat # 130-096-733, MACS Miltenyi Biotec, Auburn, CA) according to manufacturer instructions. Myelin-removed cell pellets were resuspended and incubated with CD11b MicroBeads (cat # 130-093-634, MACS Militenyi Biotec, Auburn, CA) for 15 min, then loaded onto LS columns and separated on a quadroMACS magnet. CD11b^+^ cells were flushed out from the LS columns, then washed and resuspended in the growth medium for mouse microglia (49 ml DMEM/F-12, 500 μl 100x Pen-strep/ glutamine stock, 500 μl TNS stock (including N-acetyl cysteine, sodium selenite, and apo-transferrin), 50 μl COG stock (including oleic acid, gondoic acid, and cholesterol), 50 μl TCH stock (including mouse CSF-1, human TGF-β2, and heparin sulfate))^85^. The number of viable cells was determined using a hemacytometer and 0.1% trypan blue staining. Each brain extraction yielded 1 × 10^6^ viable CD11b^+^ cells. These cells were then plated at a density of 1.4 × 10^5^ cells per well in a 12-well plate with cover glass (Cat # 12-545-100, Fisher Scientific) coated with PDL (15μg/ml) and collagen IV (cat # 354233, Corning), and placed in a humidified 37°C incubator with 5 % CO_2_. The microglial growth medium was changed every 2 days by removing 50 % of the medium and adding an equal volume of fresh microglial growth medium. 5 μM THC or Veh was added 7 days after seeding the cells. 6 hours later the cell culture was collected and cell morphology (Immunohistochemistry for phalloidin: cat # ab176753, Abcam) and apoptosis/necrosis (Apoptosis/Necrosis assay kit: cat # ab176749, Abcam) were assayed. In order to evaluate the contribution of p53 signaling to microglia apoptosis produced by 16p11dup with THC treatment, microglia culture cells were co-treated with THC and 10 μM of p53 signaling inhibitor, pifithrin-α (cat # P0122, Millipore SIGMA) for 1 h before THC or vehicle treatment.

### Isolation of microglia-enriched CD11b_+_ cells and ACSA-2_+_ astrocytes

Microglia-enriched CD11b^+^ cells and ACSA-2^+^ astrocytes were isolated from one adult mouse model according to a previously described method with minor modifications^84^. In brief, utilizing the same method used to create the primary microglia culture, CD11b^+^ cells were isolated by anti-CD11b MicroBeads (cat # 130-093-634, MACS Miltenyi Biotec, Auburn, CA). CD11b^+^ cells-removed cell pellets were resuspended and incubated with FcR blocking reagent for 10 min, followed by ACSA-2 MicroBeads (cat # 130-097-678, MACS Miltenyi Biotec, Auburn, CA) for 15 min, and then loaded onto LS columns where they were separated on a quadroMACS magnet. ACSA-2^+^ cells were flushed out from the LS columns, and ACSA-2^+^ cells-removed cell pellets were used as remaining cells.

### Protein extraction and immunoblotting

Microglia-enriched CD11b^+^ cells, ACSA-2^+^ astrocytes, and remaining cell populations were separately lysed in RIPA buffer (cat # 89901, ThermoFisher scientific) and sonicated to extract protein samples. Each protein sample was analyzed by SDA-PAGE and subsequent western blotting^25^. The following primary antibodies were used: rabbit polyclonal antibody against Cnr1 (cat # MSFR100590, CB1-Rb-Af380, Nittobo Medical Co), rabbit polyclonal antibody against Iba 1 (cat # 016-20001, FUJIFILM), chicken polyclonal antibody against GFAP (cat # ab4674, Abcam), rabbit monoclonal antibody against NeuN (cat # 24307, Cell Signaling Technology), and mouse monoclonal antibody against GAPDH (cat # sc-32233, Santa Cruz). Quantitative densitometric measurement of western blotting was performed by using ImageJ-FIJI software (NIH).

### Immunohistochemistry and TUNEL assay

Immunohistochemistry was performed using our previously published methods with some modifications^86, 87^. Mouse brains were extracted after perfusion with 4 % paraformaldehyde (PFA). The fixed brains were embedded in cryocompound (Sakura Finetek USA, Torrance, CA, USA) after replacement of PFA with 30% sucrose in phosphate buffered saline (PBS). Coronal sections including the mPFC were obtained at 100 µm with a cryostat (cat # CM 3050S, Leica, Buffalo Grove, IL, USA). The sections were heated in HistoVT One solution (cat # 06380-05, Nacalai Tesque, Kyoto, Japan) for 30 min at 60°C for antigen retrieval. The sections were then washed with PBS containing 0.5 % Triton X-100, followed by blocking with 0.5 % Triton X-100 and 1 % normal goat serum for 1 h. The sections were washed with PBS containing 0.5 % Triton X-100, then blocked with 0.5 % Triton X-100 and 1 % bovine skin gelatin for 1 h. After blocking, sections were incubated with primary antibody [Rabbit anti-Iba1 antibody (cat # 019-19471, Wako, 1:500), Rat anti-P2ry12 antibody (cat # 848001, BioLegend, 1:50), Rabbit anti-Aldh1l antibody (cat # MABN495, Millipore, 1:100), Rabbit anti-Casp3-p17 antibody (cat # sc-271028, SantaCruz, 1:100)] at 4°C overnight. An additional incubation with secondary antibodies conjugated to Alexa 488 (cat # A-32731, Invitrogen, 1:400) and Alexa 568 (cat # A-11004, Invitrogen, 1:400) was performed for 2 h. Nuclei were labeled with DAPI (cat # 10236276001, Roche, Indianapolis, IN, USA). TUNEL assay (cat # C10618, ThermoFisher Scientific) was performed according to the manufacturer’s instructions. As a positive control, adjacent brain sections were treated with 1 unit of DNase I (cat # 108068-015) diluted in 1X DNase I Reaction Buffer (20 mM Tris-HCl, pH 8.4, 2 mM MgCl2, 50 mM KCl) for 30 min at room temperature.

### Image analysis

Immunofluorescence images of mPFC were acquired using the Zeiss LSM700 confocal microscope with ZEN 2010 software (Carl Zeiss, Thornwood, New York, USA). PFC was defined as anteroposterior (AP): +2.57 to +1.53 mm, mediolateral (ML): ±2.75mm from bregma, dorsoventral (DV): −1.75 to −3.05mm from the dura according to the brain atlas. To assess the number of Iba1^+^ P2ry12^+^ cells and Iba1^+^ P2ry12^-^ cells, as well as to measure the signal intensity of Casp3-p17 in Iba1^+^ cells and TUNEL, all sections were imaged by the confocal microscope using the 20 × object lens to collect *z*-stacks image of twenty five optical slices at a step size of 1-μm thick. The signal intensity of Casp3-p17 was measured within thresholded Iba1^+^ mask to obtain the fluorescence intensity solely from Iba1^+^ microglia. To measure signal intensity of TUNEL assay, we created a 2D mask of mPFC and determined the mean fluorescence of TUNEL positive signal in all sections using ImageJ-FIJI software. Cells were counted and signal intensity was measured from two brain sections per mouse for each condition. Identical parameters for all imaging and threshold settings were used for all groups to minimize experimental bias. For the microglial morphology assessments, all sections were imaged on the confocal microscope using the 100 × oil immersion objective lens to collect *z*-stacks image of twenty five optical slices at a step size of 1-μm thick. Morphological analysis was performed by using Imaris 7.6.4 software (Bitplane, Zurich, Switzerland). 3D construction of microglia was applied, then the z-stack images were masked with the surface motif and reconstructed with the filament motif of Imaris software. We measured and compared the area of Iba1^+^ cell processes, total cell and cell body which were measured by using the filament function in the Imaris software. Iba1^+^ cell from two brain sections per mouse for each condition were used for analysis. These measurements were analyzed by experimenters who were blinded to the groups.

### RNA isolation and quantitative real-time PCR (qPCR)

Total RNA was isolated from mPFC tissue or FACS-sorted microglia cells using an RNeasy Mini Kit (QIAGEN). Quantitative real-time PCR (qPCR) was performed using TaqMan according to manufacturer’s protocol (Applied Biosystems). Briefly, cDNA from mPFC tissue or sorted microglia cells was prepared by SuperScript® III CellsDirect cDNA Synthesis Kit (Life Technologies) from total RNA in the range of 10-100 ng. Real-time PCR reactions contained diluted cDNA from the synthesis reaction and 200 nM of forward and reverse TaqMan primers specific to targets of interest (Assay IDs for Cnr1: Mm01212171_s1, Iba1: Mm00479862_g1, and Trp53: Mm01731290_g1, Applied Biosystems). Primers for GAPDH (Assay ID: Mm99999915_g1, Applied Biosystems) were used to normalize the expression data. The real-time PCR reaction and measurement was carried out with Applied Biosystems QuantStudio^TM^ 5. PCR reaction conditions are as follows: 50°C, 2 min; 95°C, 2 min; 50 cycles of 95°C, 1 sec; and 60°C, 20 sec, including a dissociation curve at the last step to verify single amplicon in the reaction.

### RNA sequencing (RNA-seq) and bioinformatic analysis

The microglia (CD11b^+^/CD45^+^/P2ry12^+^) were collected from the mPFC of mice at P51 (upon completion of adolescent THC treatment) by FACS using published methods^82, 88^. Batch effects were avoided by using balanced, randomized experimental design. These cells were transferred to the Johns Hopkins Sequencing core for subsequent processing (RNA isolation, quality control, library preparation for low input RNA (low input Nugen ovation RNA-seq V2)), followed by deep sequencing (50 million reads/sample, 150 bp paired end (PE) sequencing using NovaSeq) with standard RNA-seq pipeline at base resolution and at the transcript level. The software package “rsem-1.3.0” was used for running the alignments as well as generating gene and isoform expression levels. The ‘rsem-calculate-expression’ module was used with the following options: --star, --calc-ci, --star-output-genome-bam, --forward-prob 0.5. The data were then aligned to GRCm38/mm10 mouse reference genome. The outputs of this pipeline include count data and TPM and FPFM data, in addition to BAM formatted files of read alignments in genomic and transcript coordinates. We then performed differential expression analysis of RNA-seq data using the R package limma after voom normalization^89^, applying generalized linear models and adjusting for the percentage of mapped reads, as variable accounting for the overall sequencing accuracy, which is related to the RNA quality.

In this way, we identified specific genes that are more likely to be differentially expressed by 16p11dup, THC treatment, or the GxE effect when compared to WT littermate controls. We also used the R package ‘sva’ to identify surrogate variables accounting for unknown source of variations in our dataset^90^, and performed sensitivity analysis with five surrogate variables as covariates with consistent results (correlation between the t-statistics from the two differential expression analyses: t= 171.96, p<2e-16). We performed pathway enrichment analysis using the Reactome dataset (Panther 17.0 release); for this analysis we prioritized genes that are more likely to be differentially expressed based on uncorrected a value of *p* < 0.05, and log fold change <-2 or >2. Enriched pathways were identified based on FDR<0.05. We used the Ingenuity Pathway Analysis (IPA) software to detect common upstream regulators of differentially expressed genes, indicative of functional gene interaction networks associated with the interaction between THC and 16p11dup in the microglia. Because the upstream regulator analysis takes into account the magnitude and the directionality of the t-statistics from the differential expression analysis, we selected genes for this analysis only based on uncorrected a value of *p* < 0.05. Common upstream regulators were identified based on Benjamini-Hochberg a value of *p* < 0.05.

### Electrophysiology

For brain slice preparation, mice were anesthetized with isoflurane and decapitated. The brains were rapidly removed and chilled in ice-cold sucrose solution (pH 7.3) containing (mM) 76 NaCl, 25 NaHCO_3_, 25 glucose, 75 sucrose, 2.5 KCl, 1.25 NaH_2_PO_4_, 0.5 CaCl_2_, and 7 MgSO_4_. Acute brain slices (300 μm) including the medial prefrontal cortex (mPFC) were prepared using a vibratome (VT-1200s, Leica). Slices were then incubated in warm (32-35°C) sucrose solution for 30 min, then transferred to warm (32-34°C) artificial cerebrospinal fluid (aCSF, pH 7.3, 315 mOsm) composed of (mM) 125 NaCl, 26 NaHCO_3_, 2.5 KCl, 1.25 NaH_2_PO_4_, 1MgSO_4_, 20 glucose, 2 CaCl_2_, 0.4 ascorbic acid, 2 pyruvic acid, and 4 L-(+)-lactic acid. Slices were allowed to cool to room temperature. All solutions were continuously bubbled with 95 % O_2_ and 5 % CO_2_. For whole-cell recordings, slices were transferred to a submersion chamber on an upright microscope (Zeiss AxioExaminer, Objectives: 5x, 0.16 NA and 40x, 1.0 NA) fitted for infrared differential interference contrast (DIC). Slices were continuously superfused (2-4 ml/min) with warm oxygenated aCSF (32-34°C). Neurons were visualized with a digital camera (Sensicam QE; Cooke) using transmitted light. Prelimbic mPFC was identified based on the shape and location of the corpus callosum.

For intrinsic excitability measurements, glass recording electrodes (2-4 MΩ) were filled with an internal solution (pH 7.3, 295 mOsm) containing (mM) 2.7 KCl, 120 KMeSO_4_, 9 HEPES, 0.18 EGTA, 4 MgATP, 0.3 NaGTP, and 20 phosphocreatine (Na). Electrophysiological recordings for measuring membrane properties and intrinsic excitability were performed in the presence of the following blockers of glutamate and GABA receptors: 5 μM 2,3-Dioxo-6-nitro-1,2,3,4-tetrahydrobenzo[*f*]quinoxaline-7-sulfonamide disodium salt (NBQX; AMPA receptor antagonist), 5 μM (*RS*)-3-(2-Carboxypiperazin-4-yl)-propyl-1-phosphonic acid (CPP; NMDA receptor antagonist), and 10 μM 6-Imino-3-(4-methoxyphenyl)-1(6*H*)-pyridazinebutanoic acid hydrobromide (SR95531; GABA_A_ receptor antagonist, all from Tocris). The resting membrane potential was measured after whole-cell configuration was achieved. Neurons exhibiting a resting membrane potential greater than -60 mV were excluded from recordings. The input resistance was determined by measuring the voltage change in response to a 1-s-long -100 pA hyperpolarizing current. The current-spike frequency relationship was measured with a range of depolarizing current steps presented in a pseudorandom order (1 s long, 40 pA increments, 5 s interstimulus intervals). For each current intensity, the total number of action potentials exceeding 0 mV generated during each step was measured and averaged across the three trials. The rheobase was determined by first probing the response of the neuron with 1-s-long depolarizing steps (5 s interstimulus intervals) to define a small range of current steps that bounded the rheobase. The response of the neurons was then tested within this range using 1-s-long depolarizing steps with 1 pA increments. All signals were low-pass filtered at 10 kHz and sampled at 20-100 kHz.

Data analysis was performed in Igor Pro (WaveMetrics). IT- and PT-type neurons were categorized based on their sag amplitude (determined by the difference between peak hyperpolarized potential and steady-state potential) and sag decay slope (average of differential values of voltage trace from peak hyperpolarized potential point to 100 ms from the peak point). Cells showing <1 mV sag amplitude and <0.07 mV/ms sag decay slope were categorized as IT neurons (61 out of 115 cells). Cells showing >1 mV sag amplitude and >0.07 mV/ms sag decay slope were counted as PT neurons (43 out of 115 cells). The remaining cells, which did not meet both criteria, were categorized as Others (11 out of 115 cells) and excluded from further analysis. The access resistance (Ra) of all cells was <30 MΩ at the start of recording. 5x differential interference contrast (DIC) images were acquired and saved for all cells recorded to confirm the locations of cells in layer 5 of the mPFC. For the electrophysiological experiments using the Cx3cr1^CreER^ mice and the adult THC treatment, the following set of hardware and software was used. The electrophysiological rig consisted of Axioscope2 FS plus microscope (Zeiss) and pco.edge 4.2 bi digital camera (PCO), and pCLAMP software (Molecular Devices) was used for data acquisition and analysis. The experimenter was blinded to the mouse information during recording experiments and raw data analysis.

### Behavioral tests

Behavioral tests were conducted on male and female mice housed on a reversed 12-h light/dark cycle starting at P72 or P112. All tests were conducted during the dark period of the cycle. After 1-week habituation in reversed 12-h light/dark cycle, mice were subjected to behavior tests in the following order: olfactory habituation/dis-habituation test, open field test, elevated plus maze test, novel object recognition test, novel place recognition test, three-chamber social approach test, and 5-trial social memory test. The interval between different behavioral tests was at least 24 h. Each apparatus was cleaned with 70% ethanol between individual animals to control for odor cues.

#### Olfactory habituation/dis-habituation test

The olfactory habituation/dis-habituation test was designed to determine whether a mouse can remember and distinguish between odors ^91^. The odors were presented on a suspended cotton swab to the test mouse in a clean cage with fresh shavings. Each mouse was tested during three consecutive 2-min periods for each odor with 2-min intervals. The time that the mouse smelled the swab was recorded. After three trials (first block) with the odorless swab (water), the same three trials were repeated (second block) with the swab dipped in diluted vanilla flavor (1:33; GEL SPICE, Bayonne, NJ). After the second block of trials, the swab was dipped in diluted banana flavor (1:100; McCormick, Hunt Valley, MD) and presented to the animals (third block). In general, mice habituated quickly to odors, and sniffing time declined when the mouse was exposed several times to the same odor. When a new odor was introduced, there was an increase in sniffing time, indicating that the animal could discriminate between odors.

#### Open field test

The open field test was performed according to our previously published methods with minor modification^92, 93^. Locomotor activity was assessed over a 30-min period in 40 × 40 cm activity chambers with built-in infrared beams (PAS system, San Diego Instruments Inc., San Diego, CA, USA). Horizontal and vertical locomotor activities in the center or along the walls (periphery) of the chamber and rearing were automatically recorded as beam breaks.

#### Elevated plus maze test

The elevated plus maze test was conducted to evaluate anxiety-like behaviors using our previously published protocol^92^. A mouse was placed in the intersection (middle) of the arms in the plus maze (San Diego Instruments, Inc.), and observed and videotaped for 5 min. The number of entries into the closed and open arms, as well as the time spent in the closed vs open arms, were recorded.

#### Novel object recognition test and novel place recognition test

Mice were placed into a Plexiglas open-field arena (20 × 40 × 22 cm). Each mouse was individually habituated to the arena with 10 min of exploration in the absence of objects each day for 3 consecutive days (habituation). On day 3, an hour after habituation, two objects were secured to the floor of the arena at adjacent corners, and each animal was allowed to explore the arena for 10 min (training). The objects were identical in shape, color, and size. For novel object recognition, 15 min after the training, each mouse was placed back into the arena but with one of the familiar objects used during training replaced by a novel object. Each mouse was allowed to explore for 5 min and the time spent exploring each object was recorded (retention). The novel object was similar in size to the familiar object, but different in shape and color. Ratios comparing the amount of time spent exploring either of the two objects (training session) or the novel object (retention session), both over the total time spent exploring both objects, were used to measure object recognition memory. Similarly, individual mice were tested in the spatial component via novel place recognition test. During the retention session, one of the two familiar objects now occupied a new location in the arena with respect to the previous trial (defined as novel location). Each mouse was allowed to explore for 5 min and the time spent exploring each location was recorded. A ratio of the amount of time spent exploring the novel location over the total time spent exploring both locations was used to measure spatial recognition memory. Exploration was defined as sniffing the object.

#### Three-chamber social approach test

The three-chamber social interaction test was performed according to our previously published methods with minor modification^94^. The assay consisted of four sessions. The first session began with 10 min habituation in the center chamber followed by a second 10 min session where the subject mouse could freely explore all three chambers including two side chambers, each with a plastic cage for habituation. In the third session, the mouse was gently confined to the center chamber while a novel intruder control mouse (stranger 1) was placed in one plastic cage in one side chamber, and an inanimate mouse-like object was placed in the other. The subject mouse was allowed to freely explore all three chambers for 10 min. Before the last session, the subject mouse was gently guided to the center chamber while the inanimate mouse-like object was replaced with another novel intruder control mouse (stranger 2) to assess for social novelty recognition. The subject mouse freely explored all three chambers for 10 min. The positions of the inanimate cage and stranger 1 cage were alternated between tests to prevent side preference. The plastic cages used in the tests allowed substantial olfactory, auditory, visual, and tactile contact between subject mice and stranger mice. Individual movement (i.e., social sniffing) of the subject mice was analyzed by a researcher who was blinded to the group assignment. The heat maps were generated by Ethovision XT 11.0 (Noldus, Leesburg, VA).

#### 5-trial social memory test

For the 5-trial recognition test with social cues, we transferred mice from group to individual housing for 10 d before testing to permit establishment of a home cage territory. The test began when a stimulus female mouse was introduced into the home cage of each male mouse for a 1 min confrontation. Age-matched ovariectomized female C57BL/6 mice from Jackson Laboratories (JAX; Bar Harbor, ME, USA) were used as stimulus mice. At the end of the 1 min trial, the stimulus mouse was removed and returned to an individual holding cage. This sequence was repeated for four trials with 10 min inter-trial intervals. The same stimulus mouse was introduced to the same male resident in all four trials. In a fifth dishabituation trial, we introduced a different stimulus mouse to the resident male mouse. Behavior was recorded and interaction time (time for nosing, anogenital sniffing, and close following and pursuit) was scored. Aggressive posturing and sexual behaviors (e.g. mounting) were not included in the calculation of social interaction time.

### Statistical analyses

Behavioral, biochemical, and morphological phenotyping data were analyzed by two-way mixed analyses of variance (ANOVA) to identify a possible synergistic effect (source of variation due to interaction) with the group (16p11dup and wild type littermate control) and treatment (THC and vehicle) as independent factors. Tukey’s post-hoc test was used to determine statistically significant differences between each group. Two-tailed analyses of unpaired Student *t*-tests were also used for data analysis in comparing THC and vehicle treatment or 16p11dup and wild type littermate controls. These statistical analyses were conducted by using GraphPad Prism 7 (GraphPad Software, La Jolla, CA, USA). A value of *p* < 0.05 was considered statistically significant. All data are presented as means ± standard error of the mean (s.e.m.).

### Data Availability

The datasets generated and analyzed during the current study are available from the corresponding author on reasonable request, together with source codes. Data are also available at the NCBI Gene Expression Omnibus under accession number GSE###### (Submission in process).

## ACKNOWLEDGEMENTS

We thank Ms. Joi Haskins for critical reading of the manuscript. We thank Dr. Ken Mackie for providing us with the mouse model with *Cnr2* deletion. We thank Dr. Kun Yang for providing us with critical advice for the RNA-seq data analysis. We thank the Johns Hopkins University School of Medicine Experimental and Computational Genomics Core, Flow Cytometry Core Facility, Microscopy Core Facility, and Behavioral Core Facility. Fig 1a, Fig 4a, Fig 5b, c, d, Fig 6b, Fig 7a, and Fig 8a were created with a graphical software provided by Biorender.com. This work was supported by grants from the National Institute of Health Awards DA041208 (A.K., M.V.P.), MH094268 (A.S., M.V.P, A.K.), AG065168 (A.K.), and AT010984 (X.Z.), MH128765 (J.K., A.K.) as well as foundation grants from Kanae (Y.H.), NARSAD Young Investigator Grant from the Brain & Behavior Research Foundation (J.K.), and basic research program through Korea Brain Research Institute, 23-BR-04-04 (J.K.).

## AUTHOR CONTRIBUTIONS

Y.H., M.V.P., and A.K. designed the study. Y.H. performed cellular, behavioral, and immunohistochemical experiments, and contributed to data analysis for all experiments. Y.J., X.Z., Y.M., F.X., S.M., and M.O. assisted Y.H. in conducting cellular, behavioral, and histochemical assays. J.K. and Y.H. performed electrophysiology experiments and analyzed the data with S.P.B. G.U. analyzed gene expression data from RNA-sequencing. B.L. provided *Cnr1^flox/flox^* mice and contributed to data interpretation. A.S. contributed to making conceptual framework and data interpretation. A.K. and M.V.P. contributed to the concept or design of the work, and drafted the manuscript with Y.H. All authors have approved the final manuscript.

## COMPETING FINANCIAL INTERESTS

The authors declare no competing financial interests.

## Notes

### Competing Interest Statement

The authors have declared no competing interest.

