## Supplemental Figure and Table for "Microglial cannabinoid receptor type 1 mediates social memory deficits produced by adolescent THC exposure and 16p11.2 duplication"

### Supplementary Fig. 1

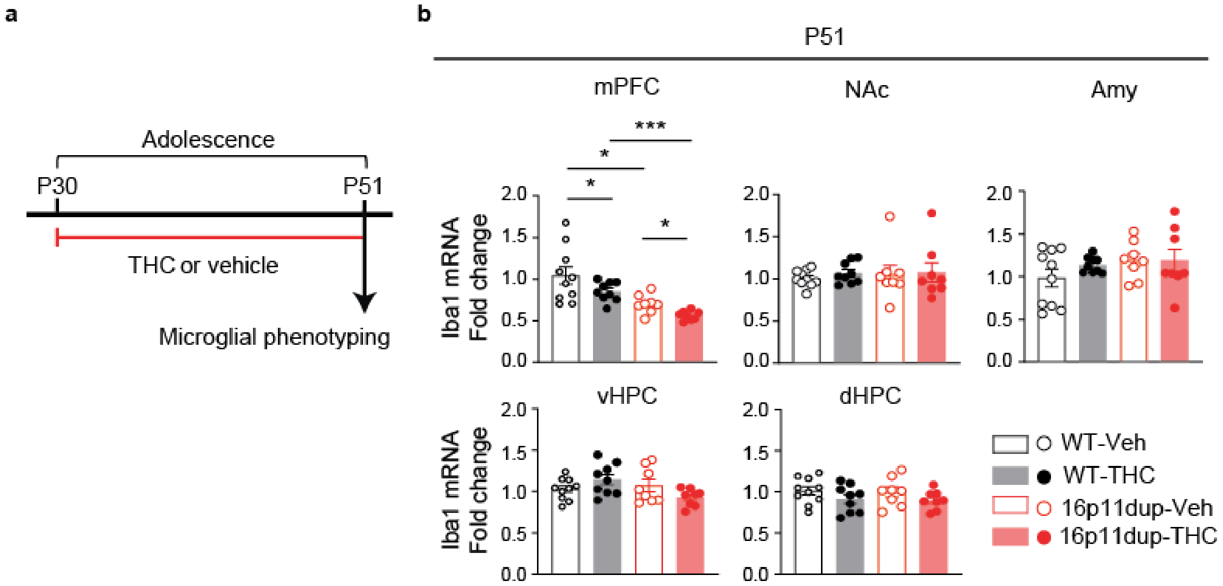

**Supplementary Fig. 1 Iba1 mRNA reduction in the mPFC produced by adolescent THC treatment and 16p11dup. a,** Schematic diagram of the adolescent THC treatment protocol. 16p11dup mice and wild type littermate controls (WT) were treated with THC (s.c., 8mg/kg) or vehicle (Veh) during adolescence (P30-P51), followed by microglial phenotyping at P51 upon completion of THC treatment. **b,** Relative mRNA expression level of Iba1 in brain regions involved in the medial prefrontal cortex (mPFC), ventral hippocampus (vHPC), dorsal hippocampus (dHPC), nucleus accumbens (NAc), and amygdala (Amy) at P51. WT-Veh ($n$ = 10 mice), WT-THC ($n$ = 9 mice), 16p11dup-Veh ($n$ = 8 mice), and 16p11dup-THC ($n$ = 8 mice). (**b**) ***$p < 0.001$, *$p < 0.05$, determined by two-way ANOVA with post hoc Tukey test. Data are presented as the mean ± s.e.m.

**Supplementary Fig. 2**

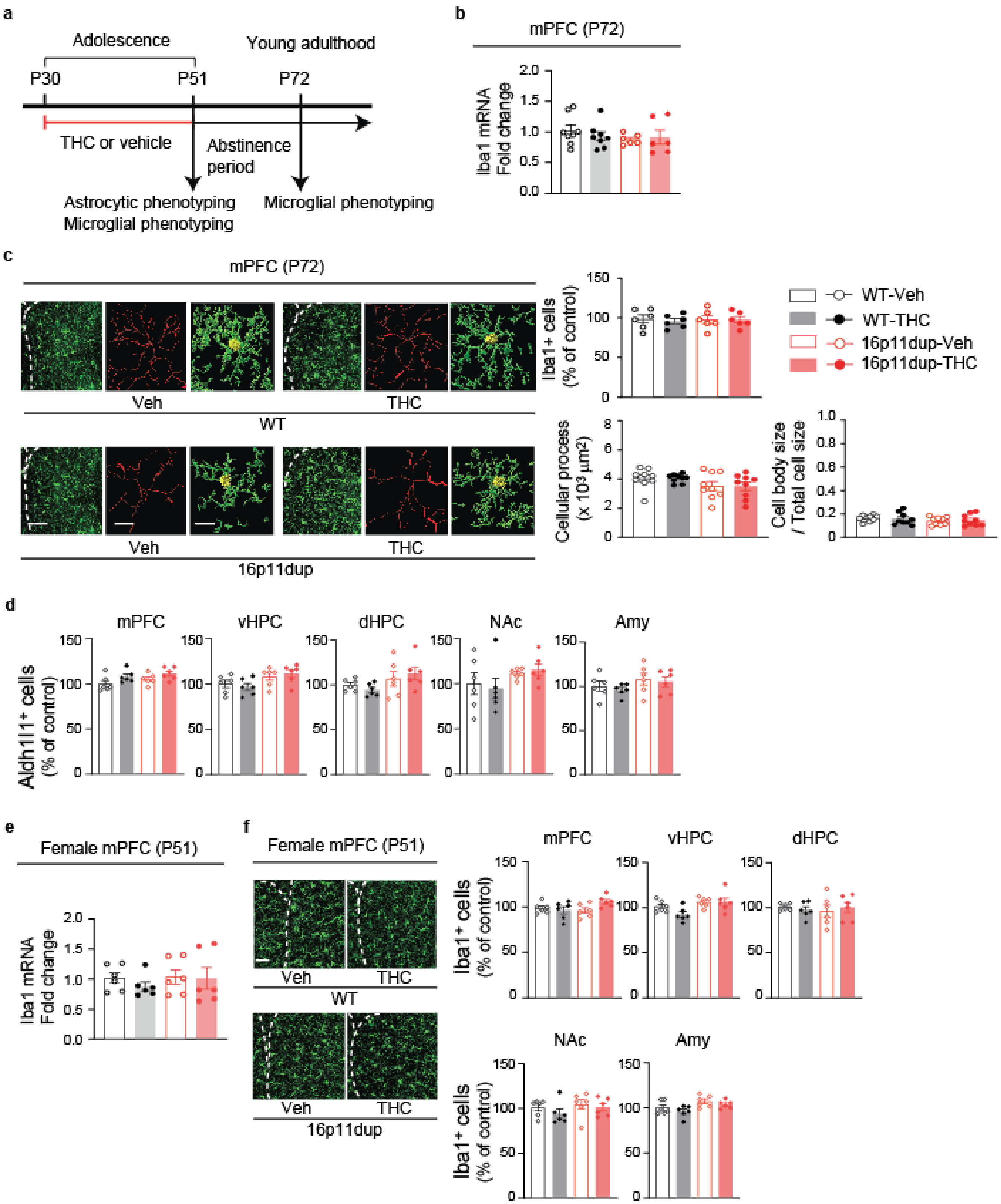

**Supplementary Fig. 2 No effect of adolescent THC treatment and 16p11dup on microglial phenotypes in adulthood, the number of astrocytes, and female microglial phenotypes. a**, Schematic diagram of the adolescent THC treatment protocol. 16p11dup mice and wild type littermate controls (WT) were treated with THC (s.c., 8mg/kg) or vehicle (Veh) during adolescence (P30-P51), followed by astrocytic phenotyping and microglial phenotyping at P51 upon completion of THC treatment and microglial phenotype at P72 after a 3-week abstinence period. **b**, Relative mRNA expression level of Iba1 in the medial prefrontal cortex (mPFC) at P72. WT-Veh ($n$ = 8 mice), WT-THC ($n$ = 8 mice), 16p11dup-Veh ($n$ = 6 mice), 16p11dup-THC ($n$ = 6 mice). **c,** Immunohistochemistry of Iba1 (green) in the mPFC at P72. (Left) Representative images of the mPFC, representative tracing images (red) along with images of cellular processes (green) and cell bodies (yellow) of Iba1$^{+}$ cells. Scale bar, 50 μm (left) and 10 μm (middle and right). (Top right) The number of Iba1$^{+}$ cells in the mPFC, presented as % of control. ($n$ = 6 slices in 3 mice per condition). (Bottom right) Quantification of cellular process area (left) and the ratio of cell body size to total cell size (right) of Iba1$^{+}$ cells. (n = 9 cells in 3 mice per condition). **d,** Immunohistochemistry of Aldh1l1 in the mPFC, ventral hippocampus (vHPC), dorsal hippocampus (dHPC), nucleus accumbens (NAc), and amygdala (Amy) at P51. The number of Aldh1l1$^{+}$ cells in these brain regions, presented as % of control. ($n$ = 6 slices in 3 mice per condition). **e**, Relative mRNA expression level of Iba1 in the mPFC of female mice at P51. ($n$ = 6 slices in 3 mice per condition). **f,** Immunohistochemistry of Iba1 (green) in the mPFC, vHPC, dHPC, NAc, and Amy at P51. (Left) Representative images of the mPFC. Scale bar, 50 μm. (Right) The number of Iba1$^{+}$ cells in these brain regions, presented as % of control ($n$ = 6 slices in 3 mice per group). Data are presented as the mean ± s.e.m.

**Supplementary Fig. 3**

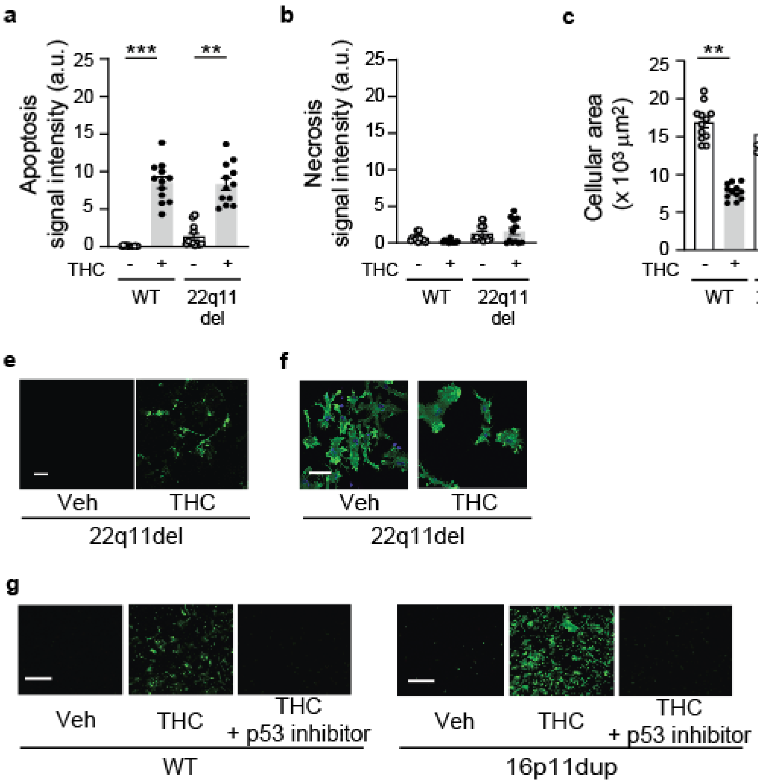

**Supplementary Fig. 3 22q11del did not enhance THC-induced microglial phenotypes and p53 inhibition blocked microglial apoptosis produced by THC treatment and 16p11dup. a**, Apoptosis assay of primary microglia cultures produced from WT and 22q11del male mice. Quantification of signal intensity of apoptosis marker apopxin (*n* = 12 fields in 3 mice per condition). **b,** Necrosis assay of primary microglia cultures produced from WT and 22q11del male mice. Quantification of signal intensity of necrosis marker 7-AAD (*n* = 12 fields in 3 mice per condition). **c,** Quantification of cellular area of phalloidin-stained microglia cultures produced from WT and 22q11del male mice (*n* = 12 fields in 3 mice per condition). **d**, Quantification of cellular process number of phalloidin-stained microglia cultures produced from WT and 22q11del male mice (*n* = 12 fields in 3 mice per condition). **e,** Representative images of microglia cell cultures produced from WT and 22q11del male mice in apoptosis and necrosis assays. Apopxin (green) and 7-AAD (red) are shown. Scale bar, 50 μm. **f,** Immunohistochemistry with antibody against phalloidin (green) of primary microglia cultures produced from WT and 22q11del male mice. Scale bar, 25 μm. **g**, Representative images of primary microglia cultures treated by THC or vehicle together with and without pifithrin-α, an inhibitor of p53 transcriptional activity, in apoptosis and necrosis assays. Apopxin (green) and 7-AAD (red) were shown. Scale bar, 100 μm. (**a, c**, **d**) ***$p < 0.001$, **$p < 0.01$, *$p < 0.05$, determined by two-way ANOVA with post hoc Tukey test. Data are presented as the mean ± s.e.m.

**Supplementary Fig. 4**

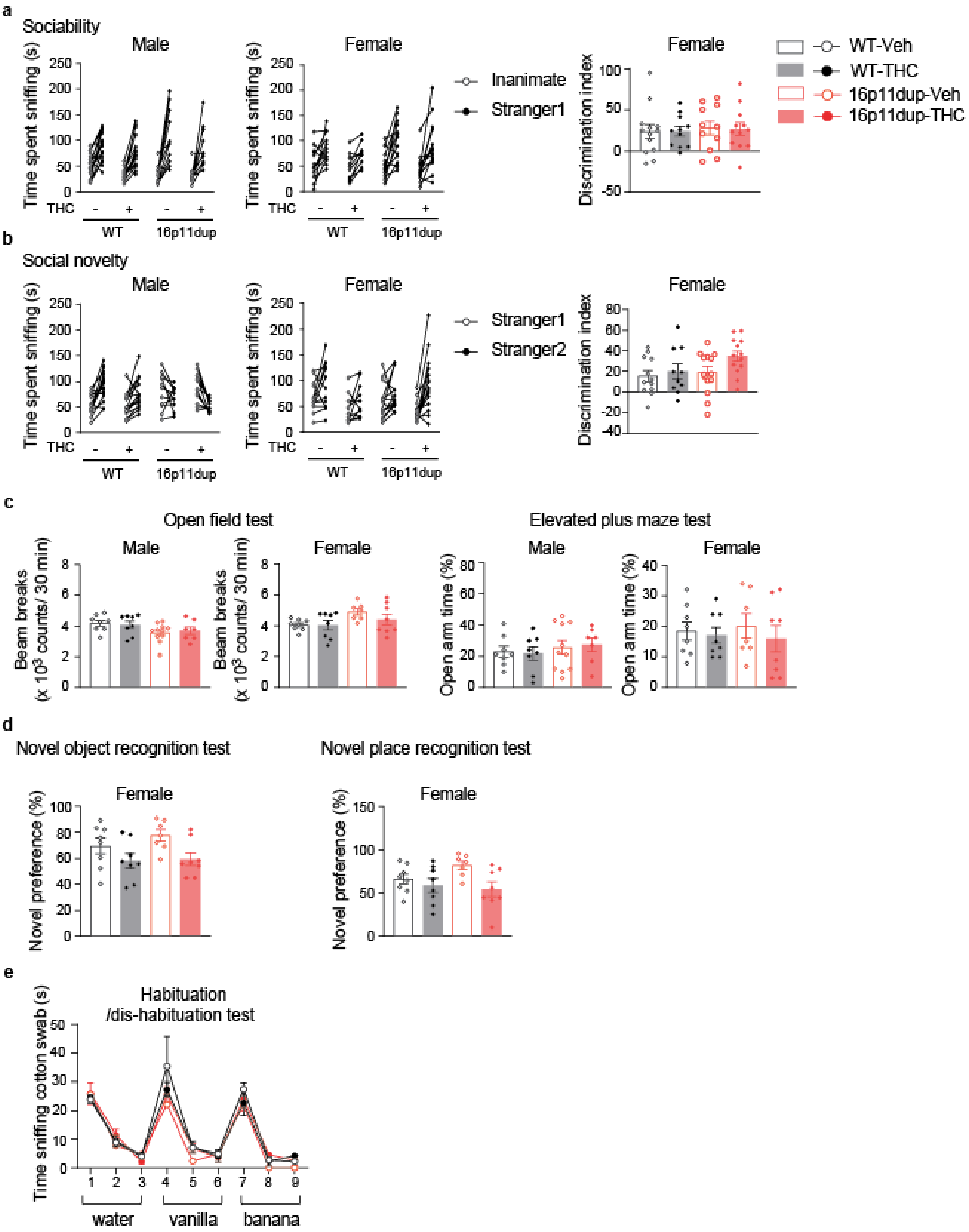

**Supplementary Fig. 4 No effect of adolescent THC treatment and 16p11 on spontaneous locomotion, anxiety, olfaction, or object and place recognition memory. a**, (Left) Time the mice spent sniffing an Inanimate vs. a Stranger 1 mouse in male and female mice. (Right) Sociability of WT and 16p11dup female mice with adolescent THC or Veh treatment as indicated by discrimination index ([Stranger 1 – Inanimate mice sniffing time] / [Stranger 1 + Inanimate mice sniffing time] × 100 (%)). **b**, (Left) Time the mice spent sniffing a Stranger 1 mouse vs. a Stranger 2 mouse in male and female mice. (Right) Preference of social novelty of WT and 16p11dup female mice with adolescent THC or Veh treatment as indicated by discrimination index ([Stranger 2 – Stranger 1 mice sniffing time] / [Stranger 2 + Stranger 1 mice sniffing time] × 100 (%)). (**a**, **b**) Male: WT-Veh ($n$ = 17 mice), WT-THC ($n$ = 15 mice), 16p11dup-Veh ($n$ = 11 mice), 16p11dup-THC ($n$ = 11 mice); Female: WT-Veh ($n$ = 10 mice), WT-THC ($n$ = 10 mice), 16p11dup-Veh ($n$ = 15 mice), 16p11dup-THC ($n$ = 15 mice). **c**, (Left) Spontaneous locomotion activity of 16p11dup and wild type littermate (WT) male and female mice with adolescent THC or Veh treatment as indicated by total beam breaks in the open field test. (Right) Anxiety-like phenotype of WT and 16p11dup male and female mice with adolescent THC or Veh treatment as indicated by percentage of time spent in the open arms in the elevated plus maze test. **d**, (Left) Object recognition memory of WT and 16p11dup female mice with adolescent THC or Veh treatment as indicated by preference of novel object in the novel object recognition test. (Right) Place recognition memory of WT and 16p11dup female mice with adolescent THC or Veh treatment as indicated by preference of novel place in the novel place recognition test. (**c**, **d**) Male: WT-Veh ($n$ = 8 mice), WT-THC ($n$ = 8 mice), 16p11dup-Veh ($n$ = 11 mice), 16p11dup-THC ($n$ = 7 mice), Female: WT-Veh ($n$ = 8 mice), WT-THC ($n$ = 8 mice), 16p11dup-Veh ($n$ = 7 mice), 16p11dup-THC ($n$ = 8 mice). **e**, Olfactory discrimination in WT and 16p11dup mice with adolescent THC or Veh treatment as indicated by time the mice spent sniffing cotton swab in habituation/dis-habituation test. WT-Veh ($n$ = 17 mice),

WT-THC (*n* = 15 mice), 16p11dup-Veh (*n* = 11 mice), 16p11dup-THC (*n* = 11 mice). Data are presented as the mean ± s.e.m.

### Supplementary Fig. 5

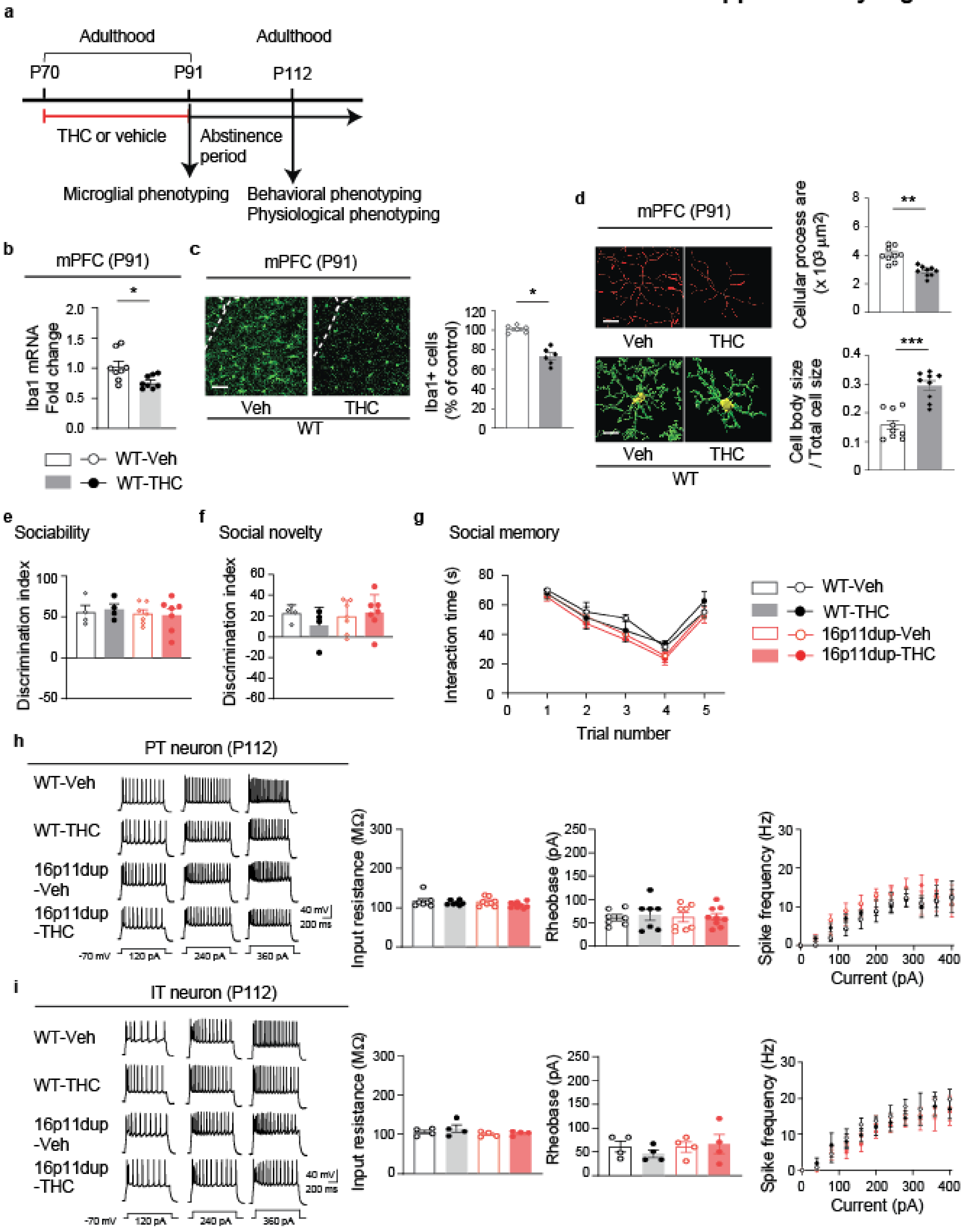

**Supplementary Fig. 5 Adult THC treatment induced microglia reduction and morphological changes, but no effect on social recognition, memory and intrinsic excitability in mPFC. a**, Schematic diagram of the adult THC treatment protocol. Wild type male mice (WT) were treated with THC (s.c., 8mg/kg) or vehicle (Veh) during adulthood (P70-P91), followed by microglial phenotyping at P91 upon completion of THC treatment and behavioral and physiological assays after a 3-week abstinence period. **b**, Relative mRNA expression level of Iba1 in the mPFC at P91. (*n* = 8 mice per condition). **c**, Immunohistochemistry of Iba1 (green) in the mPFC at P91. (Left) Representative images of the mPFC. Scale bar, 50 μm. (Right) The number of Iba1$^{+}$ cells in the mPFC, presented as % of control (*n* = 6 slices in 3 mice per condition). **d**, Microglial morphology analysis of individual Iba1$^{+}$ cells in the mPFC. (Top left) Representative tracing images (red) of Iba1$^{+}$ cells. Scale bar, 10 μm. (Top right) Quantification of cellular process area of Iba1$^{+}$ cells. (*n* = 9 cells in 3 mice per condition). (Bottom left) Representative images of cellular processes (green) and cell bodies (yellow) of Iba1$^{+}$ cells. Scale bar, 10 μm. (Bottom right) Quantification of the ratio of cell body size to total cell size of Iba1$^{+}$ cells (n = 9 cells in 3 mice per condition). **e,** Sociability of WT and 16p11dup mice with adult THC or Veh treatment as indicated by discrimination index ([Stranger1 – Inanimate mice sniffing time] / [Stranger 1 + Inanimate mice sniffing time] × 100 (%)). **f**, Preference of social novelty of WT and 16p11dup mice with adult THC or Veh treatment as indicated by discrimination index ([Stranger 2 – Stranger 1 mice sniffing time] / [Stranger 2 + Stranger 1 mice sniffing time] × 100 (%)). **g**, Interaction time of WT and 16p11dup mice receiving adult THC or Veh treatment with ovariectomized female mice. (**e, f, g**) WT-Veh (*n* = 4 mice), WT-THC (*n* = 4 mice), 16p11dup-Veh (*n* = 7 mice), 16p11dup-THC (*n* = 7 mice). **h,** (Left) Representative voltage traces recorded from PT neurons in response to current step injections in the presence of blockers of AMPA, NMDA, and GABAa receptors. (Right) The intrinsic excitability assessed by measurement of input resistance (left), rheobase (middle), and spike frequency (right). WT-

Veh (*n* = 7 cells in 4 mice), WT-THC (*n* = 7 cells in 3 mice), 16p11dup-Veh (*n* = 8 cells in 3 mice), and 16p11dup-THC (*n* = 9 cells in 3 mice). **i,** (Left) Representative voltage traces recorded from IT neurons in response to current step injections in the presence of blockers of AMPA, NMDA, and GABAa receptors.. (Right) The intrinsic excitability assessed by measurement of input resistance (left), rheobase (middle), and spike frequency (right). WT-Veh (*n* = 4 cells in 2 mice), WT-THC (*n* = 4 cells in 2 mice), 16p11dup-Veh (*n* = 4 cells in 2 mice), and 16p11dup-THC (*n* = 4 cells in 3 mice). (**b, c, d**) $***p < 0.001$, $**p < 0.01$, $*p < 0.05$, determined by Student *t*-tests. Data are presented as the mean ± s.e.m.

### Supplementary Fig. 6

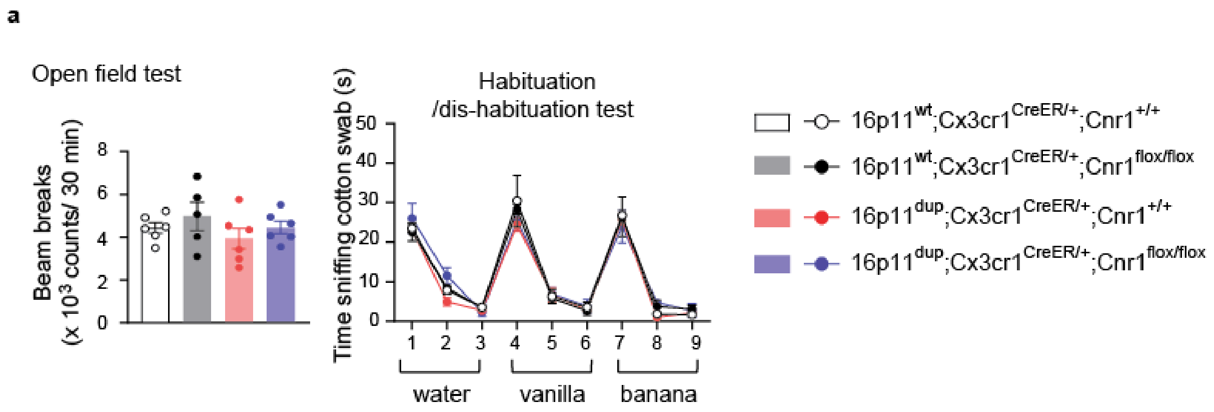

**Supplementary Fig. 6 No effects of microglial genetic deletion of *Cnr1* on locomotion and olfaction in the male mice treated with THC during adolescence. a**, (Left) Spontaneous locomotion activity of four groups of mice which include 16p11$^{dup}$;*Cx3cr1*$^{CreER/+}$; *Cnr1*$^{+/+}$, 16p11$^{dup}$;*Cx3cr1*$^{CreER/+}$;*Cnr1*$^{flox/flox}$, wild type littermate (16p11$^{wt}$);*Cx3cr1*$^{CreER/+}$;*Cnr1*$^{+/+}$, and 16p11$^{wt}$ ;*Cx3cr1*$^{CreER/+}$;*Cnr1*$^{flox/flox}$ mice with adolescent THC treatment as indicated by total beam breaks in the open field test. (Right) Olfactory discrimination of these four groups of mice with adolescent THC treatment as indicated by time that the mice spent sniffing cotton swab in habituation/dis-habituation test. 16p11$^{dup}$;*Cx3cr1*$^{CreER/+}$; *Cnr1*$^{+/+}$ ($n$ = 6 mice), and 16p11$^{dup}$;*Cx3cr1*$^{CreER/+}$;*Cnr1*$^{flox/flox}$ ($n$ = 6 mice), 16p11$^{wt}$;*Cx3cr1*$^{CreER/+}$;*Cnr1*$^{+/+}$ ($n$ = 6 mice), and 16p11$^{wt}$ ;*Cx3cr1*$^{CreER/+}$;*Cnr1*$^{flox/flox}$ ($n$ = 5 mice). Data are presented as the mean ± s.e.m.

**Supplementary Table 1: 16p11dup with THC enriched pathway and upstream regulator**

| Reactome pathways name | The number of genes in Reference list | The number of genes in analysed list | raw P value | FDR |
|---|---|---|---|---|
| Cdc20:Phospho-APC/C mediated degradation of Cyclin A | 71 | 13 | 8.48E-06 | 3.50E-03 |
| APC:Cdc20 mediated degradation of cell cycle proteins prior to satisfation of the cell cycle checkpoint | 72 | 13 | 9.69E-06 | 3.20E-03 |
| APC/C:Cdc20 mediated degradation of mitotic proteins | 74 | 13 | 1.26E-05 | 3.45E-03 |
| Activation of APC/C and APC/C:Cdc20 mediated degradation of mitotic proteins | 75 | 13 | 1.43E-05 | 3.36E-03 |
| TCR signaling | 105 | 15 | 2.52E-05 | 4.61E-03 |
| APC/C-mediated degradation of cell cycle proteins | 82 | 13 | 3.30E-05 | 4.53E-03 |
| Regulation of mitotic cell cycle | 82 | 13 | 3.30E-05 | 4.94E-03 |
| Regulation of mRNA stability by proteins that bind AU-rich elements | 85 | 13 | 4.60E-05 | 5.84E-03 |
| Asymmetric localization of PCP proteins | 61 | 11 | 4.70E-05 | 5.17E-03 |
| Transcriptional regulation by RUNX2 | 61 | 11 | 4.70E-05 | 5.54E-03 |
| Downstream TCR signaling | 86 | 13 | 5.13E-05 | 5.29E-03 |
| Autodegradation of Cdh1 by Cdh1:APC/C | 62 | 11 | 5.36E-05 | 5.20E-03 |
| Stabilization of p53 | 54 | 10 | 8.51E-05 | 7.79E-03 |
| APC/C:Cdc20 mediated degradation of Securin | 66 | 11 | 8.84E-05 | 7.68E-03 |
| Degradation of GLI1 by the proteasome | 55 | 10 | 9.73E-05 | 8.03E-03 |
| C-type lectin receptors (CLRs) | 107 | 14 | 1.09E-04 | 8.57E-03 |
| GLI3 is processed to GLI3R by the proteasome | 56 | 10 | 1.11E-04 | 8.33E-03 |
| NIK-->noncanonical NF-kB signaling | 57 | 10 | 1.26E-04 | 9.07E-03 |
| Dectin-1 mediated noncanonical NF-kB signaling | 57 | 10 | 1.26E-04 | 8.69E-03 |
| CDT1 association with the CDC6:ORC:origin complex | 58 | 10 | 1.44E-04 | 9.47E-03 |
| Post-translational protein modification | 1233 | 73 | 1.45E-04 | 9.20E-03 |
| CDK-mediated phosphorylation and removal of Cdc6 | 71 | 11 | 1.58E-04 | 9.63E-03 |
| APC/C:Cdh1 mediated degradation of Cdc20 and other APC/C:Cdh1 targeted proteins in late mitosis/early G1 | 71 | 11 | 1.58E-04 | 9.29E-03 |
| MAPK6/MAPK4 signaling | 73 | 11 | 1.96E-04 | 1.12E-02 |
| FCERI mediated NF-kB activation | 73 | 11 | 1.96E-04 | 1.08E-02 |
| The role of GTSE1 in G2/M progression after G2 checkpoint | 73 | 11 | 1.96E-04 | 1.04E-02 |
| PCP/CE pathway | 87 | 12 | 2.14E-04 | 1.07E-02 |
| CLEC7A (Dectin-1) signaling | 87 | 12 | 2.14E-04 | 1.04E-02 |
| Ubiquitin-dependent degradation of Cyclin D | 50 | 9 | 2.31E-04 | 1.09E-02 |
| Autodegradation of the E3 ubiquitin ligase COP1 | 50 | 9 | 2.31E-04 | 1.06E-02 |
| Regulation of RUNX2 expression and activity | 50 | 9 | 2.31E-04 | 1.03E-02 |
| Hedgehog ligand biogenesis | 62 | 10 | 2.33E-04 | 1.01E-02 |
| Axon guidance | 258 | 23 | 2.41E-04 | 1.02E-02 |
| Switching of origins to a post-replicative state | 89 | 12 | 2.59E-04 | 1.04E-02 |
| p53-Dependent G1 DNA Damage Response | 63 | 10 | 2.61E-04 | 1.00E-02 |
| p53-Dependent G1/S DNA damage checkpoint | 63 | 10 | 2.61E-04 | 1.03E-02 |
| Ubiquitin Mediated Degradation of Phosphorylated Cdc25A | 51 | 9 | 2.63E-04 | 9.43E-03 |
| p53-Independent DNA Damage Response | 51 | 9 | 2.63E-04 | 9.64E-03 |
| p53-Independent G1/S DNA damage checkpoint | 51 | 9 | 2.63E-04 | 9.86E-03 |
| Fc epsilon receptor (FCERI) signaling | 118 | 14 | 2.75E-04 | 9.66E-03 |
| Oxygen-dependent proline hydroxylation of Hypoxia-inducible Factor Alpha | 64 | 10 | 2.93E-04 | 1.01E-02 |
| Activation of NF-kappaB in B cells | 64 | 10 | 2.93E-04 | 9.85E-03 |
| G1/S DNA Damage Checkpoints | 65 | 10 | 3.27E-04 | 1.08E-02 |
| Regulation of RUNX3 expression and activity | 53 | 9 | 3.39E-04 | 1.10E-02 |
| Degradation of AXIN | 54 | 9 | 3.83E-04 | 1.21E-02 |
| FBXL7 down-regulates AURKA during mitotic entry and in early mitosis | 54 | 9 | 3.83E-04 | 1.19E-02 |
| Assembly of the pre-replicative complex | 67 | 10 | 4.06E-04 | 1.24E-02 |
| Degradation of DVL | 55 | 9 | 4.32E-04 | 1.29E-02 |
| AUF1 (hnRNP D0) binds and destabilizes mRNA | 55 | 9 | 4.32E-04 | 1.27E-02 |
| TNFR2 non-canonical NF-kB pathway | 96 | 12 | 4.83E-04 | 1.35E-02 |
| ABC-family proteins mediated transport | 96 | 12 | 4.83E-04 | 1.37E-02 |
| Cellular response to hypoxia | 69 | 10 | 5.00E-04 | 1.37E-02 |
| DNA Replication Pre-Initiation | 83 | 11 | 5.28E-04 | 1.43E-02 |
| Orc1 removal from chromatin | 70 | 10 | 5.54E-04 | 1.47E-02 |
| Antigen processing: Ubiquitination & Proteasome degradation | 291 | 24 | 5.61E-04 | 1.47E-02 |
| Innate Immune System | 929 | 56 | 5.71E-04 | 1.47E-02 |
| Cellular responses to stress | 404 | 30 | 5.78E-04 | 1.47E-02 |
| Metabolism of polyamines | 58 | 9 | 6.10E-04 | 1.52E-02 |
| M Phase | 353 | 27 | 6.94E-04 | 1.68E-02 |
| SCF(Skp2)-mediated degradation of p27/p21 | 60 | 9 | 7.58E-04 | 1.81E-02 |
| Cross-presentation of soluble exogenous antigens (endosomes) | 48 | 8 | 8.02E-04 | 1.89E-02 |
| G2/M Transition | 182 | 17 | 8.34E-04 | 1.94E-02 |
| Mitotic G2-G2/M phases | 184 | 17 | 9.33E-04 | 2.14E-02 |
| Downstream signaling events of B Cell Receptor (BCR) | 76 | 10 | 9.83E-04 | 2.22E-02 |
| Regulation of ornithine decarboxylase (ODC) | 50 | 8 | 1.02E-03 | 2.27E-02 |
| L1CAM interactions | 64 | 9 | 1.14E-03 | 2.52E-02 |
| Neddylation | 220 | 19 | 1.17E-03 | 2.54E-02 |
| RUNX1 regulates transcription of genes involved in differentiation of HSCs | 65 | 9 | 1.26E-03 | 2.70E-02 |
| Hedgehog 'on' state | 79 | 10 | 1.28E-03 | 2.71E-02 |
| Transport of Mature Transcript to Cytoplasm | 81 | 10 | 1.52E-03 | 3.14E-02 |
| Regulation of RAS by GAPs | 67 | 9 | 1.53E-03 | 3.11E-02 |
| Regulation of PTEN stability and activity | 68 | 9 | 1.68E-03 | 3.37E-02 |
| EPH-Ephrin signaling | 69 | 9 | 1.84E-03 | 3.65E-02 |
| Signaling by NTRKs | 70 | 9 | 2.01E-03 | 3.90E-02 |
| Cyclin E associated events during G1/S transition | 70 | 9 | 2.01E-03 | 3.94E-02 |
| Signal transduction by L1 | 21 | 5 | 2.03E-03 | 3.90E-02 |
| Signaling by the B Cell Receptor (BCR) | 100 | 11 | 2.10E-03 | 3.98E-02 |
| Signaling by NTRK1 (TRKA) | 57 | 8 | 2.15E-03 | 4.03E-02 |

| | | | | |
|---|---|---|---|---|
| Synthesis of DNA | 117 | 12 | 2.30E-03 | 4.26E-02 |
| Cyclin A:Cdk2-associated events at S phase entry | 72 | 9 | 2.39E-03 | 4.38E-02 |
| Nuclear Events (kinase and transcription factor activation) | 22 | 5 | 2.42E-03 | 4.39E-02 |
| ERKs are inactivated | 13 | 4 | 2.66E-03 | 4.72E-02 |
| Hedgehog 'off' state | 104 | 11 | 2.78E-03 | 4.87E-02 |
| Class I MHC mediated antigen processing & presentation | 349 | 25 | 2.78E-03 | 4.82E-02 |
| Separation of Sister Chromatids | 182 | 16 | 2.79E-03 | 4.80E-02 |
| Metabolism of carbohydrates | 257 | 20 | 2.85E-03 | 4.85E-02 |
| BBSome-mediated cargo-targeting to cilium | 23 | 5 | 2.86E-03 | 4.82E-02 |

| Upstream Regulator Name | raw p-value | B-H corrected p-value | Target gene name in analysed list |
|---|---|---|---|
| camptothecin | 1.37E-07 | 0.000449 | ARAP2,ASNSD1,CASP4,CCR1,CLU,CSNK1D,DUSP6,FLT1,GADD45G,GSDN |
| TP53 | 3.41E-07 | 0.000559 | ACADM,ACSF2,AK1,AKR1B1,ANAPC4,ARAP2,ARL6IP1,ARVCF,ATG4C,AT |
| 1,2-dithiol-3-thione | 0.00000226 | 0.00247 | ANG,ATP1A1,CCT3,CCT7,CTSD,EIF3G,GNA11,HMBS,HSP90AB1,LGALS8, |
| RXRB | 0.00000498 | 0.00408 | ACADM,ACOX1,CAMK1D,CLU,CNNM1,DDX60,EEF2,FGFR1,GADD45G,M |
| HNF4A | 0.0000138 | 0.00902 | ABCF3,ACOX1,AKR1B1,ANG,ANKRA2,APOB,ARHGAP35,ATG4C,BAG6,B |
| MYC | 0.0000265 | 0.0118 | ABCC4,ABCF3,ACADM,AIMP2,Arf2,ARL6IP1,BUB1B,CCNB2,CCT3,CLU,C |
| CSF1 | 0.0000284 | 0.0118 | ABCA9,CCR1,CSF1R,CTSD,CX3CR1,ETS2,FCGR2A,GAB3,GAS7,GOT2,IRF5 |
| resiquimod | 0.0000289 | 0.0118 | APOBEC3B,ARPC4,CASP4,CCR1,CSF1R,DDX60,DOCK10,ETS2,FCGR2A,H |
| MAPT | 0.0000532 | 0.0194 | Anp32a,Arf2,CCT7,CLTA,Cox6c,CTSD,CTSZ,DDX60,EEF2,FKBP1A,FYN,G |
| ESR2 | 0.00013 | 0.0398 | ABCC4,ABHD2,ARID4A,CCT3,CCT7,CLU,CNNM1,CTSD,DDX10,DOCK10,E |
| 5-fluorouracil | 0.000134 | 0.0398 | ANKRA2,APOBEC3B,CCT3,CLU,CSF1R,CTSD,EEF2,EIF3G,FBXW7,FKBP1A |
| APP | 0.000195 | 0.0504 | AK1,Anp32a,ASCC1,CCNB2,CLTA,CLU,Cox6c,CSF1R,CSF2RB,CTSD,DAXX |
| IND S1 | 0.0002 | 0.0504 | ETFA,GLOD4,HIBCH,PRDX1,TUBBETFA,GLOD4,HIBCH,PRDX1,TUBB |
| RICTOR | 0.000246 | 0.0574 | ATP6V1G2,Cox6c,DAXX,NDUFA7,NDUFAB1,NDUFB4,NDUFC2,NDUFS6, |
| copper gluconate | 0.000658 | 0.142 | BIVM,FANCF,LIG3,RBM14,USP1BIVM,FANCF,LIG3,RBM14,USP1 |
| ATN1 | 0.000692 | 0.142 | ATP1A1,CCNDBP1,CSNK1D,EVL,FBXW7,GAS7,SOX4,SPP1,SPTBN2,THRA |
| PLX5622 | 0.000746 | 0.144 | ABCA9,ATP6V1G2,CCR1,CSF2RB,CTSD,CX3CR1,FCGR2A,HEXB,HSP90AB |
| ALKBH5 | 0.000875 | 0.145 | CLU,CTSD,EIF4G3,FBXL5,FGFR1,FYN,MYO6,NONO,SLC4A2 |
| CST5 | 0.000876 | 0.145 | DNAJA3,EMP3,HS2ST1,MAP1B,MEF2C,NDUFC2,NUP107,PRDX1,PRKRA |
| HTT | 0.000907 | 0.145 | ACADM,Anp32a,ATP1A1,ATP8A1,CSF1R,CTSD,DUSP6,EEF2,EMP3,EVL,F |
| HIC1 | 0.000929 | 0.145 | ABHD3,CDO1,LGALS8,MATCAP2,SFXN3,SMPDL3A,SPP1 |
| ciprofloxacin | 0.000999 | 0.149 | ABCC4,CCR1,CPD,CSF2RB,CTSD,HSPA9,MAPK1,MSR1,PLXNB2,SPP1 |
| KLF3 | 0.00119 | 0.167 | AGGF1,ARPC4,CCT3,CEP170B,CTNNA1,DFFA,GALNT1,GPSM1,HEXB,MC |
| HERC2 | 0.00126 | 0.167 | FBXL5,PYCARD,USP16FBXL5,PYCARD,USP16FBXL5,PYCARD,USP16 |
| gentamicin | 0.00131 | 0.167 | ACADM,AGMO,AK1,ASNSD1,CAND1,CDO1,CIPC,CLU,CTSD,EXTL3,JMY,N |
| FAS | 0.00133 | 0.167 | ARAP2,CCR1,FAIM2,FLT1,GADD45G,INPP4A,MEF2C,NLRP1,P2RX1,PAK |
| Immunoglobulin | 0.00148 | 0.179 | BSCL2,C17orf99,CASP4,CLU,CSF1R,CSF2RB,CTSD,CTSZ,CX3CR1,DUSP6, |
| poly rI:rC-RNA | 0.00173 | 0.196 | Anp32a,CCNB2,CCR1,CCT7,CHD7,COQ9,DAXX,DDX60,GBP5,HELZ2,HLA |
| TRAPPC1 | 0.00205 | 0.196 | EDEM2,HSP90AB1,SEL1L,SSR3,SVIP,SYVN1,UBE2J2,UBQLN2 |
| prostaglandin J2 | 0.00213 | 0.196 | PSMB2,PSMB4,PSMD12,PSMD6,PSMD8 |
| NCSTN | 0.00213 | 0.196 | CSF1R,CSF2RB,FCGR2A,PSEN2,PSENEN |
| BDNF | 0.00216 | 0.196 | ACOT7,ARVCF,CAV2,DUSP6,EEF2,KCNIP2,MAG,MALAT1,MAP1B,MAPK |
| IL3 | 0.00223 | 0.196 | ACOX1,CSF1R,EIF3G,ELAVL3,EMP3,FCGR2A,FGFR1,GADD45G,HSPA9,LY |
| topotecan | 0.0023 | 0.196 | ANKMY2,ARL6IP1,CACNA1C,CAMK1D,CHD1,EXOC6B,FHIT,MAPK1,MYB |
| garcinol | 0.00236 | 0.196 | ARHGAP35,BAZ1A,BUB1B,CASP4,FAIM2,GTF2I,MEF2C,TPT1 |
| Z-LLL-CHO | 0.00237 | 0.196 | AKR1B1,ATP1A1,BUB1B,CHD1,CLU,DROSHA,HSPA9,MYB,NFKB2,PAK2, |
| GABA | 0.00238 | 0.196 | ACADM,ATP1A1,B4GALT6,Ccl27a,CHPT1,DOLK,EIF2AK4,HIBCH,IDS,MA |
| IL4 | 0.00238 | 0.196 | ACOX1,C17orf99,CAMK1D,CAND1,Ccl27a,CCR1,CCT3,CSF1R,CSF2RB,CT |
| disulfiram | 0.00242 | 0.196 | BIVM,FANCF,LIG3,RBM14,USP1BIVM,FANCF,LIG3,RBM14,USP1 |
| MKNK1 | 0.00245 | 0.196 | 4933406I18Rik,ACOT7,ARVCF,CLU,Gm16897,MALAT1,MAP1B,MAPK1, |
| EIF2AK3 | 0.00249 | 0.196 | ANG,Arf2,ETS2,HSPA9,NFKB2,NUPR1,SEL1L,Slc35a2,SYS1,SYVN1,USO1 |
| lipopolysaccharide | 0.00251 | 0.196 | 9930111J21Rik1/Gm12185,ABCC4,ACADM,ADRA2A,ANKMY2,ANXA7,A |
| HIVEP1 | 0.00278 | 0.211 | CARD11,NFKB2,PIK3R6,TLR5,TNFSF13B,TNIP1 |
| CLOCK | 0.003 | 0.211 | ACOX1,CPD,LGALS8,LIG3,NR2C1,PICALM,SEL1L,SERINC5,SLC39A12,ST3 |
| HSF2 | 0.00306 | 0.211 | CCT3,CCT7,CLU,PSMB2,PSMC4CCT3,CCT7,CLU,PSMB2,PSMC4 |
| Z-DEVD-FMK | 0.00311 | 0.211 | MEF2C,PDZD2,PLD1MEF2C,PDZD2,PLD1MEF2C,PDZD2,PLD1 |
| ETS2 | 0.00311 | 0.211 | CSF1R,DUSP6,FLT1,MGAT5,MSR1,PLAU,SPP1 |
| CCND1 | 0.00312 | 0.211 | ABCC4,DONSON,E2F4,FGFR1,HJURP,KDM6B,KLHL24,NRF1,PTBP3,RAB3 |
| PSEN1 | 0.0032 | 0.211 | 2700097O09Rik,Anp32a,CLTA,Cox6c,CTSD,CTSZ,FBXW7,FKBP1A,GOT2, |
| TGFB1 | 0.00323 | 0.211 | ADRA2A,AFM,AK1,AP3S1,APOB,ARHGAP35,ATG14,BUB1B,CAPRIN1,CA |
| LMO2 | 0.00334 | 0.211 | CASP4,FZD5,GALNT1,GBP5,JARID2,MEF2C,NLRP1,OXCT1,P2RX1,PDZD2 |
| mibolerone | 0.00335 | 0.211 | CHPT1,CLTA,CLU,FKBP1A,GADD45G,GOT2,HEXB,SMPDL3A,UFM1,USO1 |
| cisplatin | 0.00359 | 0.222 | ABCC4,ACADM,ACOX1,APOBEC3B,ASCC1,ASRGL1,ATG14,CCR1,CDO1,C |
| NUP133 | 0.00381 | 0.223 | NUP107,NUP153NUP107,NUP153NUP107,NUP153NUP107,NUP153 |
| UFD1 | 0.00381 | 0.223 | NFKB2,TNFSF13BNFKB2,TNFSF13BNFKB2,TNFSF13BNFKB2,TNFSF13B |

| | | | |
|---|---|---|---|
| NPLOC4 | 0.00381 | 0.223 | NFKB2,TNFSF13BNFKB2,TNFSF13BNFKB2,TNFSF13BNFKB2,TNFSF13B |
| miR-291a-3p (and other miRNA | 0.00397 | 0.228 | BAZ1A,C9orf78,FYCO1,LUC7L2,NUP58,TFAP4,TMEM9B,TNFAIP1 |
| LDB1 | 0.00435 | 0.245 | CASP4,FZD5,GALNT1,GBP5,JARID2,MEF2C,NLRP1,OXCT1,P2RX1,PDZD2 |
| vancomycin | 0.00449 | 0.249 | AGMO,AK1,CLU,EMP3,EXTL3,FCGR2A,HJURP,JMY,PTPRM,SOX4,SPP1,S |
| DTX1 | 0.00471 | 0.252 | DUSP6,NAE1,SPP1,UHMK1DUSP6,NAE1,SPP1,UHMK1 |
| interferon beta-1a | 0.00473 | 0.252 | BUB3,CSF1R,FKBP1A,GBP5,HLA-DRB5,MRPL20,PIN1,RPL3,TNFSF13B |
| PTPRR | 0.00476 | 0.252 | FYCO1,ITGA5,LENG1,PBXIP1,PFKL,PLXNB2,TNFAIP1,TNIP1 |
| miR-150-5p (and other miRNAs | 0.00495 | 0.257 | CSF1R,MALAT1,MYBCSF1R,MALAT1,MYBCSF1R,MALAT1,MYB |
| IFNG | 0.00502 | 0.257 | ADRA2A,ANKS1A,ARAP2,ATP1A1,BUB1B,CASP4,Ccl27a,CCR1,CSF1R,CS |
| KLF6 | 0.00538 | 0.268 | ACADM,CASP4,CCR1,COG2,CSF2RB,ETS2,HIBCH,ITGA5,KDM6B,NFKB2, |
| filgrastim | 0.0054 | 0.268 | ATP1A1,CATSPER2,CLU,CPD,CSF1R,DDHD1,ETS2,GBP5,HLA-DMB,HLA- |
| SETD2 | 0.00572 | 0.272 | ANG,CSNK1D,FLT1,MEF2C,SERPINF1 |
| PTEN | 0.00581 | 0.272 | ABCA9,ABCC4,ACOX1,CCNB2,CHD1,CLTA,CTNNA1,CTSZ,CX3CR1,ETS2,F |
| TCF4 | 0.00595 | 0.272 | CASP4,CCNB2,EXOSC7,FARSB,Hmgn2 (includes others),MEF2C,NANOS |
| ARID1A | 0.00603 | 0.272 | ARVCF,CCNB2,FZD5,HIBCH,ITGA5,L1CAM,LRP5,MEF2C,MMS19,MYO6, |
| SERTAD2 | 0.00607 | 0.272 | ACOX1,NRF1,PHYHACOX1,NRF1,PHYHACOX1,NRF1,PHYH |
| AF-2364 | 0.00607 | 0.272 | MAPK1,PAK2,PIK3CAMAPK1,PAK2,PIK3CAMAPK1,PAK2,PIK3CA |
| dexamethasone | 0.00612 | 0.272 | ABHD2,ACADM,ACOT7,ACOX1,ADRA2A,AP3S1,ATP1A1,BSCL2,CATSPEF |
| BLOC1S6 | 0.00624 | 0.272 | PI4K2A,STX12PI4K2A,STX12PI4K2A,STX12PI4K2A,STX12PI4K2A,STX12 |
| TMPRSS4 | 0.00624 | 0.272 | ITGA5,PLAUITGA5,PLAUITGA5,PLAUITGA5,PLAUITGA5,PLAU |
| miR-182-5p (and other miRNAs | 0.00638 | 0.273 | CARD11,DDX60,GADD45G,MALAT1,NLGN2,PIK3CA |
| GnRH analog | 0.00642 | 0.273 | ACOT7,BUB3,CLTA,DAXX,FKBP1A,FLT1,HADHB,HSP90AB1,MAPK1,N4B |
| ESR1 | 0.00652 | 0.274 | ABHD2,ACOX1,AGGF1,AK1,ARID4A,ARSG,BUB1B,BUB3,CAV2,CEP250,C |
| TLE3 | 0.00704 | 0.289 | ECSIT,ETFA,NDUFA7,NDUFAB1,NDUFB4,NDUFC2,NDUFS6 |
| colistin | 0.00707 | 0.289 | AGMO,AK1,EXTL3,JMY,PTPRM,SOX4,SPP1,SYNJ1 |
| MEL T1 | 0.00733 | 0.295 | GLOD4,HIBCH,PRDX1GLOD4,HIBCH,PRDX1GLOD4,HIBCH,PRDX1 |
| SOX7 | 0.00739 | 0.295 | FGFR1,FLT1,LRP5,PLAU,SOX4,VCAN |
| kanamycin A | 0.00786 | 0.309 | AGMO,AK1,EXTL3,JMY,PTPRM,SOX4,SPP1,SYNJ1 |
| tretinoin | 0.00792 | 0.309 | ABHD2,ACOX1,B3GALT5,CACNA1C,CCR1,CSF1R,CTSD,CX3CR1,DDX10,D |
| geldanamycin | 0.00808 | 0.311 | AKR1B1,ANXA7,CLU,FGFR1,FYN,HEXB,HLTF,HSP90AB1,IRF5,NAE1,NUD |
| 2-bromoethylamine | 0.00822 | 0.313 | CLU,EMP3,FCGR2A,SPP1,TUBBCLU,EMP3,FCGR2A,SPP1,TUBB |
| CpG ODN 2006 | 0.00835 | 0.314 | ARPC4,DOCK10,GBP5,HSP90AB1,NFAM1,NFKB2,PSMB2,PSMC4,PSMD |
| IL10 | 0.00866 | 0.314 | ANKS1A,ARAP2,BUB1B,CCR1,CSF1R,CSF2RB,CTSD,CTSZ,CX3CR1,ETS2,F |
| STK3 | 0.00873 | 0.314 | SOX4,SPP1,WWTR1SOX4,SPP1,WWTR1SOX4,SPP1,WWTR1 |
| USP1 | 0.00873 | 0.314 | FYN,ITGA5,USP1FYN,ITGA5,USP1FYN,ITGA5,USP1FYN,ITGA5,USP1 |
| Go6983 | 0.00873 | 0.314 | ACADM,FLT1,PRDX1ACADM,FLT1,PRDX1ACADM,FLT1,PRDX1 |
| metribolone | 0.00885 | 0.315 | ABCC4,ABHD2,ARL6IP1,CAV2,CLU,ETFA,FZD5,GADD45G,GOT2,HIBCH,N |
| nilotinib | 0.00894 | 0.315 | ABCC4,ANAPC4,BUB3,DBF4,PIK3CA |
| 6-hydroxydopamine | 0.00917 | 0.317 | ASCC1,GADD45G,HSP90AB1,HSPA9,MAPK1,PSMB2,PSMC4,PYCARD |
| antrodia camphorata polysacch | 0.0092 | 0.317 | ASCC1,PYCARDASCC1,PYCARDASCC1,PYCARDASCC1,PYCARD |
| fluoromethyl 2,2-difluoro-1-(tri | 0.00972 | 0.328 | ANXA7,CDO1,CLU,SPP1ANXA7,CDO1,CLU,SPP1ANXA7,CDO1,CLU,SPP1 |
| SLC22A5 | 0.00972 | 0.328 | ACADM,HADHB,PLAU,SMAD1ACADM,HADHB,PLAU,SMAD1 |
| NFE2L2 | 0.00992 | 0.331 | ABCC4,ANG,ATP1A1,CCT3,CCT7,CTSD,EIF3G,GADD45G,GNA11,HMBS,H |
| mir-322 | 0.0103 | 0.34 | FGFR1,LYPLA2,MAP2K1FGFR1,LYPLA2,MAP2K1FGFR1,LYPLA2,MAP2K1 |
| HYAL1 | 0.0105 | 0.34 | C1orf54,MAP1B,NLRP1,SPP1,TUBBC1orf54,MAP1B,NLRP1,SPP1,TUBB |
| UBQLN2 | 0.0105 | 0.34 | PSMB2,PSMB4,PSMD12,PSMD2,PSMD8 |
| BMP10 | 0.0106 | 0.34 | CCR1,CX3CR1,FCGR2A,MEF2C,MSR1,SKIL,VCAN |
| dithiothreitol | 0.0108 | 0.343 | SEL1L,SYVN1,TLR2,UBE3ASEL1L,SYVN1,TLR2,UBE3A |
| Akt | 0.0111 | 0.346 | AIMP2,CPD,CTSD,E2F4,FHIT,GCNT2,L1CAM,LMBRD1,MGAT5,MYB,NRF |
| diphtheria toxin | 0.0112 | 0.346 | AGMO,CTSD,DONSON,DUSP6,MRPL20,OPHN1,PLAU,RPL3,TLR2 |
| desmopressin | 0.0112 | 0.346 | AKR1B1,Anp32a,ATP1A1,CTSD,RBM14,SPTBN2,TUBB |
| ouabain | 0.0114 | 0.348 | ATP1A1,EIF2AK4,FLT1,HSPA9,MAPK1 |
| L-methionine | 0.0118 | 0.358 | CCNB2,CLU,ETS2,GNA11,PIN1,SPP1 |
| ERBB2 | 0.0123 | 0.367 | AIMP2,ANG,BUB1B,CCNB2,CHD7,CLU,CPD,CSF1R,CX3CR1,DDX10,DUSF |
| E2F2 | 0.0126 | 0.367 | CCNB2,ECE1,JMY,MYB,PIN1,TOP2BCCNB2,ECE1,JMY,MYB,PIN1,TOP2B |
| CDC25A | 0.0127 | 0.367 | MPHOSPH6,NAT10MPHOSPH6,NAT10MPHOSPH6,NAT10 |
| PSENEN | 0.0127 | 0.367 | PSEN2,PSENENPSEN2,PSENENPSEN2,PSENENPSEN2,PSENEN |
| juglone | 0.0127 | 0.367 | E2F4,PIN1E2F4,PIN1E2F4,PIN1E2F4,PIN1E2F4,PIN1E2F4,PIN1 |
| RXRA | 0.0137 | 0.378 | ACADM,ACOX1,CAMK1D,CDO1,CLU,CNNM1,DDX60,EEF2,GADD45G,H |

| | | | |
|---|---|---|---|
| SYVN1 | 0.0138 | 0.378 | ABCC4,ATP1A1,GGCX,L1CAM,NFKB2,RPL10,SETX,SLC4A7,STOM |
| ASPSCR1-TFE3 | 0.0138 | 0.378 | ABHD2,CTSD,RAB32,SFXN3,STOM,THEM6,UAP1L1 |
| metronidazole | 0.0141 | 0.378 | AGMO,AK1,EXTL3,JMY,PTPRM,SOX4,SPP1,SYNJ1 |
| NGLY1 | 0.0145 | 0.378 | ATG14,GABPB1,NRF1,WIPI2ATG14,GABPB1,NRF1,WIPI2 |
| Irgm1 | 0.0145 | 0.378 | APOBEC3B,CCNB2,DDX60,GBP5,GSDMD,NUPR1,TLR2 |
| SETBP1 | 0.015 | 0.378 | BUB1B,CARD11,FBXW7,GNA11,MYB,WWTR1 |
| UCHL1 | 0.015 | 0.378 | CPD,DOCK10,GABPB1,JMY,PHF20,PIK3CG |
| carboplatin | 0.0153 | 0.378 | CLU,FCGR2A,PNKD,SPP1,TUBBCLU,FCGR2A,PNKD,SPP1,TUBB |
| GRN | 0.0154 | 0.378 | CLU,CTSD,CTSZ,CX3CR1,HEXB,MEF2C,PICALM,SPP1 |
| SMARCA4 | 0.0155 | 0.378 | ABHD2,ASCC1,C1orf54,CCR1,CLK1,DUSP6,EMP3,EPHA1,EPHA4,ETS2,F[illegible] |
| PAX3-FOXO1 | 0.0155 | 0.378 | ADRA2A,AKR1B1,ATP1A1,CEP170B,CHD7,EEF2,FOXO4,ITGA5,JARID2,N[illegible] |
| miR-293-5p (and other miRNAs | 0.0158 | 0.378 | GOLPH3,MAPK1,SSR3GOLPH3,MAPK1,SSR3GOLPH3,MAPK1,SSR3 |
| NUP98-NSD1 | 0.0158 | 0.378 | MSR1,MYB,SOX4MSR1,MYB,SOX4MSR1,MYB,SOX4MSR1,MYB,SOX4 |
| GPR174 | 0.0161 | 0.378 | ARAP2,CARD11,CBLB,ETS2,FYN,KDM6B,SPATA13 |
| Tcf7 | 0.0164 | 0.378 | CX3CR1,FGFR1,FYN,GCNT2,KCNJ3,MAPKAPK2,MGAT5,NHSL2,NUPR1,P[illegible] |
| methyl methanesulfonate | 0.0164 | 0.378 | CTSD,GADD45G,NUPR1,PLXNB2,SYVN1 |
| ETV3 | 0.0166 | 0.378 | MYB,SPP1MYB,SPP1MYB,SPP1MYB,SPP1MYB,SPP1MYB,SPP1 |
| NUP155 | 0.0166 | 0.378 | CACNA1C,PlnCACNA1C,PlnCACNA1C,PlnCACNA1C,PlnCACNA1C,Pln |
| EGR2 | 0.0173 | 0.378 | CBLB,CPD,CSF1R,EPHA4,MAG,MAP7,MYB,NFKB2,NRP2,PDE7A,PPT2,S[illegible] |
| HDL-cholesterol | 0.0173 | 0.378 | ALDH1L2,APOB,KLHL24,NUPR1ALDH1L2,APOB,KLHL24,NUPR1 |
| butyric acid | 0.0177 | 0.378 | ARID4A,CCNB2,CCR1,CLU,E2F4,EMP3,FYN,HADHB,ITGA5,MYB,MYH14,[illegible] |
| RBL2 | 0.0177 | 0.378 | BUB1B,CASP4,CCNB2,E2F4,FGFR1,PSMB2,RANBP1 |
| TNF | 0.0181 | 0.378 | ACADM,ACOX1,AKR1B1,APOBEC3B,ATP1A1,BUB1B,CASP4,CCDC15,Ccl[illegible] |
| ERG | 0.0182 | 0.378 | ABCC4,CAMK1D,CTNNA1,DOCK10,FLT1,FYN,NPHP1,PLAU,PRKCI,RAPG[illegible] |
| ETS1 | 0.0182 | 0.378 | CSF1R,DNAJA3,ETS2,FCGR2A,FLT1,HS2ST1,MGAT5,MSR1,MYB,PLAU,S[illegible] |
| mir-182 | 0.0188 | 0.378 | CARD11,CSF1R,GADD45G,MALAT1,NLGN2,PIK3CA |
| LONP1 | 0.0188 | 0.378 | ALDH1L2,EARS2,ECSIT,HSPA9,MTHFD2,OXCT1 |
| MMP3 | 0.0195 | 0.378 | CAPRIN1,PPIL1,RBM14,SF3A3,SNRNP48,THOC6,YBX1 |
| TNFSF11 | 0.0198 | 0.378 | CCR1,CSF1R,DUSP6,FAM102A,ITGA5,MAP2K1,NFKB2,NME6,PLD1,SEL1[illegible] |
| IFN alpha/beta | 0.0199 | 0.378 | DAXX,GAS7,TLR2,TLR5,TNFSF13,TNFSF13B |
| levodopa | 0.02 | 0.378 | AGGF1,ASTN1,B3GALT5,CTSZ,DUSP6,ECE1,ETS2,FAM3A,GAS7,HEXB,L[illegible] |
| CLEC4A | 0.02 | 0.378 | ABCA9,LYVE1,MSR1,TLR2,VCANABCA9,LYVE1,MSR1,TLR2,VCAN |
| PGF | 0.0205 | 0.378 | ETS2,FLT1,NANOS1,NRP2ETS2,FLT1,NANOS1,NRP2 |
| LARP1 | 0.021 | 0.378 | EEF2,RPL10,RPL3,RPL37A,RPS23,TPT1 |
| APH1A | 0.021 | 0.378 | PSEN2,PSENENPSEN2,PSENENPSEN2,PSENENPSEN2,PSENEN |
| SNHG7 | 0.021 | 0.378 | FAIM2,GALNT1FAIM2,GALNT1FAIM2,GALNT1FAIM2,GALNT1 |
| CNBP | 0.021 | 0.378 | AK1,DUSP3AK1,DUSP3AK1,DUSP3AK1,DUSP3AK1,DUSP3AK1,DUSP3 |
| PTGES3 | 0.021 | 0.378 | CTSD,NUPR1CTSD,NUPR1CTSD,NUPR1CTSD,NUPR1CTSD,NUPR1 |
| MNX1 | 0.021 | 0.378 | CBLB,NONOCBLB,NONOCBLB,NONOCBLB,NONOCBLB,NONO |
| PPP1CA | 0.021 | 0.378 | SPP1,WWTR1SPP1,WWTR1SPP1,WWTR1SPP1,WWTR1SPP1,WWTR1 |
| D-thioctic acid | 0.021 | 0.378 | GABPB1,NRF1GABPB1,NRF1GABPB1,NRF1GABPB1,NRF1 |
| sivelestat | 0.021 | 0.378 | TNFSF13,TNFSF13BTNFSF13,TNFSF13BTNFSF13,TNFSF13B |
| tunicamycin | 0.0219 | 0.378 | ACOX1,Arf2,CASP4,EDEM2,HSPA9,SEC11A,SEL1L,Slc35a2,SSR3,SYS1,S[illegible] |
| CIITA | 0.0222 | 0.378 | GCNT2,HLA-DMB,HLA-DOB,HLA-DRB5 |
| IL2 | 0.0223 | 0.378 | CASP4,CBLB,CCR1,CPD,CSF1R,CSF2RB,CX3CR1,DUSP6,DUSP7,E2F4,EP[illegible] |
| TEAD1 | 0.0226 | 0.378 | ACADM,Cox6c,ETFA,HADHB,NDUFA7,NDUFAB1,NDUFB4,NDUFC2,ND[illegible] |
| RET | 0.0234 | 0.378 | CLU,DUSP6,EIF4G3,HSPA9,L3MBTL2,MAPKAPK2,PLAU,Pln,UBQLN2 |
| CCL20 | 0.024 | 0.378 | CX3CR1,GBP5,PIK3CG,ZC3H12ACX3CR1,GBP5,PIK3CG,ZC3H12A |
| RBM20 | 0.0243 | 0.378 | CSDE1,EEF2,FASTK,IMPACT,RPL3,SOX4,TNRC6B |
| PRDM5 | 0.0244 | 0.378 | APOBEC3B,CACNA1C,ETS2,EVL,MYB,SNRNP48 |
| ZC3H12A | 0.0245 | 0.378 | ABCC4,Anp32a,ARAP2,GBP5,IRF5,NFKB2,PDE7A,PIK3CG,RAPGEF6,TNI[illegible] |
| ST1926 | 0.0251 | 0.378 | AKR1B1,CCT3,EEF2,EIF3G,PIN1,RPL10,SOX4,TPT1,YBX1 |
| FLCN | 0.0252 | 0.378 | CTSZ,HEXB,L1CAM,NDUFB4,SDCBP,SLC17A5,UAP1L1,ZC3H12A |
| NPC1 | 0.0252 | 0.378 | APOB,CAV2,CCR1,CSF1R,CTSD,CTSZ,CX3CR1,HLA-DRB5,PI4K2A,PRDX1 |
| PRL | 0.0253 | 0.378 | CLU,CPD,CTSD,GPSM1,HELZ2,HNRNPH2,LYVE1,N4BP1,NUPR1,PAQR8,[illegible] |
| clofibrate | 0.0254 | 0.378 | ABCC4,ACOX1,ANXA7,HADHB,Hmgn2 (includes others),MYB,NR2C1 |
| TP73 | 0.0255 | 0.378 | AGGF1,ARVCF,ATG4C,CASP4,CASS4,CTSD,DCAF6,FLT1,GPR137B,INPP4[illegible] |
| CD247 | 0.0256 | 0.378 | ANG,ETFA,FCGR2A,MTHFD2,SRP14ANG,ETFA,FCGR2A,MTHFD2,SRP14 |
| lomustine | 0.0256 | 0.378 | CLU,EMP3,FCGR2A,SPP1,TUBBCLU,EMP3,FCGR2A,SPP1,TUBB |

| | | | |
|---|---|---|---|
| STF-1019 | 0.0256 | 0.378 | PI4K2API4K2API4K2API4K2API4K2API4K2API4K2API4K2API4K2A |
| miR-373 inhibitor | 0.0256 | 0.378 | PIK3CAPIK3CAPIK3CAPIK3CAPIK3CAPIK3CAPIK3CAPIK3CAPIK3CA |
| PIK3R6 | 0.0256 | 0.378 | PIK3CGPIK3CGPIK3CGPIK3CGPIK3CGPIK3CGPIK3CGPIK3CGPIK3CG |
| FB23-2 | 0.0256 | 0.378 | APOBEC3BAPOBEC3BAPOBEC3BAPOBEC3BAPOBEC3BAPOBEC3BAPOBEC3B |
| capsular polysaccharide | 0.0256 | 0.378 | MSR1MSR1MSR1MSR1MSR1MSR1MSR1MSR1MSR1MSR1MSR1MSR1 |
| ORG 34517 | 0.0256 | 0.378 | NR3C2NR3C2NR3C2NR3C2NR3C2NR3C2NR3C2NR3C2NR3C2NR3C2 |
| coactivator-Dtx-Notch-Rbpsuh | 0.0256 | 0.378 | MAGMAGMAGMAGMAGMAGMAGMAGMAGMAGMAGMAGMAG |
| 1H,1H,2H,2H-perfluorodecanol | 0.0256 | 0.378 | ACOX1ACOX1ACOX1ACOX1ACOX1ACOX1ACOX1ACOX1ACOX1ACOX1 |
| SNHG17 | 0.0256 | 0.378 | SOX4SOX4SOX4SOX4SOX4SOX4SOX4SOX4SOX4SOX4SOX4SOX4SOX4 |
| CWH43 | 0.0256 | 0.378 | L1CAML1CAML1CAML1CAML1CAML1CAML1CAML1CAML1CAM |
| OVGP1 | 0.0256 | 0.378 | MAPKAPK2MAPKAPK2MAPKAPK2MAPKAPK2MAPKAPK2MAPKAPK2 |
| REEP5 | 0.0256 | 0.378 | TOP2BTOP2BTOP2BTOP2BTOP2BTOP2BTOP2BTOP2BTOP2BTOP2B |
| KIF21A | 0.0256 | 0.378 | MAP1BMAP1BMAP1BMAP1BMAP1BMAP1BMAP1BMAP1BMAP1B |
| DNAJC24 | 0.0256 | 0.378 | EEF2EEF2EEF2EEF2EEF2EEF2EEF2EEF2EEF2EEF2EEF2EEF2EEF2EEF2 |
| HDLBP | 0.0256 | 0.378 | CSF1RCSF1RCSF1RCSF1RCSF1RCSF1RCSF1RCSF1RCSF1RCSF1RCSF1R |
| B3GALT1 | 0.0256 | 0.378 | B3GALT5B3GALT5B3GALT5B3GALT5B3GALT5B3GALT5B3GALT5 |
| SAMD4B | 0.0256 | 0.378 | NANOS1NANOS1NANOS1NANOS1NANOS1NANOS1NANOS1NANOS1 |
| HOOK3 | 0.0256 | 0.378 | MSR1MSR1MSR1MSR1MSR1MSR1MSR1MSR1MSR1MSR1MSR1MSR1 |
| ASTN2 | 0.0256 | 0.378 | ASTN1ASTN1ASTN1ASTN1ASTN1ASTN1ASTN1ASTN1ASTN1ASTN1 |
| CPB1 | 0.0256 | 0.378 | CASP4CASP4CASP4CASP4CASP4CASP4CASP4CASP4CASP4CASP4 |
| DPH2 | 0.0256 | 0.378 | EEF2EEF2EEF2EEF2EEF2EEF2EEF2EEF2EEF2EEF2EEF2EEF2EEF2EEF2 |
| IL34 | 0.0256 | 0.378 | CSF1RCSF1RCSF1RCSF1RCSF1RCSF1RCSF1RCSF1RCSF1RCSF1RCSF1R |
| roxadustat | 0.0256 | 0.378 | SERPINF1SERPINF1SERPINF1SERPINF1SERPINF1SERPINF1SERPINF1 |
| afuresertib | 0.0256 | 0.378 | CBLBCBLBCBLBCBLBCBLBCBLBCBLBCBLBCBLBCBLBCBLBCBLBCBLB |
| CACNB2 | 0.0256 | 0.378 | CACNA1CCACNA1CCACNA1CCACNA1CCACNA1CCACNA1CCACNA1C |
| mir-3202 | 0.0256 | 0.378 | FAIM2FAIM2FAIM2FAIM2FAIM2FAIM2FAIM2FAIM2FAIM2FAIM2 |
| miR-125a-3p (miRNAs w/seed C | 0.0256 | 0.378 | FYNFYNFYNFYNFYNFYNFYNFYNFYNFYNFYNFYNFYNFYNFYNFYN |
| miR-105-5p (and other miRNAs | 0.0256 | 0.378 | TLR2TLR2TLR2TLR2TLR2TLR2TLR2TLR2TLR2TLR2TLR2TLR2TLR2TLR2 |
| miR-1225-3p (miRNAs w/seed C | 0.0256 | 0.378 | GAB3GAB3GAB3GAB3GAB3GAB3GAB3GAB3GAB3GAB3GAB3GAB3 |
| mir-1225 | 0.0256 | 0.378 | GAB3GAB3GAB3GAB3GAB3GAB3GAB3GAB3GAB3GAB3GAB3GAB3 |
| PIK3R5 | 0.0256 | 0.378 | PIK3CGPIK3CGPIK3CGPIK3CGPIK3CGPIK3CGPIK3CGPIK3CGPIK3CG |
| KCNJ9 | 0.0256 | 0.378 | KCNJ3KCNJ3KCNJ3KCNJ3KCNJ3KCNJ3KCNJ3KCNJ3KCNJ3KCNJ3KCNJ3 |
| BARX1 | 0.0256 | 0.378 | L1CAML1CAML1CAML1CAML1CAML1CAML1CAML1CAML1CAM |
| phenanthroline-(2H,7H)-diaceta | 0.0256 | 0.378 | PIN1PIN1PIN1PIN1PIN1PIN1PIN1PIN1PIN1PIN1PIN1PIN1PIN1PIN1 |
| cyclo(D-Arg-D-Arg-DThr(P)-Pip-N | 0.0256 | 0.378 | PIN1PIN1PIN1PIN1PIN1PIN1PIN1PIN1PIN1PIN1PIN1PIN1PIN1PIN1 |
| clomoxir | 0.0256 | 0.378 | ACADMACADMACADMACADMACADMACADMACADMACADMACADM |
| BC-DXI01 | 0.0256 | 0.378 | AIMP2AIMP2AIMP2AIMP2AIMP2AIMP2AIMP2AIMP2AIMP2AIMP2 |
| dexamethasone/tobramycin | 0.0256 | 0.378 | TLR2TLR2TLR2TLR2TLR2TLR2TLR2TLR2TLR2TLR2TLR2TLR2TLR2TLR2 |
| dihydroxyphenylethylene glycol | 0.0256 | 0.378 | MAP1BMAP1BMAP1BMAP1BMAP1BMAP1BMAP1BMAP1BMAP1B |
| CIC-DUX4 | 0.0256 | 0.378 | DUSP6DUSP6DUSP6DUSP6DUSP6DUSP6DUSP6DUSP6DUSP6DUSP6 |
| miR-216b inhibitor | 0.0256 | 0.378 | GALNT1GALNT1GALNT1GALNT1GALNT1GALNT1GALNT1GALNT1 |
| CA-074 | 0.0256 | 0.378 | CTSZCTSZCTSZCTSZCTSZCTSZCTSZCTSZCTSZCTSZCTSZCTSZCTSZCTSZ |
| oxfenicine | 0.0256 | 0.378 | ACADMACADMACADMACADMACADMACADMACADMACADMACADM |
| D-mannoheptulose | 0.0256 | 0.378 | APOBAPOBAPOBAPOBAPOBAPOBAPOBAPOBAPOBAPOBAPOBAPOB |
| CAY10585 | 0.0258 | 0.378 | FLT1,NR3C2FLT1,NR3C2FLT1,NR3C2FLT1,NR3C2FLT1,NR3C2 |
| MCL1 | 0.0258 | 0.378 | TPT1,UAP1L1TPT1,UAP1L1TPT1,UAP1L1TPT1,UAP1L1TPT1,UAP1L1 |
| MLH1 | 0.0258 | 0.378 | CLU,PMS2CLU,PMS2CLU,PMS2CLU,PMS2CLU,PMS2CLU,PMS2 |
| STX3 | 0.0258 | 0.378 | SART3,VCANSART3,VCANSART3,VCANSART3,VCANSART3,VCAN |
| butylhydroxybutylnitrosamine | 0.0258 | 0.378 | FHIT,UGT1A6FHIT,UGT1A6FHIT,UGT1A6FHIT,UGT1A6FHIT,UGT1A6 |
| poloxamer | 0.0258 | 0.378 | ACADM,ACOX1ACADM,ACOX1ACADM,ACOX1ACADM,ACOX1 |
| CSF1R | 0.0259 | 0.378 | CSF1R,GAB3,MSR1,MYBCSF1R,GAB3,MSR1,MYB |
| miR-155-5p (miRNAs w/seed UA | 0.0278 | 0.404 | BRPF3,CSF1R,JARID2,MOSPD2,MYB,PICALM,PRKCI,SDCBP,SMAD1 |
| torin1 | 0.0286 | 0.409 | EEF2,NDUFA7,NDUFAB1,NDUFC2,PFKL,RPL10,RPL3,RPL37A,RPS23,TPT |
| LEF1 | 0.0287 | 0.409 | CARD11,CX3CR1,FGFR1,GCNT2,KCNJ3,MGAT5,NHSL2,PIK3R6,PLXNB2,F |
| CYLD-AS1 | 0.0287 | 0.409 | AKR1B1,IRF2BPL,MYB,NFKB2,PRDX1 |
| mitomycin C | 0.0287 | 0.409 | APOBEC3B,CCNB2,IRF2BPL,MALAT1,PYCARD |
| PRKAG3 | 0.0287 | 0.409 | ACTR1B,CCNDBP1,CTNNA1,FGFR1,GADD45G,LUC7L2,OXCT1,PDE7A,RE |
| triamterene | 0.0303 | 0.428 | CLU,EMP3,FCGR2A,SPP1,TUBBCLU,EMP3,FCGR2A,SPP1,TUBB |
| tamoxifen | 0.0303 | 0.428 | ACOX1,ANG,CAV2,CLU,CTSD,E2F4,EPHA4,GPATCH3,HSP90AB1,KLHL24 |

| | | | |
|---|---|---|---|
| OGA | 0.0305 | 0.428 | ACADM,ARPC4,CAMK1D,CCR1,CDKAL1,CSF1R,CSF2RB,CX3CR1,DAGLB, |
| uridine | 0.031 | 0.428 | ACOX1,HADHBACOX1,HADHBACOX1,HADHBACOX1,HADHB |
| (2-(2-(4-methylthiazol-5-yl)etho | 0.031 | 0.428 | NRF1,RBM14NRF1,RBM14NRF1,RBM14NRF1,RBM14NRF1,RBM14 |
| MYD88 | 0.031 | 0.428 | APOBEC3B,CASP4,DAXX,EDEM2,ELF2,ETS2,FANCF,HSP90AB1,KDM6B,M |
| mir-144 | 0.0311 | 0.428 | ETS2,PPP6C,VCANETS2,PPP6C,VCANETS2,PPP6C,VCAN |
| nilvadipine | 0.0311 | 0.428 | LGALS8,SDCBP,TEP1LGALS8,SDCBP,TEP1LGALS8,SDCBP,TEP1 |
| ESRRA | 0.0315 | 0.431 | ACADM,ACOX1,B4GALT6,CX3CR1,FAM102A,FZD5,NRP2,PIK3R6,Pln,PT |
| fenamic acid | 0.032 | 0.436 | CLU,EMP3,FCGR2A,SPP1,TUBBCLU,EMP3,FCGR2A,SPP1,TUBB |
| ZFTA-RELA | 0.0321 | 0.436 | CASP4,L1CAM,NFKB2,TNIP1CASP4,L1CAM,NFKB2,TNIP1 |
| MAP4K4 | 0.0325 | 0.438 | CBLB,CHPT1,HADHB,ITGA5,NDUFAB1,PDHX,PHYH |
| Esrra | 0.0325 | 0.438 | ACADM,ACOX1,ATP1A1,OXCT1,PFKL,SPP1,THRA |
| lenalidomide | 0.0334 | 0.448 | APTX,CAV2,CTTNBP2,DFFA,FZD5,LRP5,MYB,NEK6,NFIB,PTPRM,SEL1L,T |
| NRIP1 | 0.0338 | 0.449 | ACADM,CCNB2,HADHB,NR2C1,OTUB2 |
| PTP4A1 | 0.0338 | 0.449 | EEF2,FKBP1A,HNRNPH2,NCK1,PIK3CA,PRDX1,VCAN |
| RASSF1 | 0.034 | 0.449 | CASP4,CAV2,CLU,MAP1B,SPP1,TUBB |
| IND S7 | 0.0342 | 0.449 | ETFA,GLOD4,TUBBETFA,GLOD4,TUBBETFA,GLOD4,TUBB |
| ammonium | 0.0342 | 0.449 | ACSF2,ARHGAP33,ATP1A1,BRPF3,CLU,CTSD,FAM98B,PSMB4,UPF1,US |
| CPT1B | 0.0343 | 0.449 | ACADM,CBLB,Cox6c,MAP2K1,MAPK1,NDUFA7,NDUFAB1,NDUFS6,PIK3 |
| AR | 0.0354 | 0.46 | ABCC4,ATP8A1,B4GALT6,BUB1B,CACNA1C,CAV2,CCT3,CDO1,CHPT1,ES |
| ATP5IF1 | 0.0356 | 0.46 | ACADM,ETFA,HADHB,HIBCH,OXCT1 |
| (-)-epicatechin gallate | 0.0366 | 0.46 | SPP1,WWTR1SPP1,WWTR1SPP1,WWTR1SPP1,WWTR1SPP1,WWTR1 |
| quizartinib | 0.0366 | 0.46 | GADD45G,SPP1GADD45G,SPP1GADD45G,SPP1GADD45G,SPP1 |
| BRMS1 | 0.0366 | 0.46 | PLAU,SPP1PLAU,SPP1PLAU,SPP1PLAU,SPP1PLAU,SPP1PLAU,SPP1 |
| KPT-9274 | 0.0366 | 0.46 | FANCF,PAK4FANCF,PAK4FANCF,PAK4FANCF,PAK4FANCF,PAK4 |
| azetidyl-2-carboxylic acid | 0.0366 | 0.46 | CLU,HSPA9CLU,HSPA9CLU,HSPA9CLU,HSPA9CLU,HSPA9CLU,HSPA9 |
| 4-aminophenol | 0.0366 | 0.46 | CLU,SPP1CLU,SPP1CLU,SPP1CLU,SPP1CLU,SPP1CLU,SPP1CLU,SPP1 |
| DLL4 | 0.0366 | 0.46 | FLT1,ITGA5,LYVE1,NFKB2FLT1,ITGA5,LYVE1,NFKB2 |
| guanidinopropionic acid | 0.0366 | 0.46 | ATP8A1,NDUFA7,NDUFB4,NRF1ATP8A1,NDUFA7,NDUFB4,NRF1 |
| IRF7 | 0.0371 | 0.46 | CASP4,DAXX,GBP5,HELZ2,IRF5,NT5C3A,TNFSF13B,TOR1B,USP8 |
| BRD4 | 0.0371 | 0.46 | ABCC4,ANKRA2,CCR1,ITGA5,JARID2,MYB,PLAU,SPP1,TUBB |
| DNMT3A | 0.0372 | 0.46 | CACNA1C,CASP4,CSF1R,IRF5,KDM6B,PIK3CG,PRKCI,SKIL,SPP1,TLR5 |
| xanthine derivative CB002 analo | 0.0374 | 0.46 | APOB,PNKD,RANBP10APOB,PNKD,RANBP10APOB,PNKD,RANBP10 |
| MST1 | 0.0374 | 0.46 | MSR1,SOX4,SPP1MSR1,SOX4,SPP1MSR1,SOX4,SPP1MSR1,SOX4,SPP1 |
| MEL S3 | 0.0374 | 0.46 | ETFA,GLOD4,TUBBETFA,GLOD4,TUBBETFA,GLOD4,TUBB |
| TCF7L2 | 0.0375 | 0.46 | ARAP2,CPD,CTNNA1,DOCK10,FKBP1A,KCNJ3,MAG,MAP7,MOSPD2,MY |
| SORL1 | 0.0383 | 0.466 | EPHA1,EPHA4,FLT1,FZD5,HSP90AB1,ITGA5,LYVE1,NRP2,PLAU |
| Notch | 0.0386 | 0.466 | CSF2RB,FCGR2A,NFAM1,NHSL2,PIK3R6,PLAU,SMAD1,SVIP |
| TNFSF13B | 0.0386 | 0.466 | CCR1,FIGNL1,GCNT2,HLA-DMB,NFKB2,P2RX1,PTPRM,SPP1 |
| NFKB2 | 0.0386 | 0.466 | AKR1B1,FZD5,GBP5,HLA-DMB,MYB,NFKB2,RAB3B,UGT1A6 |
| TRAP1 | 0.0388 | 0.466 | ALDH1L2,EARS2,ECSIT,L1CAM,MTHFD2,NDUFB4 |
| MYBL2 | 0.039 | 0.466 | CCNB2,CLTA,CLU,COPACCNB2,CLTA,CLU,COPACCNB2,CLTA,CLU,COPA |
| SOX10 | 0.039 | 0.466 | GAS7,L1CAM,MAG,SUFUGAS7,L1CAM,MAG,SUFU |
| gentamicin C | 0.0394 | 0.466 | CLU,EMP3,FCGR2A,SPP1,TUBBCLU,EMP3,FCGR2A,SPP1,TUBB |
| SENP1 | 0.0408 | 0.466 | CTSD,GABPB1,NRF1CTSD,GABPB1,NRF1CTSD,GABPB1,NRF1 |
| KCNJ2 | 0.0408 | 0.466 | APBA1,CEP250,FYCO1,GAS7,MAP1B,MYH14,SPTBN2 |
| APOE | 0.0412 | 0.466 | ACADM,ACOX1,APOB,CLU,CSF2RB,CTSD,GAS7,HSP90AB1,ITGA5,MSR1, |
| RNASEH2A | 0.0415 | 0.466 | IRF5,NFKB2,PSMD6,TLR2IRF5,NFKB2,PSMD6,TLR2 |
| thioacetamide | 0.0417 | 0.466 | ABCC4,Anp32a,B4GALT6,CLU,CSNK1D,EMP3,FCGR2A,LYVE1,PCOLCE,P |
| KRAS | 0.042 | 0.466 | ABCC4,AIMP2,CLU,CSF2RB,DUSP6,E2F4,EIF3G,FCGR2A,FGFR1,GALNT1, |
| CREB3L2 | 0.0425 | 0.466 | SEC24C,USO1SEC24C,USO1SEC24C,USO1SEC24C,USO1SEC24C,USO1 |
| HES5 | 0.0425 | 0.466 | FBXW7,MYBFBXW7,MYBFBXW7,MYBFBXW7,MYBFBXW7,MYB |
| CX3CR1 | 0.0431 | 0.466 | BBS7,CEP250,DPCD,DUSP7,FYCO1,MAP1B,MAP7,MSR1,ZXDB |
| STIM1 | 0.0435 | 0.466 | CAMK1D,FLT1,ITGA5,PRDX1,RPS23CAMK1D,FLT1,ITGA5,PRDX1,RPS23 |
| NFATC2 | 0.0439 | 0.466 | CASP4,CX3CR1,DAXX,HELZ2,JARID2,KDM6B,PDZD2,PLD1,TLR2,TNFSF1 |
| VCP | 0.0444 | 0.466 | NFKB2,SEL1L,TNFSF13BNFKB2,SEL1L,TNFSF13B |
| KDM3B | 0.0456 | 0.466 | ADRA2A,ARL6IP1,FKBP1A,HJURP,TSPAN7 |
| HOXA9 | 0.0457 | 0.466 | ETS2,FZD5,MYB,NRF1,RPL37A,SDCBP,SOX4,SPP1,USP1 |
| Ppar | 0.048 | 0.466 | ACADM,ACOX1,HADHBACADM,ACOX1,HADHBACADM,ACOX1,HADHB |
| NOSTRIN | 0.048 | 0.466 | FLT1,ITGA5,PLAUFLT1,ITGA5,PLAUFLT1,ITGA5,PLAUFLT1,ITGA5,PLAU |

| | | | |
|---|---|---|---|
| FOXF1 | 0.048 | 0.466 | CACNA1C,FLT1,ITGA5CACNA1C,FLT1,ITGA5CACNA1C,FLT1,ITGA5 |
| dicarbethoxydihydrocollidine | 0.048 | 0.466 | ABCC4,CCNB2,SPP1ABCC4,CCNB2,SPP1ABCC4,CCNB2,SPP1 |
| PML | 0.0483 | 0.466 | ACADM,ACOX1,DDX60,DNAJC9,HADHB,HLA-DMB,MAP2K1,MAPK1,PRD |
| GON4L | 0.0487 | 0.466 | CCNB2,MYBCCNB2,MYBCCNB2,MYBCCNB2,MYBCCNB2,MYB |
| mGluR | 0.0487 | 0.466 | MAP1B,OPHN1MAP1B,OPHN1MAP1B,OPHN1MAP1B,OPHN1 |
| miR-378a-3p (and other miRNA | 0.0487 | 0.466 | MAPKAPK2,SUFUMAPKAPK2,SUFUMAPKAPK2,SUFUMAPKAPK2,SUFU |
| SPINT2 | 0.0487 | 0.466 | ELF2,PLAUELF2,PLAUELF2,PLAUELF2,PLAUELF2,PLAUELF2,PLAU |
| THZ1 | 0.0488 | 0.466 | CLU,NUPR1,PCOLCE,PDZD2,Prg4,SERPINF1,VCAN |
| beta-estradiol | 0.0491 | 0.466 | ACADM,ACOX1,ANAPC4,Anp32a,APBA1,ARHGAP35,ARID4A,ARPC4,AR |
| CpG oligonucleotide | 0.0497 | 0.466 | CASP4,CCNB2,CCR1,DDX60,ETS2,MSR1,TLR2,TNFSF13,TNFSF13B |
| ZBTB10 | 0.053 | 0.479 | 9930111J21Rik1/Gm12185,CASP4,FLT1,GBP5,Gm5431,LY9,NT5C3A,SE |
| SP2509 | 0.059 | 0.501 | ABCC4,BUB1B,FGFR1,HSPA9,L1CAM,NEK6,NFIB,PAK2,SETDB1,SMAD1, |
| SLC15A4 | 0.0594 | 0.501 | CBLB,CCNB2,DOCK10,GBP5,MSR1,MTHFD2,TUBB |
| IRF6 | 0.0712 | 0.507 | APOBEC3B,PELP1,PLAU,TFAP4APOBEC3B,PELP1,PLAU,TFAP4 |
| 3,5-dihydroxyphenylglycine | 0.0802 | 0.528 | KCNQ1OT1,L1CAM,RPL10,RPL3,RPL37A,RPS23,TPT1 |
| ETV6-RUNX1 | 0.117 | 0.562 | CARD11,EMP3,FGFR1,GPSM3,IRF5,MAP2K1,NCK1,NEK6,P2RX1,PLXNB |
| SERPINE1 | 0.118 | 0.562 | DUSP6,FLT1,NFKB2,PLAU,TLR2DUSP6,FLT1,NFKB2,PLAU,TLR2 |
| tazemetostat | 0.123 | 0.567 | ABCC4,FGFR1,HSPA9,L1CAM,NEK6,NFIB,PAK2,SETDB1,SMAD1,SPP1,V |
| mir-34 | 0.129 | 0.579 | CPD,CSF1R,MYB,SCLT1,SLC4A7CPD,CSF1R,MYB,SCLT1,SLC4A7 |
| CLPP | 0.131 | 0.579 | ACADM,ETFA,HADHB,HSPA9ACADM,ETFA,HADHB,HSPA9 |
| ID3 | 0.141 | 0.579 | CCR1,CSF2RB,ETS2,G6PC3,GADD45G,IRF5,MYB,SOX4 |
| MYCN | 0.157 | 0.605 | ABCC4,ABCF3,CLU,DUSP7,EEF2,HSP90AB1,JARID2,RPL10,RPL3,RPL37A |
| bortezomib | 0.161 | 0.605 | CSF2RB,E2F4,FANCF,MAPK1,NRF1,PSMB4,PSMC4,PSMD12,PSMD2,PSN |
| FBXW7 | 0.174 | 0.605 | CARD11,MCAT,MTHFD2,NFKB2CARD11,MCAT,MTHFD2,NFKB2 |
| indomethacin | 0.199 | 0.615 | ACOX1,CLU,CSF2RB,CTSD,EMP3,FCGR2A,FLT1,SPP1,TUBB,VCAN |
| QKI | 0.211 | 0.62 | CLTA,CSF1R,FYN,MAP1BCLTA,CSF1R,FYN,MAP1B |
| MAPK14 | 0.222 | 0.627 | DUSP6,FLT1,MEF2C,PIN1,PLAU,PRDX1,SPP1,TLR2 |
| HRAS | 0.223 | 0.627 | ACADM,ADRA2A,ARHGAP26,CLU,CTSZ,DUSP6,EIF4G3,GADD45G,GIMA |
| ID2 | 0.229 | 0.627 | CCR1,CSF2RB,ETS2,G6PC3,GADD45G,IRF5,MYB,SOX4 |
| aldosterone | 0.247 | 0.634 | ATP1A1,DAXX,FKBP1A,FLT1,NR3C2,SPP1,TBC1D4 |
| valproic acid | 0.249 | 0.634 | AKR1B1,CCR1,CHPT1,IDS,PSMB2,PSMB4,PSMC4,PSMD12,PSMD2,PSM |
| FOXM1 | 0.271 | 0.653 | BUB1B,CCNB2,FLT1,ITGA5,NFKB2,VCAN |
| dihydrotestosterone | 0.275 | 0.653 | ABCC4,BUB1B,CCT3,CDO1,CLU,FGFR1,GTF2I,MYB,PLAU,POLDIP3,RAB3 |
| estrogen | 0.349 | 0.691 | APOB,CCNB2,CLU,CTSD,FLT1,ITGA5,MTA3,MYB,SPP1 |
| memantine | 0.376 | 0.704 | CACNA1C,EPHA4,HSP90AB1,SYNJ1CACNA1C,EPHA4,HSP90AB1,SYNJ1 |
| DUSP1 | 0.386 | 0.709 | DDX60,DUSP6,HELZ2,NT5C3A,TLR2DDX60,DUSP6,HELZ2,NT5C3A,TLR2 |
| PTGER4 | 0.402 | 0.711 | DAXX,RASSF2,RNF144B,SPP1,TBC1D4 |
| KAT5 | 0.402 | 0.711 | ALDH1L2,MRPL20,OTUB2,PSMC4,PSMD12,PSMD6 |
| PDX1 | 0.402 | 0.711 | AKR1B1,CDO1,DUSP6,FGFR1,MALAT1,SPP1 |
| NFKB1 | 0.415 | 0.719 | AKR1B1,CASP4,DROSHA,HLA-DMB,MSR1,MYB,NFKB2,PLAU,TLR2 |
| lithium chloride | 0.479 | 0.749 | IDS,MYB,PLD1,VCANIDS,MYB,PLD1,VCANIDS,MYB,PLD1,VCAN |
| Growth hormone | 0.51 | 0.762 | AKR1B1,CLU,FZD5,HADHB,SPP1,TBC1D4,Ubb |
| INSR | 0.515 | 0.764 | ACADM,Anp32a,ATP1A1,ETFA,GOT2,HADHB,ITGA5,MRPS21,NDUFB4,N |
| ANGPT2 | 0.54 | 0.772 | CCT7,KDM6B,MAP2K1,NR3C2,PLAU,RBM14 |
| GNAS | 0.549 | 0.773 | ATP1A1,ETS2,FYN,GADD45G,SPP1ATP1A1,ETS2,FYN,GADD45G,SPP1 |
| MLXIPL | 0.554 | 0.774 | RPL10,RPL3,RPL37A,RPS23RPL10,RPL3,RPL37A,RPS23 |
| IgG | 1 | 1 | ACOX1,AFM,ETS2,FCGR2A,FLT1,LY9,SERPINF1,TNIP1 |
| FSH | 1 | 1 | ARHGAP35,ARPC4,CASP4,DUSP3,FGFR1,ITGA5,MAP2K1,PI4K2A,RASSF |
| MRTFA | 1 | 1 | ARPC4,CACNA1C,ITGA5,NFIB,P2RX1 |
| FOXA2 | 1 | 1 | APOB,ASTN1,MAP1B,NFKB2,PDHX,UST |
| SIRT1 | 1 | 1 | ACADM,CCNB2,DDX60,FGFR1,HELZ2,HJURP,HLA-DRB5,MEF2C,NLGN2, |
| CALCA | 1 | 1 | CSF2RB,FLT1,IRF5,RBM14CSF2RB,FLT1,IRF5,RBM14 |

| Master Regulator | Molecule Type | Participating regulators | Depth | p-value of overlap | B-H corrected p-value | Network bias-corrected p-value | Target gene name in analysed list | Causal network | Target-connected regulators |
|---|---|---|---|---|---|---|---|---|---|
| NOP53 | other | 26s Proteasome,ABL1,AKT | 3 | 2.6E-13 | 8.53E-10 | 0.0001 | ABCA9,ABCC4,ABCF3,ACADM,ACOT7,A | 212 (57) | 56 |
| Ep300/Pcaf | complex | 26s Proteasome,ABL1,Akt, | 3 | 3.33E-13 | 8.53E-10 | 0.0001 | ABCF3,ACADM,ACOT7,ACSF2,AIMP2,AK | 219 (66) | 64 |
| SGI 1776 | chemical drug | Akt,AKT1,AR,ATF4,BAD,BR | 3 | 3.51E-13 | 8.53E-10 | 0.0001 | ABCA9,ABCC4,ABCF3,ABHD2,ACADM,A | 199 (47) | 42 |
| ARVib-7 | chemical reagent | AHR,AR,ARVib-7,ATXN3,CF | 3 | 5.49E-13 | 8.53E-10 | 0.0001 | ABCA9,ABCC4,ABCF3,ACOX1,ACSF2,AG | 184 (36) | 36 |
| STUB1 | enzyme | AHR,AR,ATXN3,CFTR,EIF4E | 2 | 1E-12 | 8.53E-10 | 0.0001 | ABCA9,ABCC4,ABCF3,ACOX1,ACSF2,AG | 182 (35) | 35 |
| TT-00420 | chemical drug | AKT1,AURKA,AURKB,BRCA | 3 | 1.01E-12 | 8.53E-10 | 0.0001 | ABCC4,ABCF3,ACOX1,ACSF2,AIMP2,AK | 128 (20) | 17 |
| XMD16-144 | chemical drug | AKT1,AURKA,AURKB,BRCA | 3 | 1.01E-12 | 8.53E-10 | 0.0001 | ABCC4,ABCF3,ACOX1,ACSF2,AIMP2,AK | 128 (20) | 17 |
| SNS 314 | chemical drug | AKT1,AURKA,AURKB,BRCA | 3 | 1.01E-12 | 8.53E-10 | 0.0001 | ABCC4,ABCF3,ACOX1,ACSF2,AIMP2,AK | 128 (20) | 17 |
| AMG 900 | chemical drug | AKT1,AMG 900,AURKA,AU | 3 | 1.06E-12 | 8.53E-10 | 0.0001 | ABCC4,ABCF3,ACOX1,ACSF2,AIMP2,AK | 128 (20) | 17 |
| TTP607 | chemical drug | AKT1,AURKA,AURKB,BRCA | 3 | 1.06E-12 | 8.53E-10 | 0.0001 | ABCC4,ABCF3,ACOX1,ACSF2,AIMP2,AK | 128 (20) | 17 |
| rocuronium | chemical drug | Akt,AKT1,CHRM2,CHRNA3 | 3 | 1.79E-12 | 1.31E-09 | 0.0001 | ABCC4,ABCF3,ACADM,ACOT7,ACSF2,AK | 135 (22) | 16 |
| MAPKAPK5 | kinase | ABL1,AKT1,AR,BHLHE40,C | 3 | 2.02E-12 | 1.36E-09 | 0.0002 | ABCA9,ABCF3,ACADM,ACOT7,ACSF2,A | 211 (61) | 56 |
| TFDP1 | transcription regulator | 26s Proteasome,AKT1,AR, | 3 | 3.38E-12 | 1.97E-09 | 0.0001 | ABCF3,ACADM,ACSF2,AGGF1,AK1,AKR1 | 229 (73) | 72 |
| SOX2 | transcription regulator | 26s Proteasome,ABL1,Akt, | 3 | 3.42E-12 | 1.97E-09 | 0.0002 | ABCF3,ACADM,ACOT7,ACSF2,ADRA2A, | 221 (52) | 51 |
| harmine | chemical - endogenous non-mammalian | 2-amino-1-methyl-6-pheny | 3 | 4.93E-12 | 2.64E-09 | 0.0001 | ABCF3,ACADM,ACOX1,ACSF2,ADRA2A, | 227 (64) | 62 |
| CYBB | enzyme | Akt,AKT1,CHUK,CYBB,EGF | 2 | 5.51E-12 | 2.77E-09 | 0.0001 | ACADM,ACOT7,ACSF2,AIMP2,AK1,AKR1 | 129 (20) | 19 |
| triflusal | chemical drug | ATF1,BTRC,CDX1,Creb,EGF | 3 | 6.2E-12 | 2.82E-09 | 0.0001 | ABCC4,ABCF3,ACADM,ACOT7,ACSF2,A | 155 (34) | 31 |
| RPL23A | other | 26s Proteasome,ABL1,Akt, | 3 | 6.31E-12 | 2.82E-09 | 0.0001 | ABCA9,ABCF3,ACADM,ACSF2,AGGF1,A | 206 (49) | 48 |
| CEBPA | transcription regulator | aflatoxin B1,APP,AR,Calcin | 3 | 7.71E-12 | 3.26E-09 | 0.0001 | 2810025M15Rik,ABCC4,ABCF3,ACOT7, | 230 (87) | 81 |
| mechlorethamine | chemical drug | 26s Proteasome,ABL1,Akt, | 3 | 8.34E-12 | 3.35E-09 | 0.0002 | ABCF3,ACADM,ACOT7,ACSF2,ADRA2A, | 217 (65) | 62 |
| CTDSP1 | phosphatase | ATF4,BMI1,BRCA1,CASP8, | 3 | 9.05E-12 | 3.47E-09 | 0.0002 | ABCA9,ABCC4,ABCF3,ABHD2,ACADM,A | 184 (34) | 33 |
| ZEB | group | Akt,AKT1,BBC3,CCND1,CH | 3 | 1.01E-11 | 3.58E-09 | 0.0001 | ABCF3,ACADM,ACSF2,AIMP2,AK1,AKR1 | 139 (19) | 18 |
| Jmy-p300 | complex | 26s Proteasome,ABL1,Akt, | 3 | 1.13E-11 | 3.58E-09 | 0.0001 | ABCF3,ACADM,ACOT7,ACSF2,AIMP2,AK | 214 (64) | 62 |
| PLK1 | kinase | 26s Proteasome,Akt,CDC2 | 2 | 1.17E-11 | 3.58E-09 | 0.0001 | ABCA9,ABCC4,ABCF3,ABHD2,ACOX1,A | 167 (25) | 25 |
| ACTL6A | other | ACTL6A,AR,ATF4,BRCA1,C | 3 | 1.2E-11 | 3.58E-09 | 0.0002 | ABCA9,ABCC4,ABCF3,ABHD2,ACADM,A | 202 (32) | 32 |
| DNMT3A | enzyme | 26s Proteasome,Akt,Alp,B | 3 | 1.22E-11 | 3.58E-09 | 0.0002 | ABCA9,ABCF3,ACADM,ACOT7,ACSF2,A | 218 (62) | 60 |
| CHCHD4 | enzyme | AHR,Akt,AKT1,APLN,ATF4, | 3 | 1.22E-11 | 3.58E-09 | 0.0001 | ABCA9,ABCF3,ACADM,ACOT7,ACSF2,A | 228 (64) | 62 |
| mustard gas | chemical toxicant | 26s Proteasome,ABL1,Akt, | 3 | 1.27E-11 | 3.58E-09 | 0.0002 | ABCA9,ABCF3,ACADM,ACOT7,ACSF2,A | 221 (67) | 65 |
| ACP-319 | chemical drug | ACP-319,Akt,AKT1,Creb,EL | 3 | 1.29E-11 | 3.58E-09 | 0.0001 | 9930111J21Rik1/Gm12185,ABCA9,ABC | 158 (36) | 34 |
| PFDN5 | transcription regulator | Akt,ATF4,BRCA1,CASP8,CC | 3 | 1.53E-11 | 4.1E-09 | 0.0002 | ABCA9,ABCC4,ABCF3,ABHD2,ACADM,A | 180 (29) | 28 |
| TRIM29 | transcription regulator | 26s Proteasome,ABL1,Akt, | 3 | 1.59E-11 | 4.11E-09 | 0.0001 | ABCF3,ACADM,ACOT7,ACSF2,AGGF1,A | 208 (59) | 55 |
| CUL7 | enzyme | 26s Proteasome,ABL1,AKT | 3 | 1.7E-11 | 4.28E-09 | 0.0001 | ABCA9,ABCF3,ACADM,ACOT7,ACSF2,A | 208 (61) | 58 |
| NPM1 | transcription regulator | 26s Proteasome,ABL1,AHR | 3 | 1.92E-11 | 4.69E-09 | 0.0003 | ABCF3,ACADM,ACOT7,ACSF2,ADRA2A, | 226 (66) | 63 |
| ZEB1 | transcription regulator | BAX,BBC3,CCND1,CHEK1,N | 2 | 2.08E-11 | 4.86E-09 | 0.0001 | ABCF3,ACADM,ACSF2,AIMP2,AK1,AKR1 | 128 (12) | 12 |
| TRIM31 | enzyme | 26s Proteasome,ABL1,Akt, | 3 | 2.19E-11 | 4.86E-09 | 0.0001 | ABCF3,ACADM,ACOT7,ACOX1,ACSF2,A | 202 (53) | 53 |
| PI3K (family) | group | Akt,AKT1,Creb,ELAVL1,ERK | 2 | 2.27E-11 | 4.86E-09 | 0.0001 | 9930111J21Rik1/Gm12185,ABCC4,ABC | 156 (32) | 32 |
| FOXD1 | transcription regulator | 26s Proteasome,AHR,Akt, | 3 | 2.27E-11 | 4.86E-09 | 0.0001 | ABCA9,ABCF3,ACOX1,ACSF2,AGGF1,AK | 242 (69) | 67 |
| OXSR1 | kinase | 26s Proteasome,ABL1,AR, | 3 | 2.3E-11 | 4.86E-09 | 0.0004 | ABCF3,ACADM,ACOT7,ACSF2,AK1,AKR1 | 219 (56) | 55 |
| MTORC2 | complex | Akt,AKT1,AKT2,EIF4B,Gsk3 | 2 | 2.36E-11 | 4.86E-09 | 0.0001 | ACSF2,AIMP2,AK1,AKR1B1,ANAPC4,AR | 110 (14) | 13 |
| NCOA4 | transcription regulator | AHR,Akt,AKT1,ALDH1A1,A | 3 | 2.46E-11 | 4.93E-09 | 0.0003 | ABCA9,ABCC4,ABHD2,ACADM,ACOT7,A | 206 (53) | 53 |
| APEX1 | enzyme | 26s Proteasome,ABL1,AHR | 3 | 2.75E-11 | 5.38E-09 | 0.0004 | ABCA9,ABCF3,ACOT7,ACSF2,ACTR1B,A | 226 (72) | 68 |
| TSC22D1 | transcription regulator | 26s Proteasome,ABL1,AKT | 3 | 2.97E-11 | 5.68E-09 | 0.0001 | ABCF3,ACADM,ACOT7,ACSF2,AGGF1,A | 216 (57) | 56 |
| FOXJ1 | transcription regulator | 26s Proteasome,AHR,AKT1 | 3 | 3.34E-11 | 6.13E-09 | 0.0001 | ABCA9,ABCF3,ACOX1,ACSF2,AGGF1,AK | 240 (69) | 66 |
| FRAX120 | chemical drug | ABL1,Akt,CASP1,CASP8,Cr | 3 | 3.47E-11 | 6.13E-09 | 0.0002 | ABCC4,ABCF3,ABHD2,ACOT7,ACSF2,AD | 162 (38) | 34 |
| FRAX355 | chemical drug | ABL1,Akt,CASP1,CASP8,Cr | 3 | 3.47E-11 | 6.13E-09 | 0.0002 | ABCC4,ABCF3,ABHD2,ACOT7,ACSF2,AD | 162 (38) | 34 |
| RPL11 | other | MDM2,MYC,RPL11,TP53 | 2 | 3.63E-11 | 6.13E-09 | 0.0001 | ABCC4,ABCF3,ACADM,ACSF2,AIMP2,AK | 111 (4) | 3 |
| miR-30c-5p (and other miR | mature microRNA | 26s Proteasome,ABL1,Akt, | 3 | 3.66E-11 | 6.13E-09 | 0.0003 | ABCF3,ACADM,ACOT7,ACSF2,ADRA2A, | 214 (55) | 54 |
| stigmatellin | chemical reagent | 26s Proteasome,Akt,AKT1, | 3 | 3.71E-11 | 6.13E-09 | 0.0001 | ABCF3,ACADM,ACSF2,AGGF1,AIMP2,AK | 229 (74) | 73 |
| butyric acid | chemical - endogenous mammalian | AR,bezafibrate,bortezomib | 2 | 3.8E-11 | 6.13E-09 | 0.0001 | ABCC4,ABCF3,ABHD2,ACOX1,ACSF2,AD | 167 (33) | 33 |
| Arf | group | 26s Proteasome,ABL1,AKT | 3 | 3.82E-11 | 6.13E-09 | 0.0005 | ABCC4,ABCF3,ACADM,ACSF2,AK1,AKR1 | 215 (64) | 60 |
| SP100 | transcription regulator | 26s Proteasome,Akt,AKT1, | 3 | 3.9E-11 | 6.15E-09 | 0.0007 | ABCF3,ACADM,ACOT7,ACSF2,ADRA2A, | 225 (60) | 59 |
| MUS81 | enzyme | 26s Proteasome,ABL1,AKT | 3 | 4.2E-11 | 6.49E-09 | 0.0001 | ABCF3,ACADM,ACOT7,ACSF2,AGGF1,A | 196 (59) | 57 |
| KCTD12 | ion channel | AKT1,APP,AR,AURKA,BRCA | 3 | 4.55E-11 | 6.79E-09 | 0.0001 | ABCA9,ABCC4,ABCF3,ACOX1,ACSF2,AG | 163 (41) | 38 |
| SMARCD1 | transcription regulator | 26s Proteasome,ABL1,AKT | 3 | 4.56E-11 | 6.79E-09 | 0.0005 | ABCF3,ABHD2,ACADM,ACOT7,ACSF2,A | 220 (61) | 60 |
| PARK7 | enzyme | Akt,AKT1,ERK1/2,GSK3B,H | 2 | 4.83E-11 | 7.02E-09 | 0.0001 | ABCA9,ABCC4,ACADM,ACOX1,ACSF2,A | 114 (11) | 10 |
| oltipraz | chemical drug | ABL1,aflatoxin B1,AKT1,ari | 3 | 4.89E-11 | 7.02E-09 | 0.0003 | ABCF3,ACADM,ACSF2,AGGF1,AK1,AKR1 | 202 (71) | 65 |
| LEF1 | transcription regulator | AR,ATF4,BRCA1,CCND1,CD | 3 | 5E-11 | 7.05E-09 | 0.0002 | ABCA9,ABCC4,ABCF3,ABHD2,ACADM,A | 210 (48) | 47 |
| pentachlorophenol | chemical toxicant | 2-amino-1-methyl-6-pheny | 3 | 5.32E-11 | 7.26E-09 | 0.0003 | ABCA9,ABHD2,ACADM,ACOT7,ACOX1,A | 186 (45) | 43 |
| APLP2 | other | 26s Proteasome,ABL1,Akt, | 3 | 5.33E-11 | 7.26E-09 | 0.0002 | ABCA9,ABCF3,ACADM,ACOT7,ACSF2,A | 206 (56) | 54 |
| DACH1 | transcription regulator | 26s Proteasome,Akt,AKT1, | 3 | 5.52E-11 | 7.39E-09 | 0.0001 | ABCA9,ABCC4,ABCF3,ACADM,ACOT7,A | 217 (76) | 75 |
| TWIST | group | 26s Proteasome,ABL1,AKT | 3 | 6.12E-11 | 8.06E-09 | 0.0001 | ABCA9,ABCF3,ACADM,ACOT7,ACOX1,A | 194 (60) | 59 |
| ING1 | transcription regulator | 26s Proteasome,ABL1,Akt, | 3 | 6.52E-11 | 8.45E-09 | 0.0002 | ABCA9,ABCF3,ACADM,ACOT7,ACSF2,A | 211 (60) | 59 |
| HEY1 | transcription regulator | 26s Proteasome,ABL1,Akt, | 3 | 7.03E-11 | 8.97E-09 | 0.0004 | ABCC4,ABCF3,ACADM,ACOT7,ACSF2,A | 212 (58) | 57 |
| COL1A1 | other | Akt,AKT1,Calcineurin prote | 3 | 7.32E-11 | 9.09E-09 | 0.0001 | ABCA9,ABHD2,ACOT7,ACSF2,AGGF1,A | 184 (59) | 55 |
| HUWE1 | transcription regulator | AR,EGFR,HUWE1,KRAS,MC | 2 | 7.35E-11 | 9.09E-09 | 0.0001 | ABCF3,ACOT7,ACSF2,AIMP2,AK1,AKR1E | 125 (8) | 7 |
| IFI16 | transcription regulator | 26s Proteasome,ABL1,AHR | 3 | 7.47E-11 | 9.1E-09 | 0.0003 | ABCF3,ACADM,ACOT7,ACOX1,ACSF2,A | 217 (70) | 68 |
| WDR5 | transcription regulator | 26s Proteasome,ABL1,Akt, | 3 | 7.68E-11 | 9.21E-09 | 0.0006 | ABCF3,ACADM,ACOT7,ACSF2,ADRA2A, | 210 (56) | 55 |
| lovastatin | chemical drug | 26s Proteasome,Akt,AR,CA | 2 | 7.8E-11 | 9.22E-09 | 0.0002 | ABCC4,ABCF3,ACOT7,ACOX1,ACSF2,AD | 155 (24) | 24 |
| TRRAP | transcription regulator | APP,AR,ATF4,BRCA1,CASP | 3 | 8.45E-11 | 9.65E-09 | 0.0004 | ABCA9,ABCC4,ABCF3,ABHD2,ACADM,A | 201 (47) | 45 |
| 5,5'-dithiobis(2-nitrobenzoi | chemical reagent | 5,5'-dithiobis(2-nitrobenzo | 3 | 8.47E-11 | 9.65E-09 | 0.0001 | ABCA9,ABCC4,ACOT7,ACOX1,ACSF2,AD | 132 (28) | 26 |
| MELK-8a | chemical drug | Akt,AKT1,APP,AR,AURKA,A | 3 | 8.53E-11 | 9.65E-09 | 0.0003 | ABCA9,ABCC4,ABCF3,ACOX1,ACSF2,AG | 166 (48) | 43 |
| CCDC6 | other | Akt,AKT1,AR,CASP8,CCDC6 | 3 | 8.85E-11 | 9.88E-09 | 0.0003 | ABCA9,ABCC4,ABHD2,ACADM,ACOT7,A | 180 (51) | 50 |
| warfarin | chemical drug | Akt,AKT1,Alp,AR,AXL,CASP | 3 | 9.37E-11 | 1.03E-08 | 0.0001 | ABCA9,ABCC4,ABCF3,ACADM,ACOT7,A | 164 (49) | 43 |
| miR-221-3p (and other miR | mature microRNA | Akt,AKT1,AR,BBC3,CASP8, | 3 | 9.71E-11 | 1.05E-08 | 0.0003 | ABCA9,ABCC4,ABHD2,ACADM,ACOT7,A | 180 (52) | 52 |
| trichosanthin | chemical drug | 26s Proteasome,ABL1,AKT | 3 | 9.84E-11 | 1.05E-08 | 0.0002 | ABCA9,ABCF3,ACADM,ACOT7,ACSF2,A | 207 (67) | 62 |
| ZMIZ1 | transcription regulator | Akt,AKT1,AR,ATF4,BRCA1, | 3 | 1.01E-10 | 1.06E-08 | 0.0005 | ABCA9,ABCC4,ABCF3,ACADM,ACOT7,A | 218 (50) | 49 |
| pidnarulex | chemical drug | 26s Proteasome,ABL1,Akt, | 3 | 1.02E-10 | 1.06E-08 | 0.0002 | ABCA9,ABCF3,ACADM,ACOT7,ACSF2,A | 205 (61) | 58 |
| HOXA2 | transcription regulator | 26s Proteasome,ABL1,Akt, | 3 | 1.04E-10 | 1.07E-08 | 0.0003 | ABCF3,ACADM,ACOT7,ACSF2,ADRA2A, | 212 (56) | 54 |
| APH-1 | group | Akt,AKT1,APH-1,APH1A,AP | 3 | 1.11E-10 | 1.11E-08 | 0.0001 | 2700097O09Rik,ACADM,ACSF2,AIMP2, | 102 (16) | 14 |
| cobalt | chemical toxicant | 26s Proteasome,ABL1,AHR | 3 | 1.14E-10 | 1.11E-08 | 0.0007 | ABCA9,ABCF3,ACADM,ACSF2,ADRA2A, | 211 (51) | 48 |
| TRIM28 | transcription regulator | E2F1,E2F3,MYC,RB1,RELA, | 2 | 1.15E-10 | 1.11E-08 | 0.0001 | ABCC4,ABCF3,ACSF2,AIMP2,AK1,AKR1E | 123 (8) | 8 |
| ELL | transcription regulator | 26s Proteasome,ABL1,AKT | 3 | 1.16E-10 | 1.11E-08 | 0.0006 | ABCA9,ABCF3,ACADM,ACOT7,ACSF2,A | 215 (63) | 61 |
| SMYD2 | enzyme | 26s Proteasome,ABL1,AKT | 3 | 1.16E-10 | 1.11E-08 | 0.0002 | ABCF3,ACADM,ACOT7,ACSF2,AGGF1,A | 200 (54) | 52 |
| KAT5 | transcription regulator | EOMES,KAT5,MAPK8,MAP | 2 | 1.17E-10 | 1.11E-08 | 0.0001 | ABCC4,ABCF3,ACSF2,AIMP2,AK1,AKR1E | 128 (8) | 8 |
| FBXL14 | enzyme | Akt,ATF4,BMI1,BRCA1,CAS | 3 | 1.18E-10 | 1.11E-08 | 0.0002 | ABCA9,ABCC4,ABCF3,ACADM,ACOX1,A | 184 (36) | 34 |
| selpercatinib | chemical drug | AKT1,ATF4,Creb,EGFR,EIF4 | 3 | 1.21E-10 | 1.12E-08 | 0.0003 | ABCA9,ABCC4,ABCF3,ABHD2,ACADM,A | 173 (44) | 41 |
| RUNX1-RUNX1T1 | fusion gene/product | CBFB,CEBPA,FOXO3,GATA | 3 | 1.22E-10 | 1.12E-08 | 0.0001 | ABCC4,ABCF3,ACOX1,ACSF2,AGGF1,AI | 140 (17) | 17 |
| CSF2 receptor | complex | AKT1,ATF4,BCR-ABL1,BEC | 3 | 1.27E-10 | 1.15E-08 | 0.0002 | ABCA9,ABCC4,ABCF3,ACADM,ACOT7,A | 191 (64) | 59 |
| PAXIP1 | other | 26s Proteasome,ABL1,Akt, | 3 | 1.27E-10 | 1.15E-08 | 0.0003 | ABCF3,ACADM,ACOT7,ACSF2,ADRA2A, | 205 (54) | 53 |
| BAIAP2 | kinase | Akt,AKT1,BAIAP2,CASP1,C | 3 | 1.32E-10 | 1.18E-08 | 0.0002 | ABCA9,ABCC4,ABCF3,ABHD2,ACADM,A | 153 (37) | 34 |
| NSD2 | enzyme | 26s Proteasome,ABL1,Akt, | 3 | 1.38E-10 | 1.21E-08 | 0.0011 | ABCA9,ABCF3,ACADM,ACOT7,ACSF2,A | 224 (64) | 63 |
| Smad2/3/4 | group | Akt,AKT1,Ap1,AR,BRCA1,B | 3 | 1.39E-10 | 1.21E-08 | 0.0005 | ABCC4,ABCF3,ABHD2,ACOT7,ACTR1B,A | 210 (54) | 52 |
| RPS6KB1 | kinase | Akt,Creb,CREB1,E2f,EEF2K, | 2 | 1.42E-10 | 1.23E-08 | 0.0001 | ABCC4,ACSF2,AIMP2,AK1,AKR1B1,ANA | 125 (19) | 19 |
| YAP/TAZ | group | Akt,AKT1,AR,CASP8,CCNE1 | 3 | 1.45E-10 | 1.24E-08 | 0.0002 | ABCA9,ABCC4,ABHD2,ACADM,ACOT7,A | 182 (56) | 55 |
| TOPBP1 | other | 26s Proteasome,ABL1,Akt, | 3 | 1.49E-10 | 1.26E-08 | 0.0004 | ABCA9,ABCF3,ACADM,ACSF2,AGGF1,A | 214 (69) | 65 |
| G3BP1 | enzyme | 26s Proteasome,AKT1,AR, | 3 | 1.51E-10 | 1.26E-08 | 0.0004 | 2810025M15Rik,ABCA9,ABCF3,ACADM | 208 (67) | 64 |
| DNMT3B | enzyme | Akt,AKT1,AR,CASP8,CCNE1 | 3 | 1.52E-10 | 1.26E-08 | 0.0004 | ABCA9,ABCC4,ABHD2,ACADM,ACOT7,A | 181 (51) | 50 |
| YWHAH | transcription regulator | 26s Proteasome,Akt,AKT1, | 3 | 1.57E-10 | 1.29E-08 | 0.0001 | ABCA9,ABCC4,ABHD2,ACADM,ACOT7,A | 154 (50) | 47 |
| USP35 | peptidase | AHR,Akt,AKT1,APP,AR,Calc | 3 | 1.6E-10 | 0.000000013 | 0.0003 | ABCA9,ABCC4,ABHD2,ACADM,ACOT7,A | 189 (45) | 44 |
